# Chromosome-level *de novo* genome assembly of *Telopea speciosissima* (New South Wales waratah) using long-reads, linked-reads and Hi-C

**DOI:** 10.1101/2021.06.02.444084

**Authors:** Stephanie H Chen, Maurizio Rossetto, Marlien van der Merwe, Patricia Lu-Irving, Jia-Yee S Yap, Hervé Sauquet, Greg Bourke, Timothy G Amos, Jason G Bragg, Richard J Edwards

## Abstract

*Telopea speciosissima,* the New South Wales waratah, is an Australian endemic woody shrub in the family Proteaceae. Waratahs have great potential as a model clade to better understand processes of speciation, introgression and adaptation, and are significant from a horticultural perspective. Here, we report the first chromosome-level genome for *T. speciosissima*. Combining Oxford Nanopore long-reads, 10x Genomics Chromium linked-reads and Hi-C data, the assembly spans 823 Mb (scaffold N50 of 69.0 Mb) with 97.8 % of Embryophyta BUSCOs complete. We present a new method in Diploidocus (https://github.com/slimsuite/diploidocus) for classifying, curating and QC-filtering scaffolds, which combines read depths, k-mer frequencies and BUSCO predictions. We also present a new tool, DepthSizer (https://github.com/slimsuite/depthsizer), for genome size estimation from the read depth of single copy orthologues and estimate the genome size to be approximately 900 Mb. The largest 11 scaffolds contained 94.1 % of the assembly, conforming to the expected number of chromosomes (2*n* = 22). Genome annotation predicted 40,158 protein-coding genes, 351 rRNAs and 728 tRNAs. We investigated *CYCLOIDEA* (*CYC*) genes, which have a role in determination of floral symmetry, and confirm the presence of two copies in the genome. Read depth analysis of 180 ‘Duplicated’ BUSCO genes suggest almost all are real duplications, increasing confidence in protein family analysis using annotated protein-coding genes, and highlighting a possible need to revise the BUSCO set for this lineage. The chromosome-level *T. speciosissima* reference genome (Tspe_v1) provides an important new genomic resource of Proteaceae to support the conservation of flora in Australia and further afield.

## INTRODUCTION

*Telopea* R.Br. is an eastern Australian genus of five species of large, long-lived shrubs in the flowering plant family Proteaceae. The New South Wales waratah, *Telopea speciosissima* (Sm.) R.Br., is a striking and iconic member of the Australian flora, characterised by large, terminal inflorescences of red flowers (Figure 1). It has been the state floral emblem of New South Wales since 1962 and was one of the first Australian plant species collected for cultivation in Europe (Nixon, 1987). The species is endemic to the state of New South Wales, occurring on sandstone ridges in the Sydney region. Previous studies have investigated variation among *Telopea* populations by phenetic analysis of morphology (Crisp & Weston, 1993) and evolutionary relationships using cladistics (Weston & Crisp, 1994). Population structure and patterns of divergence and introgression between *T. speciosissima* populations have been characterised using several loci (Rossetto et al., 2011). Further, microsatellite data and modelling suggest a history of allopatric speciation followed by secondary contact and hybridization among *Telopea* species (Rossetto et al., 2012). These studies point to the great potential of *Telopea* as a model clade for understanding processes of divergence, environmental adaptation and speciation. Our understanding of these processes can be greatly enhanced by a genome-wide perspective, enabled by a reference genome (Ellegren et al., 2012; Hoban et al., 2016; Lewin et al., 2018; Radwan & Babik, 2012; Seehausen et al., 2014).

**Figure 1.**
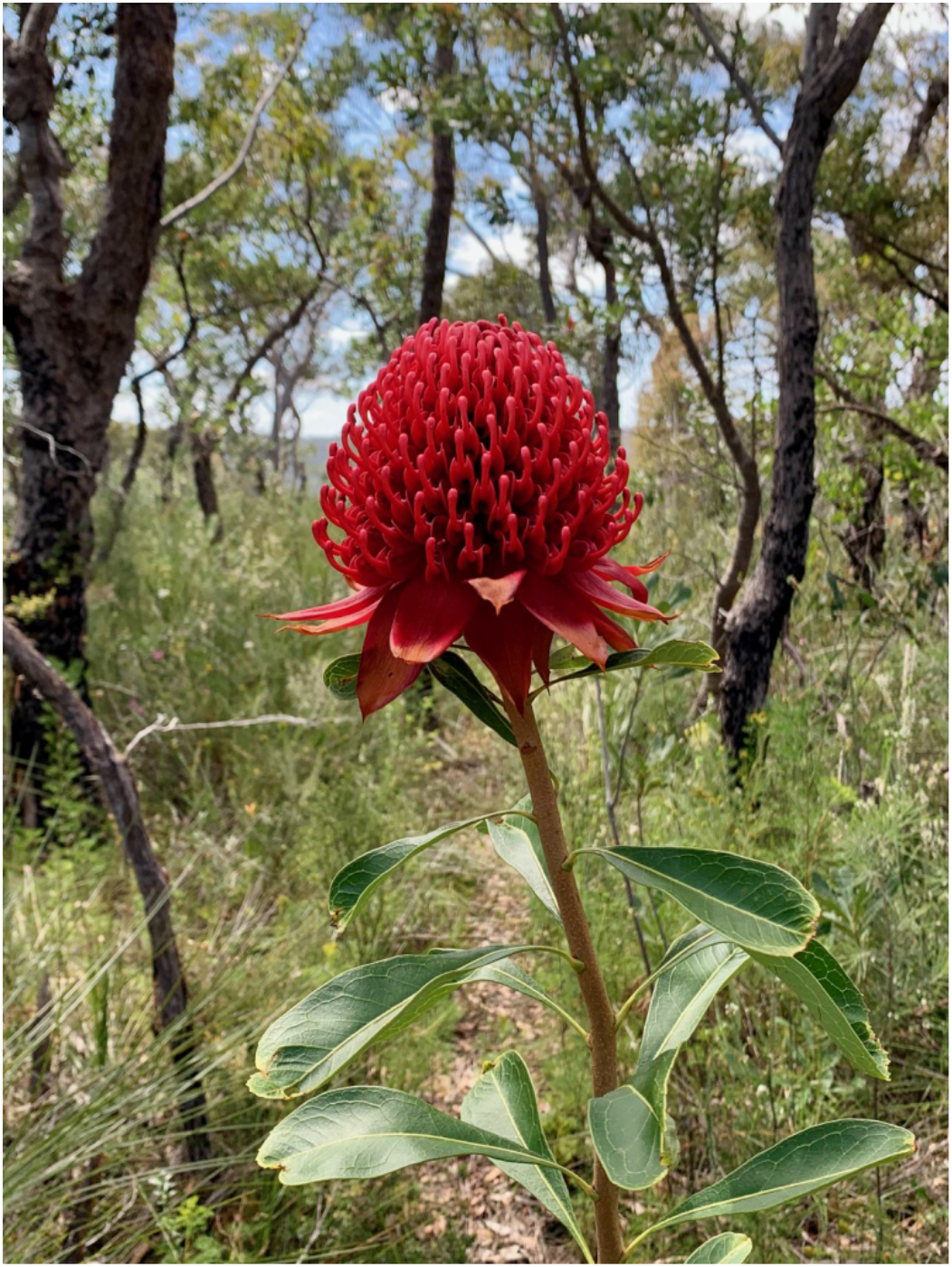
New South Wales waratah (*Telopea speciosissima*). Photo taken by SH Chen.

Genome sequencing efforts have traditionally focused on model species, crops and their wild relatives, resulting in a highly uneven species distribution of reference genomes across the plant tree of life (Royal Botanic Gardens, Kew, 2017). Despite Proteaceae occurring across several continents and encompassing 81 genera and ca. 1700 species (Mast et al., 2008; Weston, 2006), the only publicly available reference genome in the family is a widely-grown cultivar of the most economically important crop in the family, *Macadamia integrifolia* (macadamia nut) HAES 74 (Nock et al., 2016, 2020). Waratahs are significant to the horticultural and cut flower industries, with blooms cultivated for the domestic and international markets (Offord et al., 1987; Worrall & Gollnow, 2013). A reference genome will accelerate efforts in breeding for traits such as resistance to pests and diseases (e.g. *Phytophthora* and *Cylindrocapon destructans* infection; Summerell, 1997; Summerell et al., 1990) as well as desirable floral characteristics (Offord, 2003, 2006).

Technological advances in sequencing and decreasing costs will facilitate the generation of more flowering plant reference genomes, including within the Proteaceae family, and advance research into links between the evolution of genomes and traits that exhibit exceptional diversity, such as floral morphology (Soltis & Soltis, 2014; Zheng et al., 2021). *CYCLOIDEA* (*CYC*) genes belong to the TPC transcription factor gene family, and are known to have an essential role in determining floral symmetry and inflorescence architecture in many angiosperm lineages (Busch & Zachgo, 2009; Fambrini & Pugliesi, 2017; Horn et al., 2015; Luo et al., 1996); studies have characterised recurrent duplications of members of the *CYC2* clade, especially in eudicots (Howarth & Donoghue, 2006), including Fabales (Citerne et al., 2003; Feng et al., 2006), Asterales (Chapman et al., 2008), and Lamiales (Yang et al., 2015; Zhong & Kellogg, 2015). In Proteaceae, a single duplication of *CYC*-like genes occurred prior to diversification and two genes, *ProtCYC1* and *ProtCYC2*, have been characterised (Citerne et al., 2017). In particular, *Grevillea juniperina* has been studied in detail (Damerval et al., 2019) and the existence of both *ProtCYC1* and *ProtCYC2* in *Telopea mongaensis* has been supported by phylogenetic analysis (Citerne et al., 2017). However, *CYC* copy number has not been established in *T. speciosissima*.

Here, we provide a high quality chromosome-level *de novo* assembly of the *Telopea speciosissima* genome, using Oxford Nanopore long-reads, 10x Genomics Chromium linked-reads and Hi-C proximity ligation scaffolding, which will serve as an important platform for evolutionary genomics and the conservation of the Australian flora. We present an analysis of *CYC* genes in the genome to contribute to the understanding of floral evolution in the Proteaceae family.

## MATERIALS AND METHODS

### Sampling and DNA extraction

Young leaves (approx. 8 g) were sampled from the reference genome individual (NCBI BioSample SAMN18238110) where it grows naturally along the Tomah Spur Fire Trail (-33.53° S, 150.42° E) on land belonging to the Blue Mountains Botanic Garden, Mount Tomah in New South Wales, Australia. Leaves were immediately frozen in liquid nitrogen and stored at -80° C prior to extraction.

High-molecular-weight (HMW) genomic DNA (gDNA) was obtained using a sorbitol pre-wash step prior to a CTAB extraction adapted from Inglis et al. (2018). The gDNA was then purified with AMPure XP beads (Beckman Coulter, Brea, CA, USA) using a protocol based on Schalamun et al. (2019) – details available on protocols.io (Lu-Irving & Rutherford, 2021). The quality of the DNA was assessed using Qubit, NanoDrop and TapeStation 2200 System (Agilent, Santa Clara, CA, USA).

### ONT PromethION sequencing

We performed an in-house sequencing test on the MinION (MinION, <u>RRID:SCR_017985</u>) using a FLO-MINSP6 (R9.4.1) flow cell with a library prepared with the ligation kit (SQK-LSK109). The remaining purified genomic DNA was sent to the Australian Genome Research Facility (AGRF) where size selection was performed to remove small DNA fragments using the BluePippin High Pass Plus Cassette on the BluePippin (Sage Science, Beverly, MA, USA). Briefly, 10 µg of DNA was split into 4 aliquots (2.5 µg) and diluted to 60 µL in TE buffer. Then, 20 µL of RT equilibrated loading buffer was added to each aliquot and mixed by pipetting. Samples were loaded on the cassette by removing 80 µL of buffer from each well and adding 80 µL of sample or external marker. The cassette was run with the 15 kb High Pass Plus Marker U1 cassette definition. Size selected fractions (approximately 80 µL) were collected from the elution module following a 30 min electrophoresis run. The library was prepared with the ligation sequencing kit (SQK-LSK109). The sequencing was performed using MinKNOW v.19.12.2 (MinION) and v12.12.8 (PromethION) and MinKNOW Core v3.6.7 (in-house test), v3.6.8 (AGRF MinION) and v3.6.7 (AGRF PromethION). A pilot run was first performed on the MinION using the FLO-MIN106 (R9.4.1) flow cell followed by two FLO-PRO002 flow cells (R9.4) on the PromethION (PromethION, <u>RRID:SCR_017987</u>)

Basecalling was performed after sequencing with GPU-enabled Guppy v3.4.4 using the high-accuracy flip-flop models, resulting in 54x coverage. The output from all ONT basecalling was pooled for adapter removal using Porechop (Porechop, <u>RRID:SCR_016967</u>) v.0.2.4 (Wick et al., 2017) and quality filtering (removal of reads less than 500 bp in length and Q lower than 7) with NanoFilt (NanoFilt, <u>RRID:SCR_016966</u>) v2.6.0 (De Coster et al., 2018) followed by assessment using FastQC (FastQC, <u>RRID:SCR_014583</u>) v0.11.8 (Andrews, 2010).

### 10x Genomics Chromium sequencing

High-molecular-weight gDNA was sent to AGRF for 10x Genomics Chromium sequencing. Size selection was performed to remove DNA fragments <40 kb using the BluePippin 0.75 % Agarose Gel Cassette, Dye Free on the BluePippin (Sage Science, Beverly, MA, USA). Briefly, 5 µg of DNA was diluted to 30 µL in TE buffer and 10 µL of RT equilibrated loading buffer was added to each aliquot and mixed by pipetting. Samples were loaded on the cassette by removing 40 µL of buffer from each well and adding 40 µL of sample or external marker. The cassette was run with the 0.75 % DF Marker U1 high-pass 30-40 kb v3 cassette definition. Size selected fractions (approximately 40 µL) were collected following the 30 min electrophoresis run. The library was prepared using the Chromium Genome Library Kit & Gel Bead Kit and sequenced (2 x 150 bp paired-end) on the NovaSeq 6000 (Illumina NovaSeq 6000 Sequencing System, <u>RRID:SCR_016387</u>) with NovaSeq 6000 SP Reagent Kit (300 cycles) and NovaSeq XP 2-Lane Kit for individual lane loading.

### Hi-C sequencing

Hi-C library preparation and sequencing was conducted at the Ramaciotti Centre for Genomics at the University of New South Wales (UNSW Sydney) using the Phase Genomics Plant kit v3.0. The library was assessed using Qubit and the Agilent 2200 TapeStation system (Agilent Technologies, Mulgrave, VIC, Australia). A pilot run on an Illumina iSeq 100 with 2 x 150 bp paired end sequencing run was performed for QC using hic_qc v1.0 (Phase Genomics, 2019) with i1 300 cycle chemistry. This was followed by sequencing on the Illumina NextSeq 500 (Illumina NextSeq 500, <u>RRID:SCR_014983</u>) with 2 x 150 bp paired-end high output run and NextSeq High Output 300 cycle kit v2.5 chemistry.

### Genome assembly and validation

Our assembly workflow consisted of assembling a draft long-read assembly, hybrid polishing of the assembly with long- and short-reads, and scaffolding the assembly into chromosomes using Hi-C data (Figure 2). Computational steps were carried out on the UNSW Sydney cluster Katana.

**Figure 2.**
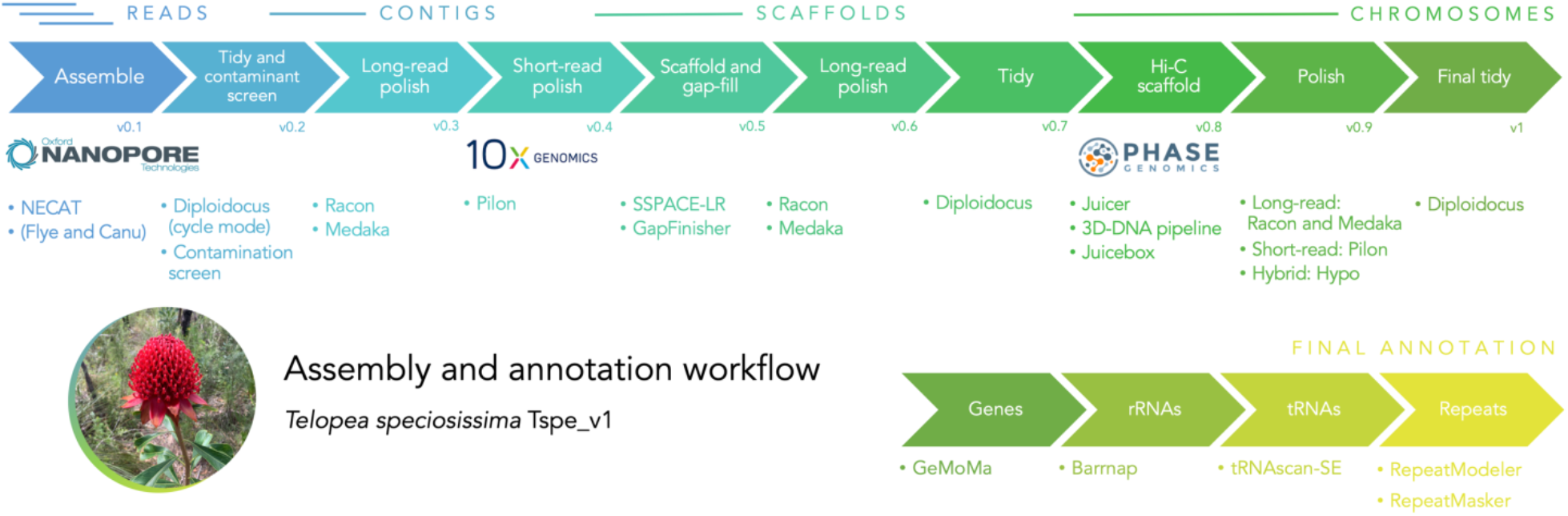
Assembly and annotation workflow for the *Telopea speciosissima* reference genome Tspe_v1. Logos reproduced with permission. Waratah photo by SH Chen.

The first stage of our assembly approach involved comparing three long-read assemblers using the ONT data as input: NECAT v0.01 (Chen et al., 2021), Flye (Flye, <u>RRID:SCR_017016</u>) v2.6 (Kolmogorov et al., 2019) and Canu (Canu, <u>RRID:SCR_015880</u>) v1.9 (Koren et al., 2017). The genome size parameter used for the assemblers was 1,134 Mb, as previously reported for *Telopea truncata* (Jordan et al., 2015). We later refined genome size estimates for *T. speciosissima* (see ‘DepthSizer: genome size estimation using single-copy orthologue sequencing depths’ section below). We chose the draft long-read assembly for use in downstream assembly steps based on contiguity (N50), BUSCO completeness and assembly size in relation to the DepthSizer estimated genome size. As a comparison to the long-read assemblies, the 10x data were assembled with Supernova (Supernova assembler, <u>RRID:SCR_016756</u>) v2.1.1 (Weisenfeld et al., 2017) with 332 Mb reads subsampled by Supernova (54x raw coverage, as recommended by Supernova documentation) as input. We generated pseudohaploid output (pseudohap2 output ‘1’).

### Assembly completeness and accuracy

Completeness was initially evaluated by BUSCO (BUSCO, <u>RRID:SCR_015008</u>) v3.0.2b (Simão et al., 2015), implementing BLAST+ v2.2.31, Hmmer v3.2.1 and EMBOSS v6.6.0 with the embryophyta_odb9 dataset (*n* = 1,440). To investigate the robustness of BUSCO completeness statistics, assemblies were also evaluated with BUSCO v5.0.0 (Manni et al., 2021), implementing BLAST+ v2.11.0 (Altschul et al., 1990), SEPP v4.3.10 (Mirarab et al., 2011) and Hmmer (Hmmer, <u>RRID:SCR_005305</u>) v3.3 (Eddy, 2011), against the embryophyta_odb10 dataset (*n* = 1,614). BUSCO results were calculated with both Augustus (Augustus, <u>RRID:SCR_008417</u>) v3.3.2 (Stanke & Morgenstern, 2005) and MetaEuk v732bcc4b91a08e69950ce0e25976f47c3bb6b89d (Levy Karin et al., 2020) as the gene predictor.

BUSCO results were collated using BUSCOMP (BUSCO Compilation and Comparison Tool; <u>RRID:SCR_021233</u>) v0.11.0 (Stuart et al., 2021) to better evaluate the gains and losses in completeness between different assembly stages, and compare different BUSCO versions. Assembly quality (QV) was also estimated using k-mer analysis of trimmed and filtered 10x linked-read data by Merqury v1.0 with k = 20 (Rhie et al., 2020). First, 30 bp from the 5’ end of read 1 and 10 bp from the 5’ end of read 2 were trimmed using BBmap (BBmap, <u>RRID:SCR_016965</u>) v38.51 (Bushnell, 2014). In addition, reads were trimmed to Q20, then those shorter than 100 bp were discarded.

### Genome size estimation and ploidy

*Telopea speciosissima* has been reported as a diploid (2*n* = 22) (Darlington & Wylie, 1956; Ramsay, 1963). We confirmed the individual’s diploid status using Smudgeplot v0.2.1 (Ranallo-Benavidez et al., 2019). The 1C-value of *T. truncata* (Tasmanian waratah) has been estimated at 1.16 pg (1.13 Gb) using flow cytometry (Jordan et al., 2015). We used the 10x data to estimate the genome size using Supernova v2.1.1 and GenomeScope (GenomeScope, <u>RRID:SCR_017014</u>) v1.0 (Vurture et al., 2017).

We sought to refine the genome size estimate of *T. speciosissima* using the ONT data and draft genome assemblies, implementing a new tool, DepthSizer (https://github.com/slimsuite/depthsizer, <u>RRID:SCR_021232</u>, **Box 1**). ONT reads were mapped onto each draft genome using Minimap2 (Minimap2, <u>RRID:SCR_018550</u>) v2.17 (Li, 2018) (--secondary=no -ax map-ont). The single-copy read depth for each assembly was then calculated as the modal read depth across single copy complete BUSCO genes, which should be reasonably robust to poor-quality and/or repeat regions within these genes (Edwards et al., 2021).

### DepthSizer benchmarking

DepthSizer was benchmarking using five PacBio reference genomes, plus the high-quality genome assembly and PacBio long reads for the German Shepherd Dog (Field et al., 2020; Table S1). Accuracy was calculated as the estimated genome size, divided by the documented genome size. Additional benchmarking of the robustness of DepthSizer predictions was performed using ONT and PacBio sequence data for three high-quality dog genomes: Basenji (Edwards et al., 2021), Dingo (Yadav et al., 2020), and German Shepherd Dog (Field et al., 2020). Raw reads from each technology were analysed independently using both the breed-specific reference genome, and the CanFam 3.1 dog reference (GCA_000002285.2; Lindblad-Toh et al., 2005). For all benchmarking, reads were mapped with Minimap2 (Minimap2, <u>RRID:SCR_018550</u>) v2.17 (Li, 2018). Summary violin plots were generated with ggstatsplot (Patil, 2021) in R.

#### Box 1.

**DepthSizer: genome size estimation using single-copy orthologue sequencing depths**

GitHub: https://github.com/slimsuite/depthsizer

Genome size prediction is a fundamental task in genome assembly. DepthSizer is a tool for estimating genome size using single-copy long-read sequencing depth profiles.

By definition, sequencing depth (X) is the volume of sequencing divided by the genome size. Given a known volume of sequencing, it is therefore possible to estimate the genome size by estimating the achieved sequencing depth. DepthSizer works on the principle that the modal read depth across single copy BUSCO genes provides a good estimate of the true depth of coverage. This assumes that genuine single copy depth regions will tend towards the same, true, single copy read depth. In contrast, assembly errors or collapsed repeats within those genes, or incorrectly-assigned single copy genes, will give inconsistent read depth deviations from the true single copy depth. The exception is regions of the genome only found on one haplotig – half-depth alternative haplotypes for regions also found in the main assembly – such as heterogametic sex chromosomes (Edwards et al., 2021), but these are unlikely to outnumber genes present in single copy on both homologous chromosomes. As a consequence, the dominant (i.e. modal) depth across these regions should represent single copy (2n) sequencing depth. First, the distribution of read depth for all single copy genes is generated using Samtools (Samtools, <u>RRID:SCR_002105</u>) v0.11 (Li et al., 2009) mpileup, and the modal peak calculated using a smoothed ‘density’ function of R (R Project for Statistical Computing, <u>RRID:SCR_001905</u>) v3.5.3 (R Core Team, 2019) to allow non-integer estimation (see DepthSizer documentation for details). Genome size, *G*, was then estimated from the modal peak single-copy depth, *X_sc_*, and the total volume of sequencing data, *T*, using the formula: *G* = *T / X_SC_*.

DepthSizer has six different genome size adjustment modes that modify *T* using different core assumptions (see documentation for details):

• None: no adjustment. Assumes zero contamination and perfect read mapping.
• IndelRatio: adjusts total sequencing volume for mismatch between read data being mapped and assembly coverage. Assumes no contamination in raw reads.
• CovBases: sets *T* as the total number of sequencing read bases covering the assembly. (Assembly Length x Mean depth)
• MapBases: sets *T* as the total number bases from sequencing reads mapped on to the genome. Assumes perfect mapping and all unmapped reads are contamination.
• MapAdjust: adjusts total sequencing volume by the ratio of mapped reads to mapped bases to account for depth losses during mapping. Assumes no contamination in raw reads.
• MapRatio: adjusts the MapBases by the IndelRatio sequencing:mapping bias.

It is expected that the true genome size should fall between IndelRatio (upper) and MapRatio (lower). CovBases should provide an absolute lower bound for genome size. If there is a very large difference between CovBases and MapBases, this could indicate a problem with the reads and/or assembly (e.g. some kind of incompatibility) and will result in a very inaccurate MapAdjust. If there is a very big difference between MapBases and None, this could indicate a very incomplete assembly, or a lot of contamination. In these cases, it is advisable to establish which before deciding which prediction size to use.

Benchmarking on PacBio data from six model organisms demonstrates robust genome size estimates, with a tendency to slightly overestimate genome size as expected (Figure 3, Table S1 and Table S2). Additional benchmarking on three high-quality canid genomes further revealed robustness to both assembly used (breed-specific genome versus CanFam v3.1) and sequencing technology (PacBio vs ONT), although PacBio data appears to over-estimate genome size more than ONT data (Figure S1).

**Figure 3.**
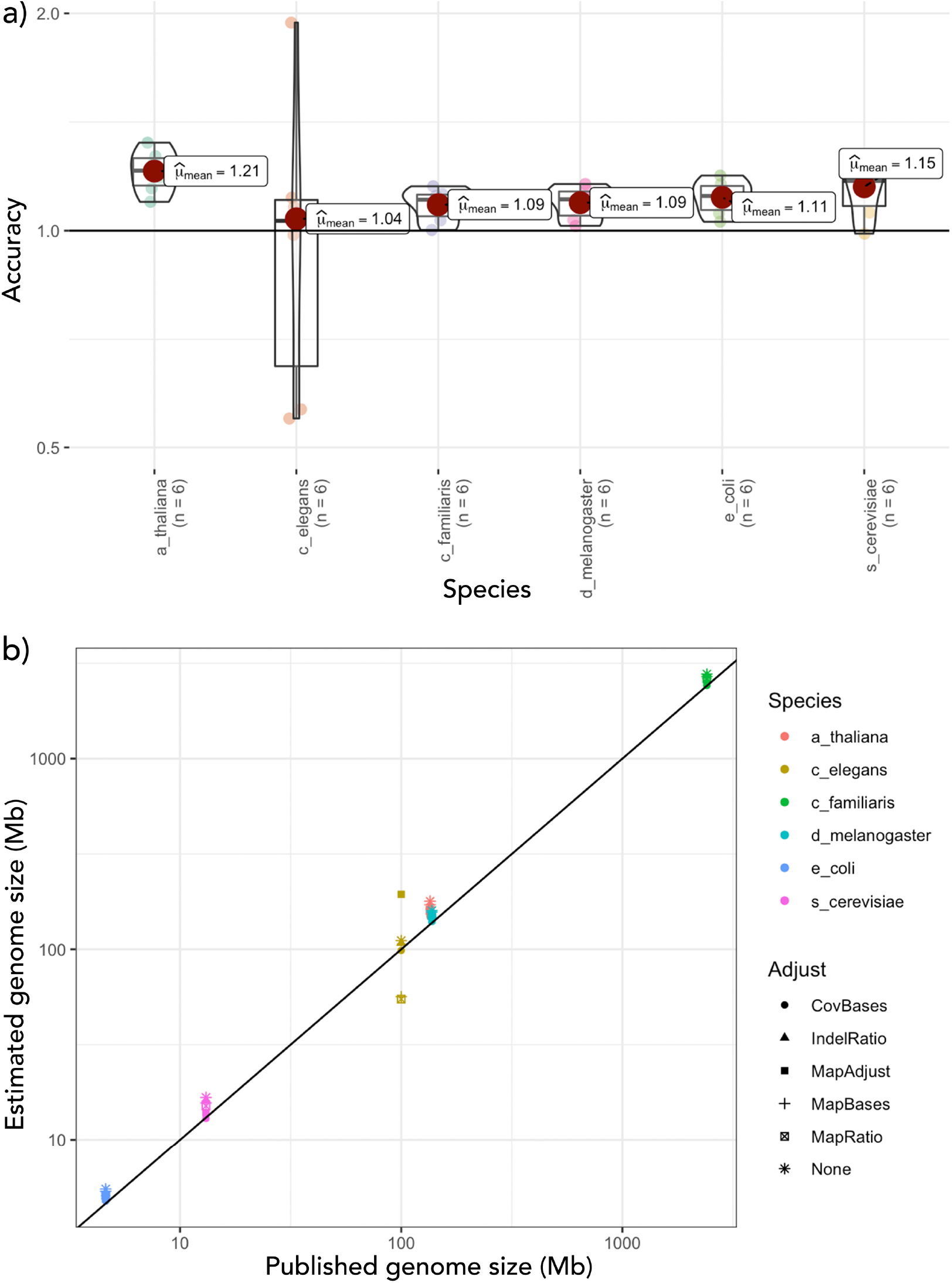
Model organism benchmarking of DepthSizer with six core methods using Minimap2 read mapping and Mpileup depth calculations with a) accuracy for each species and b) estimated genome size vs published genome size.

### Assembly tidying and contamination screening

The draft genome was screened and filtered to remove contamination, low-quality contigs and putative haplotigs, using more rigorous refinement of the approach taken for the Canfam_GSD (German Shepherd) and CanFam_Bas (Basenji) dog reference genomes (Edwards et al., 2021; Field et al., 2020), implemented in Diploidocus v0.9.6 (https://github.com/slimsuite/diploidocus, <u>RRID:SCR_021231</u>, **Box 2**).

BUSCO Complete genes were used to estimate a single-copy read depth of 54X. This was used to set low-, mid- and high-depth thresholds for Purge Haplotigs (Purge_haplotigs, <u>RRID:SCR_017616</u>) v20190612 (Roach et al., 2018) (implementing Perl v5.28.0, BEDTools (BEDTools, <u>RRID:SCR_006646</u>) v2.27.1 (Quinlan & Hall, 2010), R v3.5.3 (R Core Team, 2019), and SAMTools v1.9 (Li et al., 2009) of 13X, 40X and 108X. For the draft genome, convergence was reached after three cycles with 148 core sequences and 62 repeat sequences retained (see Table S6 for summary of cycles and Table S7 for full output).

#### Box 2.

**Automated genome assembly tidying with Diploidocus**

GitHub: https://github.com/slimsuite/diploidocus

Diploidocus is a tool that assists with tidying and curating genome assemblies. The tool combines read depth, KAT k-mer frequencies, Purge Haplotigs depth bins, Purge Haplotigs best sequence hits, BUSCO gene predictions, telomere prediction and vector contamination into a single seven-part (PURITY|DEPTH|HOM|TOP|MEDK|BUSCO+EXTRA) classification (Table S4). Diploidocus then performs a hierarchical rating of scaffolds, based on their classifications and compiled data (Table S5 and Figure 4). Based on these ratings, sequences are divided into sets:

1. Core. Predominantly diploid scaffolds and unique haploid scaffolds with insufficient evidence for removal.
2. Repeats. Unique haploid scaffolds with insufficient evidence for removal but dominated by repetitive sequences. High coverage scaffolds representing putative collapsed repeats.
3. Quarantine. Messy repetitive sequences and strong candidates for alternative haplotigs.
4. Junk. Low coverage, short and/or high-contaminated sequences.

**Figure 4.**
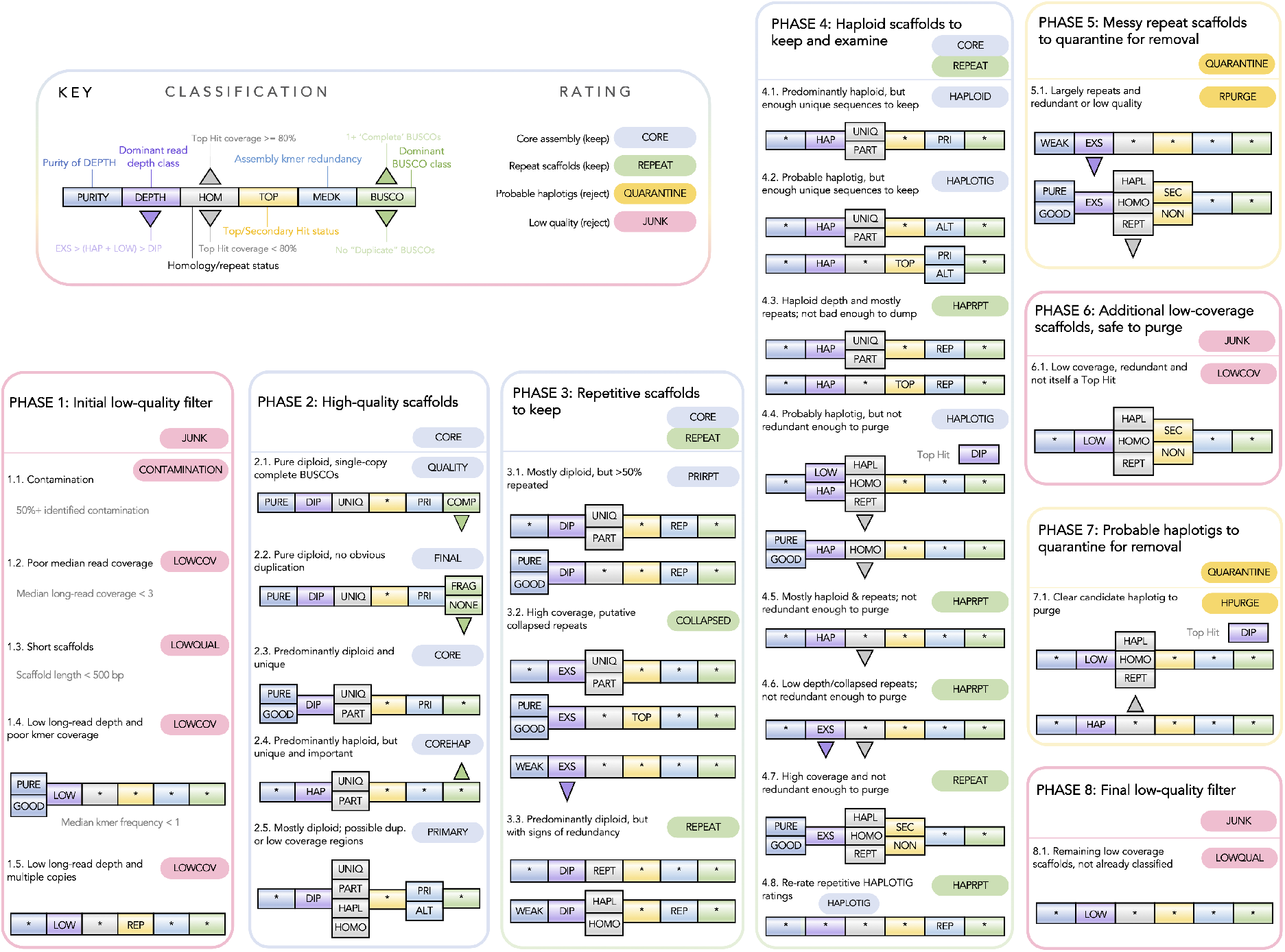
Diploidocus scaffold rating process based on a six-part classification. Asterisks indicate any class value is accepted. Phases are executed in order. Consequently, rules for later phases appear less restrictive than the full set of criteria required to receive that rating.

If any sequences are marked as ‘Quarantine’ or ‘Junk’, sequences in the ‘Core’ and ‘Repeat’ sets are retained and used as input for another round of classification and filtering.

First, the assembly is screened against the NCBI UniVec database (ftp://ftp.ncbi.nlm.nih.gov/pub/UniVec/, downloaded 05/08/2019) to identify and remove contaminants. Hits are first scored using rules derived from NCBI Vecscreen (https://www.ncbi.nlm.nih.gov/tools/vecscreen/) and regions marked as ‘Terminal’ (within 25 bp of a sequence end), ‘Proximal’ (within 25 bases of another match) or ‘Internal’ (>25 bp from sequence end or vecsreen match). Then, any segment of fewer than 50 bases between two vector matches or between a match and a sequence end are marked as ‘Suspect’. In our experience, default Vecscreen parameters appear prone to excessive false positives in large genomes (data not shown), and so Diploidocus features two additional contaminant identification filters. First, the ‘Expected False Discovery Rate’ (eFDR) is calculated for each contaminant sequence. This is simply the BLAST+ Expect value for that hit, divided by the total number of hits at that Expect value threshold. Any hits with an eFDR value exceeding the default threshold of 1.0 were filtered from the vecscreen results. Short matches in long-read assemblies are unlikely to be real contamination and a second filter was applied, restricting contaminant screening to a minimum hit length of 50 bp. Finally, the percentage coverage per scaffold is calculated from the filtered hits. This is performed first for each contaminant individually, before being collapsed into total non-redundant contamination coverage per query. Diploidocus then removes any scaffolds with at least 50 % contamination, trims off any vector hits within 1 kb of the scaffold end, and masks any remaining vector contamination of at least 900 bp. This masking replaces every other base with an N to avoid an assembly gap being inserted: masked regions should be manually fragmented if required. Diploidocus can also report the number of mapped long reads that completely span regions flagged as contamination.

After contamination screening, a sorted BAM file of ONT reads mapped to the filtered assembly is generated using Minimap2 v2.17 (−ax map-ont --secondary = no) (Li, 2018). Purge Haplotigs coverage bins were adjusted to incorporate zero-coverage bases, excluding assembly gaps (defined as 10+ Ns). Counts of Complete, Duplicate and Fragmented BUSCO genes were also generated for each sequence. General read depth statistics for each sequence were calculated with BBMap v38.51 pileup.sh (Bushnell, 2014). The sect function of KAT (KAT, <u>RRID:SCR_016741</u>) v2.4.2 (Mapleson et al., 2017) was used to calculate k-mer frequencies for the 10x linked reads (first 16 bp trimmed from read 1), and the assembly itself. Telomeres were predicted using a method adapted from https://github.com/JanaSperschneider/FindTelomeres, searching each sequence for 5’ occurrences of a forward telomere regular expression sequence, C{2,4}T{1,2}A{1,3}, and 3’ occurrences of a reverse regular expression, T{1,3}A{1,2}G{2,4}. Telomeres were marked if at least 50 % of the terminal 50 bp matches the appropriate sequence.

### Assembly polishing and gap-filling

The assembly was first long-read polished with Racon (Racon, <u>RRID:SCR_017642</u>) v1.4.5 (Vaser et al., 2017) using the parameters -m 8 -x -6 -g -8 -w 500 and medaka v1.0.2 (Oxford Nanopore Technologies Ltd., 2018) using the r941_prom_high_g303 model. Then, the 10x reads were incorporated by short-read polishing using Pilon (Pilon, <u>RRID:SCR_014731</u>) v1.23 (Walker et al., 2014) with reads mapped using Minimap2 v2.12 (Li, 2018) and correcting for indels only; we found correcting for indels only resulted in a higher BUSCO score than correcting for indels and SNPs following the steps described in this section. We scaffolded using SSPACE-LongRead v1.1 (Boetzer & Pirovano, 2014) with -k 1 followed by gap-filling using gapFinisher v20190917 (Kammonen et al., 2019) with default parameters. After another round of long-read polishing with Racon v1.4.5 (Vaser et al., 2017) and medaka v1.0.2 (Oxford Nanopore Technologies Ltd., 2018), we moved forward with a second round of tidying in Diploidocus v0.9.6 (default mode).

### Hi-C scaffolding

Hi-C data were aligned to the draft genome assembly using the Juicer (Juicer, <u>RRID:SCR_017226</u>) pipeline v1.6 (Durand et al., 2016) then scaffolds were ordered and orientated using the 3D *de novo* assembly pipeline (3D de novo assembly, <u>RRID:SCR_017227</u>) v180922 (Dudchenko et al., 2017). The contact map was visualised using Juicebox Assembly Tools v1.11.08 and errors over 3 review rounds were corrected manually to resolve 11 chromosomes (Dudchenko et al., 2018). The resulting assembly was tidied again using Diploidocus v0.10.6 (default mode).

### Final polishing and assembly clean-up

A further round of long-read polishing with Racon v1.4.5 (Vaser et al., 2017) and medaka v1.0.2 (Oxford Nanopore Technologies Ltd., 2018) was performed as described above. We then short-read polished using Pilon v1.23 (Walker et al., 2014). Two Pilon strategies were applied: (1) indel-only correction; (2) indel and SNP correction. We retained the indel and SNP corrected assembly as it resulted in a marginally higher BUSCO score compared to indel only correction (1311 vs 1310 complete BUSCOs); there was no change to contig nor scaffold numbers. A final hybrid polish was performed using Hypo v1.0.3 (Kundu et al., 2019). The assembly was concluded with a final tidy with Diploidocus v0.14.1 (default mode). All gaps in the assembly were then standardised to 100 bp.

Genome-wide heterozygosity was estimated using trimmed 10x reads with GenomeScope (Vurture et al., 2017) from the k-mer 20 histogram computed using Jellyfish (Jellyfish, <u>RRID:SCR_005491</u>) v2.2.10 (Marçais & Kingsford, 2011).

### Genome annotation

The genome was annotated using the homology-based gene prediction program GeMoMa (GeMoMa, <u>RRID:SCR_017646</u>) v1.7.1 (Keilwagen et al., 2019) with four reference genomes downloaded from NCBI: *Macadamia integrifolia* (SCU_Mint_v3, GCA_013358625.1), *Nelumbo nucifera* (Chinese Lotus 1.1, GCA_000365185.2), *Arabidopsis thaliana* (TAIR10.1, GCA_000001735.2) and *Rosa chinensis* (RchiOBHm-V2, GCA_002994745.2). The annotation files for *M. integrifolia* were downloaded from the Southern Cross University data repository (<u>doi.org/10.25918/5e320fd1e5f06</u>). *Macadamia* (Nock et al., 2020) and *Nelumbo* (Ming et al., 2013) genomes were chosen as they are related to *Telopea* i.e. in the order Proteales. The other two high-quality genomes represented the core eudicots and included the model flowering plant *Arabidopsis* (Lamesch et al., 2012) and *Rosa* (Hibrand Saint-Oyant et al., 2018) where the publication focused on genetic regulators of ornamental traits which is of interest for *Telopea*. Annotation completeness was assessed using BUSCO v3.0.2b and v5.0.0 in proteome mode.

Ribosomal RNA (rRNA) genes were predicted with Barrnap (Barrnap, <u>RRID:SCR_015995</u>) v0.9 (Seemann, 2018) and transfer RNAs (tRNAs) were predicted with tRNAscan-SE (tRNAscan-SE, <u>RRID:SCR_010835</u>) v2.05 (Lowe & Chan, 2016), implementing Infernal (Infernal, <u>RRID:SCR_011809</u>) v1.1.2 (Nawrocki & Eddy, 2013). A set of 2,419 tRNAs was initially predicted and filtered to 760 using the recommended protocol for eukaryotes. Then, 22 tRNAs with mismatched isotype and 10 with unexpected anticodon were removed to form the high-confidence set.

The genome has also been annotated by the NCBI Eukaryotic Genome Annotation Pipeline using RNAseq data from other Proteaceae (RefSeq accession GCF_018873765.1).

### Genome-wide copy number analysis

Estimated single-copy (2n) sequencing depth was calculated for different regions of the genome using the same smoothed density profile as employed by DepthSizer (Box 1) and comparing this to the BUSCO-derived single-copy (2n) sequencing depth of DepthSizer. This analysis was performed on: (1) BUSCO v5 (MetaEuk) single-copy ‘Complete’ genes; (2) BUSCO v5 ‘Duplicated’ genes; (3) All NCBI gene annotations; (4) Each final assembly scaffold; (5) 100 kb non-overlapping windows across the genome. For convenience, this method has been made available as DepthKopy (https://github.com/slimsuite/depthkopy).

### Repeat annotation

Following the approach from the *Macadamia integrifolia* genome paper (Nock et al., 2020), we identified and quantified repeats in the Telopea genome as well as the other four species used in the GeMoMa annotation for comparison. A custom repeat library was generated with RepeatModeler (RepeatModeler, <u>RRID:SCR_015027</u>) v2.0.1 (-engine ncbi) and the genome was masked with RepeatMasker (RepeatMasker, <u>RRID:SCR_012954</u>) v4.1.0 (Tarailo-Graovac & Chen, 2009), both with default parameters. The annotation table was generated using the buildSummary.pl RepeatMasker script.

### Orthologous clusters and synteny analyses

Synteny between the *Telopea* (Tspe_v1) and *Macadamia* (SCU_Mint_v3) genomes was explored with satsuma2 version untagged-2c08e401140c1ed03e0f with parameters -l 3000 - do_refine 1 -min_matches 40 -cutoff 2 -min_seed_length 48 and visualised with the ChromosomePaint function (Grabherr et al., 2010) and MizBee v1.0 (Meyer et al., 2009). The protein sequences of Tspe_v1 and the four species used in the GeMoMa annotation were clustered into orthologous groups and tests for gene ontology (GO) enrichment were conducted for waratah-specific clusters using OrthoVenn2 (Xu et al., 2019). Intersection of clusters was visualised using the R package UpSetR (Conway et al., 2017).

### *CYCLOIDEA* transcription factor gene family analysis

Complete and partial protein sequences for *CYCLOIDEA* transcription factors were downloaded from NCBI using identifiers listed in Table S3 of Citerne et al., 2017. GABLAM v2.30.5 (Davey et al., 2006) was used to identify all homologous proteins (BLAST+ v2.11.0, blastp e-value <1e-4) in the waratah GeMoMa annotation, which was annotated with protein descriptions from closest Swissprot hits using SAAGA v0.7.6 (Stuart et al., 2021). Each *Telopea speciosissima* homologue was then used as query sequence for HAQESAC v1.14.0 (Edwards et al., 2007) to generate a high-quality multiple sequence alignment and inferred phlyogenetic tree of close homologues (limited to a maximum of 100 closest hits). A search database was constructed from all angiosperm proteins in Uniprot (taxid 3398), the three reference proteomes used for GeMoMa annotation (*Macadamia integrifolia*, *Nelumbo nucifera* and *Rosa chinensis*), and all angiosperm reference proteomes from Quest For Orthologues (March 2021 release; (Forslund et al., 2018). To this were added the original CYC sequences and full GeMoMa annotation of *T. speciosissima*. BLAST+ searches and HAQESAC runs were controlled by MultiHAQ v1.5.0 (Jones et al., 2011). To generate a comprehensive but non-redundant tree of *CYC* genes, all homologues meeting initial HAQESAC screening criteria (min 40 % global identity and 60 % global coverage to query, <50 % gaps relative to nearest homologue) were combined into a single non-redundant dataset of *CYCLOIDEA* homologues and their homologues. A candidate *Telopea CYCLOIDEA*-like 1 gene (TSPEV1G03060) was identified based on SAAGA annotation and HAQESAC homologues. This was used as a query for a second, manually curated HAQESAC run against the full non-redundant protein dataset, screening out any proteins with an unknown species designation (including sequence assigned the 9MAGSP species code). Multiple sequence alignments were performed with Clustal Omega (Clustal Omega, <u>RRID:SCR_001591</u>) v1.2.4 (Sievers et al., 2011). The final tree was generated with IQ-TREE (IQ-TREE, <u>RRID:SCR_017254</u>) v2.0.4 (Nguyen et al., 2015) with 1,000 bootstraps.

## RESULTS AND DISCUSSION

### High-quality chromosome-level Tspe_v1 reference genome

The ONT, 10x and Hi-C sequencing yielded a total of 48.3, 123.4 and 25.0 Gb of sequence, respectively (Table 1). At the initial long-read assembly stage, NECAT resulted in the most contiguous assembly, at 365 contigs and the highest BUSCO completeness at 81.2 %. This was followed by Flye at 2,484 contigs and 81.0 % complete, then Canu at 3,983 contigs at 78.4 % complete. The BUSCO completeness of the 10x pseudohaploid assembly was higher than each of the long-read assemblies at 91.8 %. However, the 10x assembly had much lower contiguity at 43,951 contigs, as expected (Table S3). Whilst Supernova had a higher BUSCO completeness (91.9 % versus 81.2 %), NECAT was orders of magnitude better in terms of contiguity (10.7 Mb N50 on 365 contigs vs 874 kb N50 on 27,610 scaffolds). Furthermore, BUSCOMP analysis revealed that the NECAT assembly contained more complete BUSCO genes when base accuracy is not considered (Figure 5; Supplementary Files – BUSCOMP full report). Guided by these metrics, NECAT was selected as the core assembly for additional processing. We confirmed the individual’s diploid status with Smudgeplot (Figure S2a).

**Figure 5.**
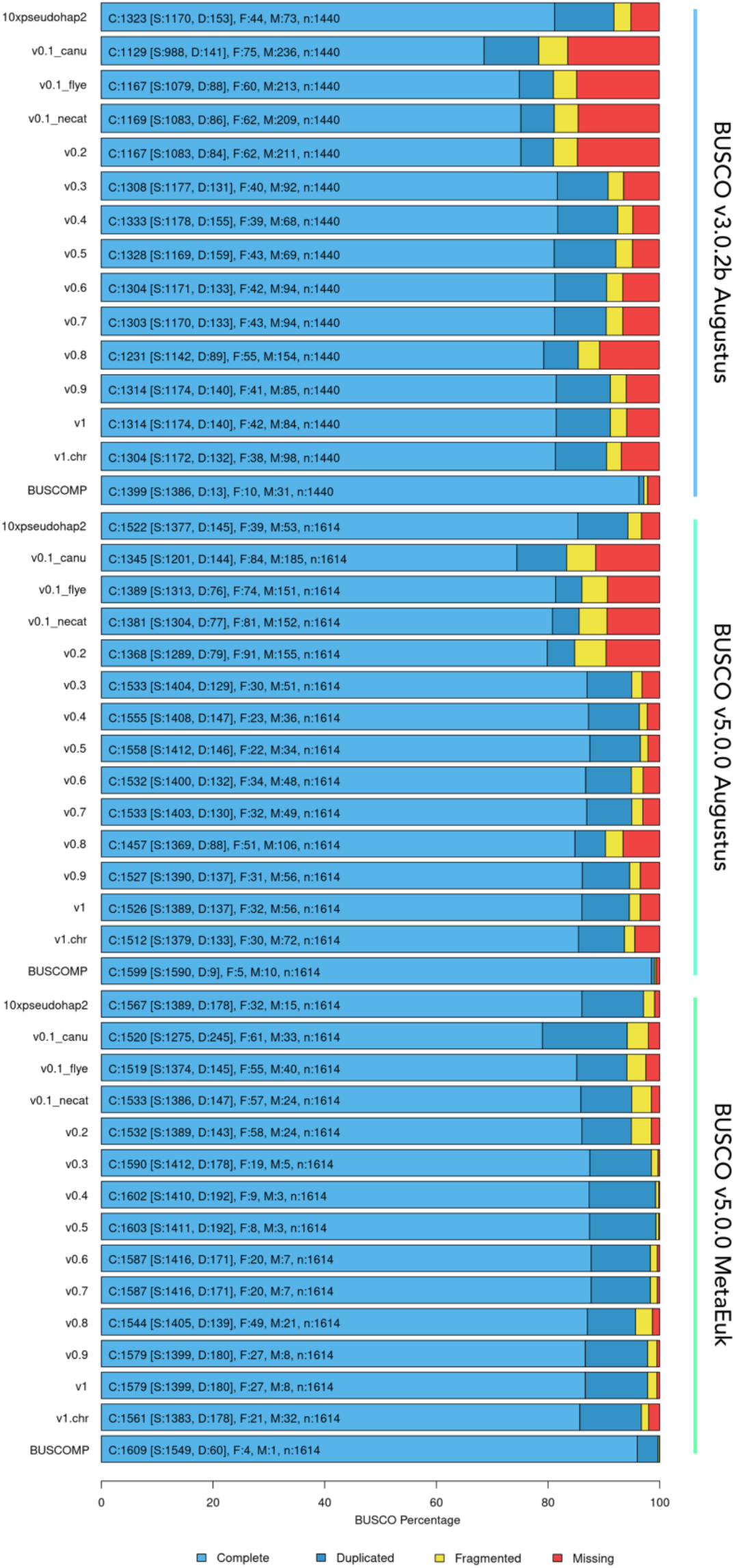
BUSCOMP summary of BUSCO completeness rating compiled over different stages (see Figure 2) of the *Telopea speciosissima* genome assembly. The final BUSCOMP rating uses the best rating per BUSCO gene across any of the assemblies.

**Table 1.** Library information of *Telopea speciosissima* reference genome (Tspe_v1).

| Sequencing platform | Library | Median insert size (bp) | Mean read length (bp) | No. of reads | Sequence bases (Gb) |
| --- | --- | --- | --- | --- | --- |
| Oxford Nanopore Technologies<sup>†</sup> | Ligation (SQK-LSK109) | - | 13,449 | 3,595,148 | 48.3 |
| Illumina NovaSeq 6000 | Paired-end 10x Chromium | 336 | 2 x 150 | 822,558,750 | 123.4 |
| **Total gDNA** | - | - | - | **826,153,898** | **171.7** |
| Illumina NextSeq 500<sup>‡</sup> | Phase Genomics<br>Proximo Hi-C (Plant) | 174 | 2 x 151 | 165,573,702 | 25.0 |

<sup>†</sup> Two PromethION flow cells and two partial flow cells from a MinION pilot run
<sup>‡</sup> Includes a pilot iSeq run used to QC the library

Rounds of polishing and tidying improved the contiguity and quality of the genome as the genome progressed through the assembly workflow (Table S3). The first round of polishing markedly improved the BUSCO score – long-read polishing increased complete BUSCOs from 1,532 (v0.2) to 1,590 (v0.3) and short-read polishing further increased this to 1,602 (v0.4). The assembly was scaffolded by SSPACE-LongRead from 209 contigs into 138 scaffolds, however, no gaps were filled by gapFinisher. After further long-read polishing, a run of Diploidocus (v0.7) retained 128 scaffolds out of 138, which consisted of 87 core, 41 repeat, 10 quarantine and 0 junk scaffolds. Following incorporation of Hi-C data, the assembly was in 2,357 scaffolds, and the N50 increased substantially from 16.5 Mb to 68.9 Mb. Surprisingly, the contig number increased considerably from 148 to 3,537, suggesting that the Hi-C data and NECAT assembly were frequently in conflict. The resulting assembly was tidied with Diploidocus and 1643 scaffolds (824,534,974 bp) were retained out of 2,357 (833,952,765 bp; 1,347 core, 296 repeat, 548 quarantine and 166 junk scaffolds). The removal of many sequences by Diploidocus, and the less contiguous initial assemblies from widely-used long-read assemblers Canu and Flye (Table S3), suggest that the NECAT assembly contained erroneously joined sequences, and these were corrected by Hi-C. However, it is also possible that limitations of the Hi-C library contributed to the high degree of fragmentation. The assembly contiguity improved to 1,399 scaffolds and 1,595 contigs following a further round of long-read polishing (Table S3). Following hybrid polishing with Hypo (v0.9), the number of scaffolds remained as 1,399 and the BUSCO score improved slightly. Notably, Hypo polishing improved the Merqury QV score from 29.8 to 33.9. A final iteration of Diploidocus Tidy removed 72 putative haplotigs and 38 low quality ‘junk’ scaffolds, keeping 1,084 core and 250 repetitive scaffolds.

The conclusion of the assembly workflow produced an 823.3 Mb haploid genome assembly (Tspe_v1) on 1,289 scaffolds, with an N50 of 69.0 Mb and L50 of 6 (Table 2). The Hi-C data facilitated scaffolding into 11 chromosomes (Figure 6), conforming to previous cytological studies (Darlington & Wylie, 1956), and the anchored proportion of Tspe_v1 spanned 94.2 % of the final assembly; the chromosomes were numbered by descending length (Table S8) as this is the first instance *Telopea* chromosomes have been studied in detail.

**Figure 6.**
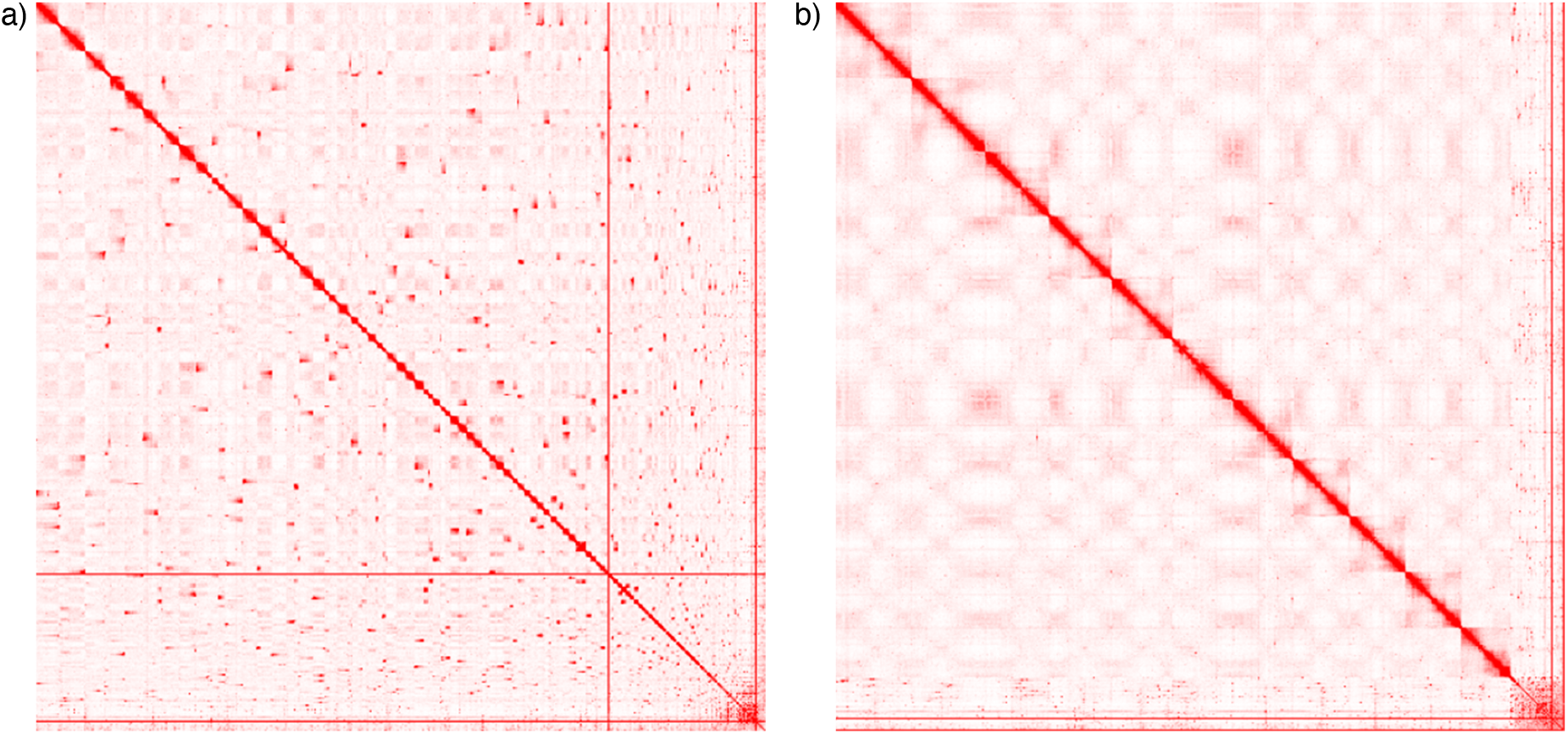
Hi-C contact matrices visualised in Juicebox.js in balanced normalisation mode a) before and b) after correction.

**Table 2.** Genome assembly and annotation statistics for the *Telopea speciosissima* reference genome.

| Statistic | Tspe_v1 |
| --- | --- |
| **Total length (bp)** | **823,061,212** |
| **No. of scaffolds** | **1,289** |
| N50 (bp) | 69,013,595 |
| L50 | 6 |
| **No. of contigs** | **1,452** |
| N50 (bp) | 12,206,888 |
| L50 | 21 |
| N bases | 18,174 |
| GC (%) | 40.11 |
| **BUSCO<sup>†</sup> complete (genome; n = 1,614)** | **97.8 % (1,579)** |
| Single copy (genome) | 86.7 % (1,399) |
| Duplicated (genome) | 11.2 % (180) |
| BUSCO fragmented (genome) | 1.7 % (27) |
| BUSCO missing (genome) | 0.5 % (8) |
| **Protein-coding genes** | **40,158** |
| mRNAs | 46,877 |
| rRNAs | 351 |
| tRNAs | 728 |
| **BUSCO<sup>†</sup> complete (proteome; n = 1,614)** | **94.4 % (1,524)** |
| Single copy (proteome) | 82.7 % (1,334) |
| Duplicated (proteome) | 11.8 % (190) |
| BUSCO fragmented (proteome) | 3.2 % (52) |
| BUSCO missing (proteome) | 2.4 % (38) |

<sup>†</sup> BUSCO v5 MetaEuk (embryophyta\_odb10)

From a core set of 1,614 single-copy orthologues from the Embryophyta lineage, 97.8 % were complete in the assembly (86.7 % as single-copy, 11.2 % as duplicates), 1.7 % were fragmented and only 0.5 % were not found, suggesting that the assembly includes most of the waratah gene space. Interestingly, BUSCO scores vary by many percentages between different BUSCO versions and gene predictors. BUSCO v5.0.0 with MetaEuk as the gene predictor consistently produced the highest scores (Table S3). BUSCO v3.0.2b with Augustus benchmarked the assembly against 1,440 single-copy orthologues only found 91.3 % complete in the assembly (81.5 % as single-copy, 9.7 % as duplicates), with 2.9 % fragmented and 5.8 % missing. BUSCO v5.0.0 with Augustus as the gene predictor reported higher scores than v3.0.2b but lower than when MetaEuk was used as the gene predictor (Table S3). We recovered a maximal non-redundant set of 1,549 complete single copy BUSCOs across the set of assemblies. BUSCOMP analysis revealed that only one gene out of 1614 was not found by BUSCO v5 MetaEuk in any version of the assembly (Figure 5; Supplementary File – BUSCOMP full report). The Tspe_v1 assembly completeness is favourable in comparison to the *Macadamia integrifolia* (SCU_Mint_v3) assembly (Nock et al., 2020), which also combined long-read and Illumina sequences (BUSCO v5 MetaEuk 96.7 % vs 81.9 % complete, respectively, in the anchored portion of the assembly). The Merqury QV score of the assembly was 34.03, indicating a base-level accuracy of >99.99 % (Figure S3). Genome-wide heterozygosity was estimated to be 0.756 % (Figure S2b).

### The *Telopea speciosissima* genome is approximately 900 Mb

The 1C-value of *T. truncata* (Tasmanian waratah) has been estimated at 1.16 pg (1.13 Gb) using flow cytometry (Jordan et al., 2015). Supernova v2.1.1 predicted a genome size of 953 Mb from the assembly of the 10x linked-reads whilst GenomeScope predicted a smaller genome of 794 Mb from the same data (Figure S2b). DepthSizer analysis of the six different versions of the genome assembly (four raw assemblies, Tspe_v1, and Tspe_v1 chromosomes) estimated the genome size of *T. speciosissima* to fall within a range from 850 Mb to 950 Mb (Table S9), and shows good robustness to both assembly version and BUSCO dataset used (Figure 7). This falls between the Supernova and GenomeScope estimates. We report an estimated genome size of approximately 900 Mb, considering the mean of estimates of the six adjustment methods using the BUSCO v5 MetaEuk data, based on the highest quality Tspe_v1 assemblies.

**Figure 7.**
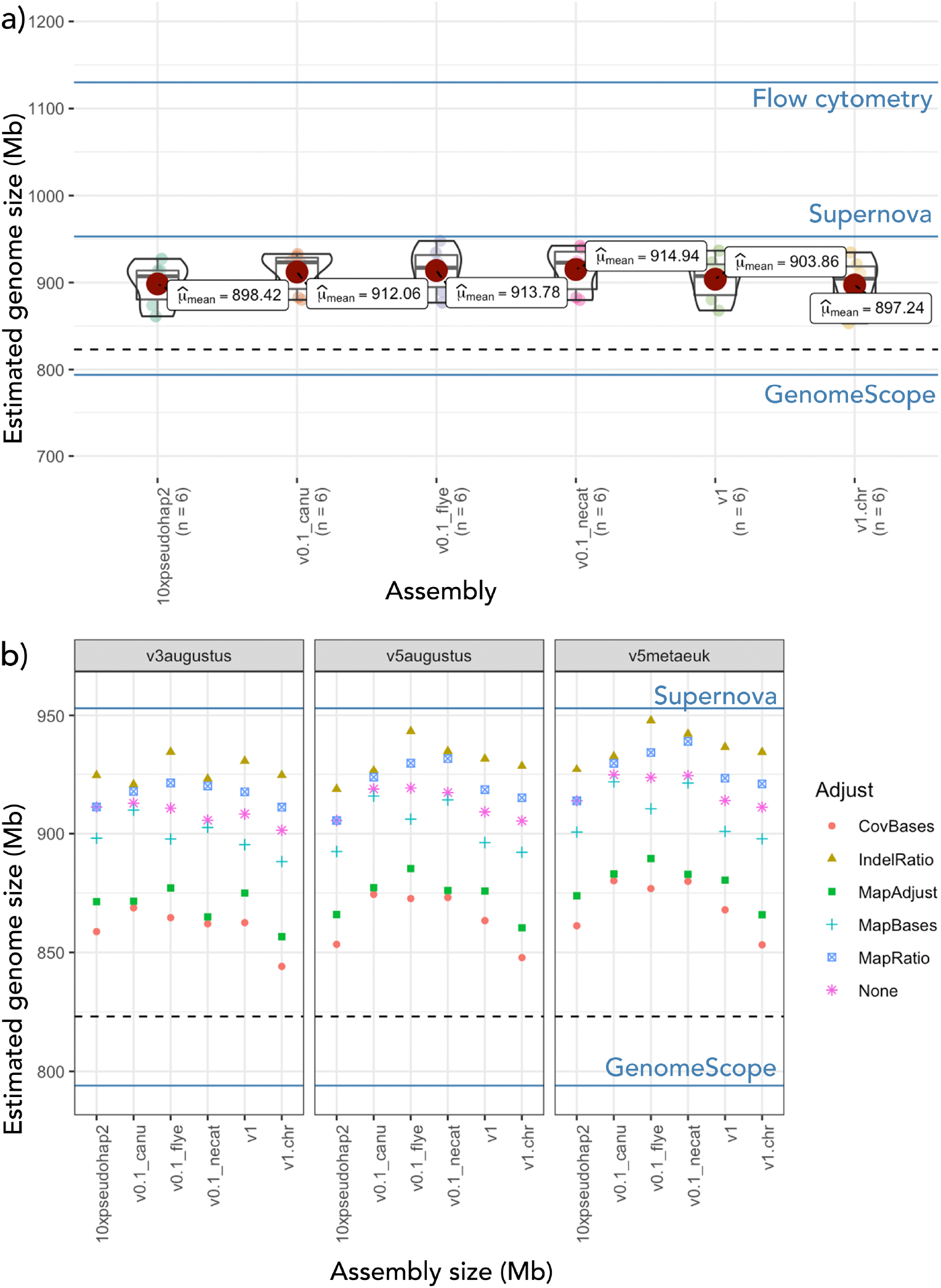
DepthSizer *Telopea* assembly size prediction using read depth of BUSCO v5 MetaEuk genes a) sits between estimates from flow cytometry, Supernova and GenomeScope at mean of 904 Mb for the v1 final assembly and is b) robust to BUSCO versions, with variation across the four adjustment methods. Dotted line represents the final assembly size.

### The majority of Tspe_v1 is at single-copy (2*n*) read depth

Read depth copy number analysis reveals that the majority of the assembly is at the expected 2*n* depth (Figure 8). Single-copy ‘Complete’ BUSCO genes strongly cluster around CN = 1, further supporting the robustness of the method underpinning DepthSizer. Notably, the 180 ‘Duplicated’ BUSCO genes are also predominantly at single-copy depth, with a similar copy number distribution to the BUSCOs classified as single-copy and complete. This indicates that the vast majority are likely to be real duplications found in *T. speciosissima*, with only a few representing potential sequencing errors (Table S10). This was supported by HAQESAC phylogenetic analysis of all 180 genes (Supplementary File – Tspe_v1.buscodup_HAQESAC.zip). Copy number analysis of all 14,882 NCBI annotated genes shows a similar clustering around a median copy number of 1. However, the mean copy number is surprisingly high at 2.36. Further inspection of the data revealed that this is being driven by a reasonably small number of very high copy number genes, derived from highly collapsed repeat regions (Table S11). This is further supported by the elevated mean copy number for both whole scaffolds and 100 kb windows. This is consistent with the identification by Diploidocus of 250 repetitive scaffolds, and a final assembly of approx. 91.5 % of the predicted genome size. Consistent with other Hi-C scaffolded assemblies (e.g. Rhie et al., 2021), it is likely that Tspe_v1 still contains some misassemblies that will need to be corrected with additional curation in future.

**Figure 8.**
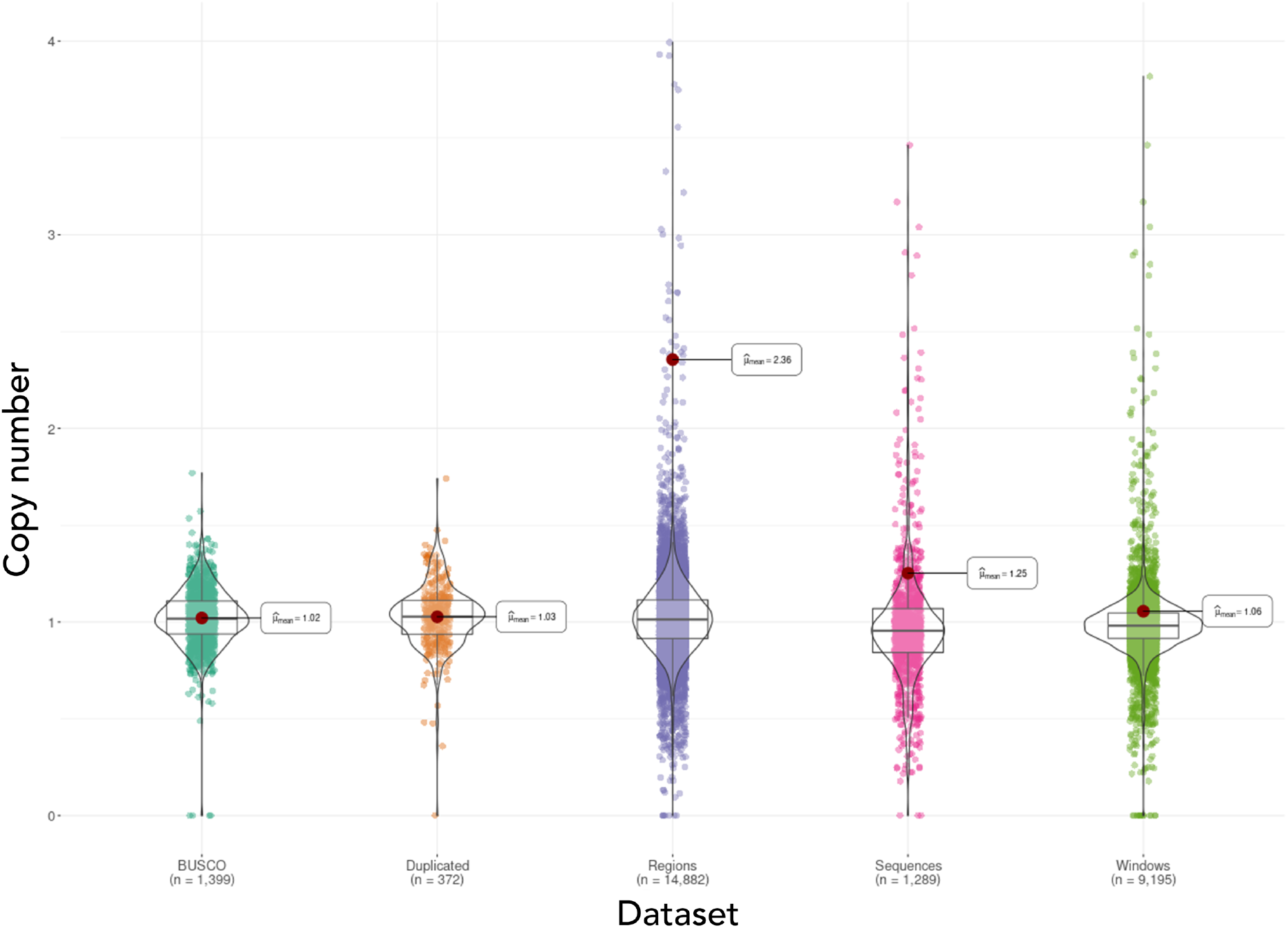
Genome-wide regional copy number analysis. Copy number (CN) is relative to a single diploid (2n) copy in the genome, truncated at CN = 4. Violin plots and means generated with ggstatsplot. Each data point represents a different genomic region. BUSCO, BUSCO v5 (MetaEuk) single-copy ‘Complete’ genes; Duplicated, BUSCO v5 ‘Duplicated’ genes; Regions, NCBI gene annotations; Sequences, assembly scaffolds; Windows, 100 kb non-overlapping windows across the genome.

### Repetitive elements and gene prediction

The *Telopea* genome is highly repetitive, with repeats accounting for 62.3 % of the total sequence length and has a similar repeat content to *Macadamia*, previously reported as 55.1 % (Nock et al., 2020) and found to be 58.5 % in our analyses (Table S12). Class I transposable elements (TEs) or retrotransposons were the most pervasive classified repeat class (20.3 % of the genome) and were dominated by long terminal repeat (LTR) retrotransposons (18.1 %). Class II TEs (DNA transposons) only accounted for 0.03 % of the genome. A high percentage of repeats remained unclassified (40.6 %) and the genome will serve as a resource for future studies into repetitive elements in *Telopea* and related species.

Genome annotation predicted 40,126 protein-coding genes and 46,842 mRNAs in the *T. speciosissima* assembly, which fits the expectation for plant genomes (Sterck et al., 2007). Of these genes, 38,427 appeared in the 11 chromosomes (Table S8). Of 1,440 Embryophyta orthologous proteins, 94.0 % were complete in the annotation (79.3 % as single-copy, 14.7 % as duplicates), 3.4 % were fragmented and 2.6 % were missing. Additionally, 351 rRNA genes and a set of 728 high-confidence transfer RNAs (tRNAs) were predicted. The NCBI Annotation Release 100 had a higher completeness, as expected, than the GeMoMa annotation; of 1,614 Embryophyta genes, 98.3 % were complete in the annotation (54.2 % as single-copy, 44.1 % as duplicated), 1.1 % were fragmented and 0.6 % were missing. When comparing the assembly completeness with proteome completeness using BUSCO v3.0.2b, the proteome completeness at 94.0 % (79.3 % as single-copy and 14.7 % as duplicated) was unexpectedly higher than the genome completeness at 91.3 % (81.5 % as single-copy and 9.7 % as duplicated). However, this issue was resolved with a later version of BUSCO (v5.0.0). The improvements in BUSCO likely meant that genes could be better discerned in the genome assembly, where they are more difficult to identify, compared to a proteome. An inverse pattern in the incidence of genes and repeats was observed across all chromosomes, with repeat content generally peaking towards the centre of each chromosome (Figure 9), suggesting predominantly metacentric and submetacentric chromosomes. This pattern may represent enriched repeat content and reduced coding content in pericentromeric regions, although further study is required to identify the centromeres (Jiang et al., 2003; Oliveira & Torres, 2018; Simon et al., 2015).

**Figure 9.**
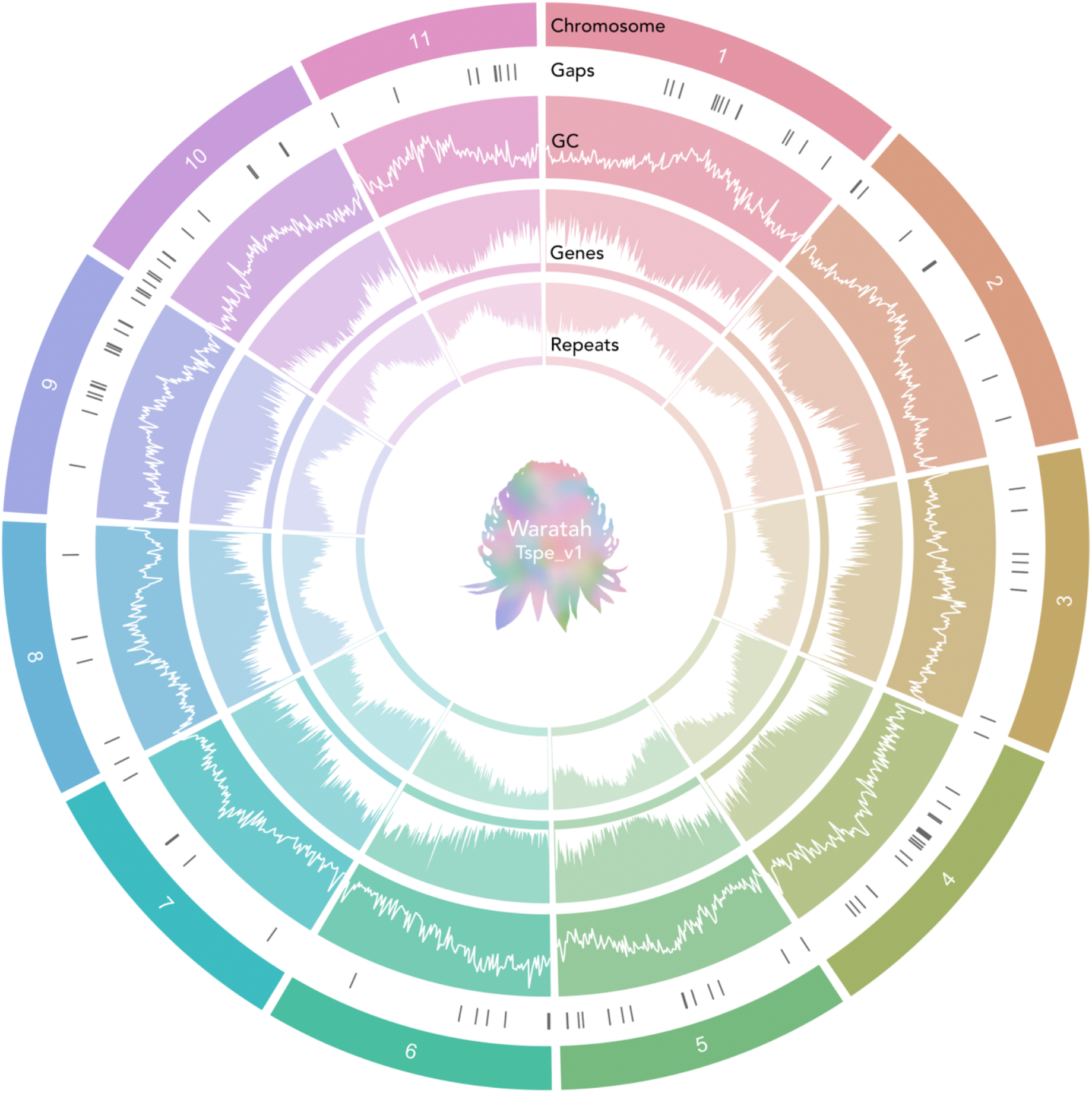
Features of the 11 chromosomes of the *Telopea speciosissima* reference genome. Concentric tracks from the outside inward represent: chromosomes, gaps (gaps of unknown length appear as 100 bp in the assembly), GC content calculated using BEDTools v2.27.1 (Quinlan & Hall, 2010), gene density and repeat density. The latter three tracks denote values in 500 kb sliding windows. Density was defined as the fraction of a genomic window that is covered by genomic regions. Plots are white on a solid background coloured by chromosome. Visualisation created using the R package circlize v0.4.12 (Gu et al., 2014).

### BUSCO completeness statistics must be matched by version and gene predictor

One surprising observation from our BUSCO analysis was a jump in completeness of over 6 % when moving from BUSCO v3 Augustus predictions to BUSCO v5 MetaEuk predictions (Figure 5 and Table S3). This is explained in part by the change to the lineage database used. However, completeness scores for BUSCO v5 Augustus are only about 3 % higher. This is particularly pronounced for the raw assemblies, where Augustus scores can be over 10 % lower than MetaEuk scores. Great care must be taken in naïve comparison of published BUSCO scores, even if using the same version of BUSCO. MetaEuk scores seem to be both higher and more stable. However, nucleotide sequences for Complete BUSCO genes are currently only output from Augustus mode. We have therefore updated BUSCOMP to extract the missing sequences from MetaEuk runs so that they can be used with downstream tools such as BUSCOMP that require these sequences.

### Orthologous clusters and synteny between *Telopea* and *Macadamia*

The five species formed 24,140 clusters: 23,031 orthologous clusters (containing at least 2 species) and 1,109 single-copy gene clusters. There were 9,463 orthologous families common to all of the species. The three members of the order Proteales (*T. speciosissima*, *M. integrifolia* and *N. nucifera*) shared 456 families (Figure 10 and Figure S4). Tests for GO enrichment of 912 waratah-specific clusters identified 12 significant terms (Table S13). The most enriched GO terms were DNA recombination (GO:0006310, P = 1.8 x 10^-27^), retrotransposon nucleocapsid (GO:0000943, P = 3.5 x 10^-12^) and DNA integration (GO:0015074, P = 4.1 x 10^-11^).

**Figure 10.**
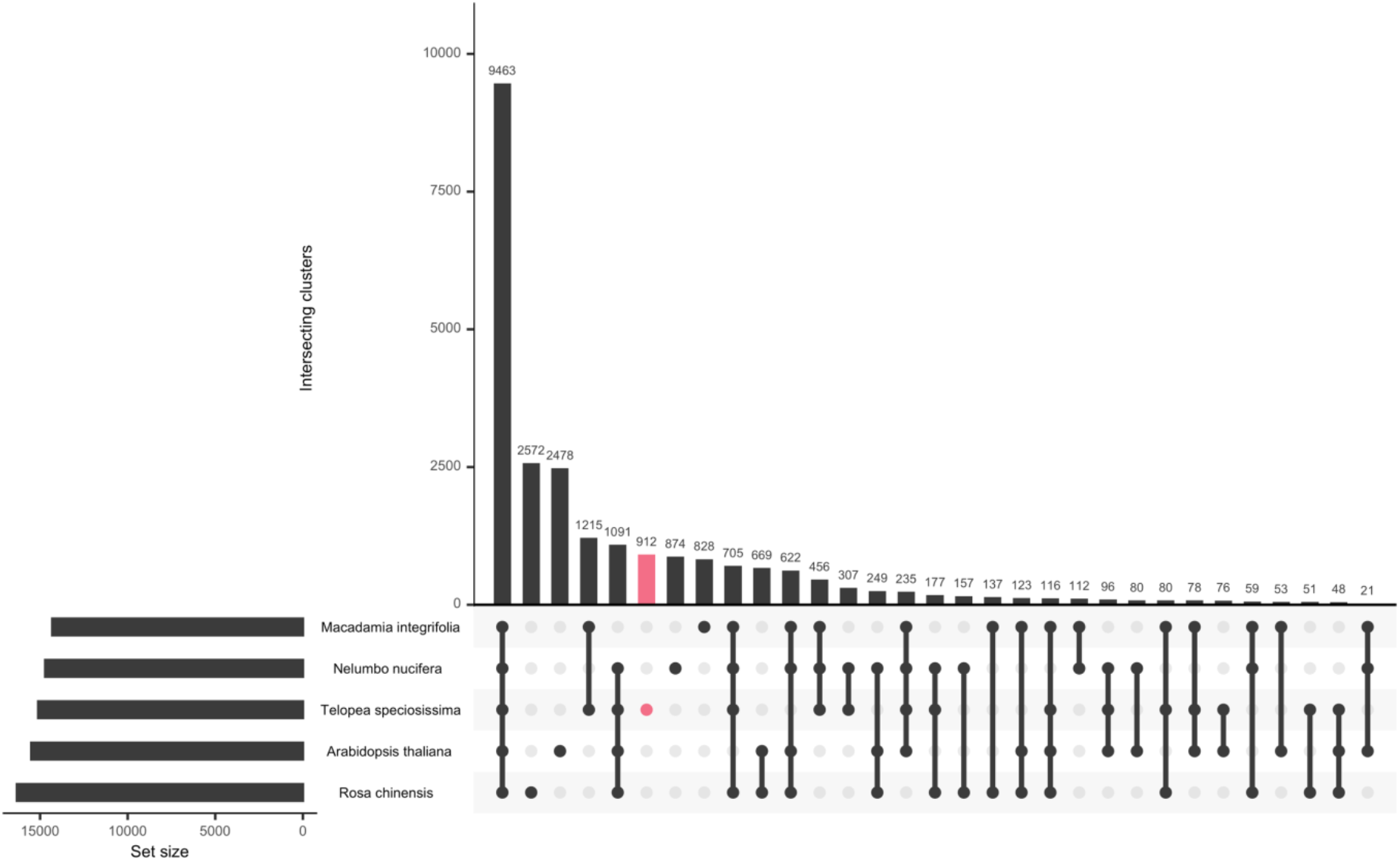
Orthologous gene clusters shared among the three members of the order Proteales – *Telopea speciosissima*, *Macadamia integrifolia* and *Nelumbo nucifera* – and the core eudicots – *Arabidopsis thaliana* (Brassicales) and *Rosa chinensis* (Rosales).

The *Macadamia* genome (2*n* = 28) has six more chromosomes than the *Telopea* genome (2*n* = 22), but the two species have similar estimated genome sizes – 896 Mb (Nock et al., 2020) compared to 874 Mb. It is thought that the ancestral Proteaceae had a chromosome number of *x* = 7 (Carta et al., 2020; L. A. S. Johnson & Briggs, 1963, 1975; Murat et al., 2017), although the occurrence of paleo-polyploidy in family has been debated (Stace et al., 1998). Overall, synteny analyses reveal an abundance of interchromosomal rearrangements between the *Telopea* and *Macadamia* genomes (Figure 11), reflecting the long time since their divergence (73-83 Ma; Sauquet et al., 2009). However, a number of regions exhibit substantial collinearity, for example, *Telopea* chromosome 09 and *Macadamia* chromosome 11 (Figure S5).

**Figure 11.**
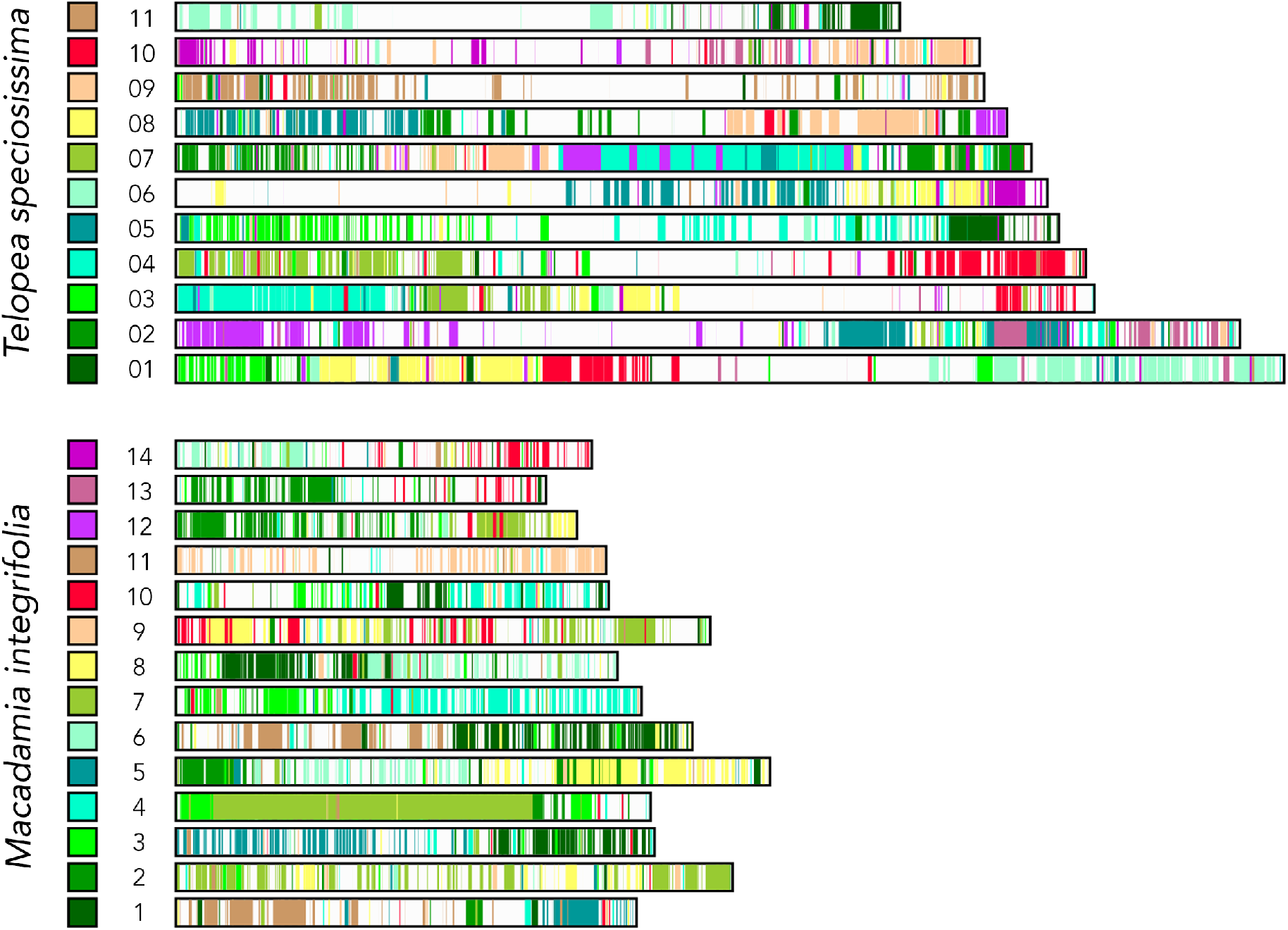
Synteny between *Telopea speciosissima* (2*n* = 22) and *Macadamia integrifolia* (2*n* = 28). Coloured squares for each species match painted chromosome regions in the other species. More detail of the underlying synteny and rearrangements can be found in Figure S5.

### *CYC* gene copy number and the genetic control of floral symmetry

In total, 210 predicted waratah sequences (longest isoform per gene) were identified as homologous to the 49 Citerne et al. *CYC* protein sequences. Of these, 198 generated multiple sequence alignments and phylogenetic trees. These combined to form a non-redundant dataset of 12,238 proteins. HAQESAC reduced this to a high-quality alignment of 46 homologous proteins, including two waratah proteins, TSPEV1G03060 – *CYC1* and TSPEV1G20406 – *CYC2.* Consistent with previous work (Citerne et al., 2017), these two proteins belonged to two distinct clades (Figure 12). While the exact role of the two paralogues in determining floral symmetry in Proteaceae would require a study of gene expression and remains incompletely understood in the species examined so far (Citerne et al., 2017; Damerval et al., 2019), this is the first study to quantify the total number of *CYCLOIDEA* paralogues in Proteaceae based on a complete genome sequence. Our results hence lend further support to the pattern of a single gene duplication in the stem lineage of Proteaceae that had so far emerged from Sanger and transcriptome sequencing.

**Figure 12.**
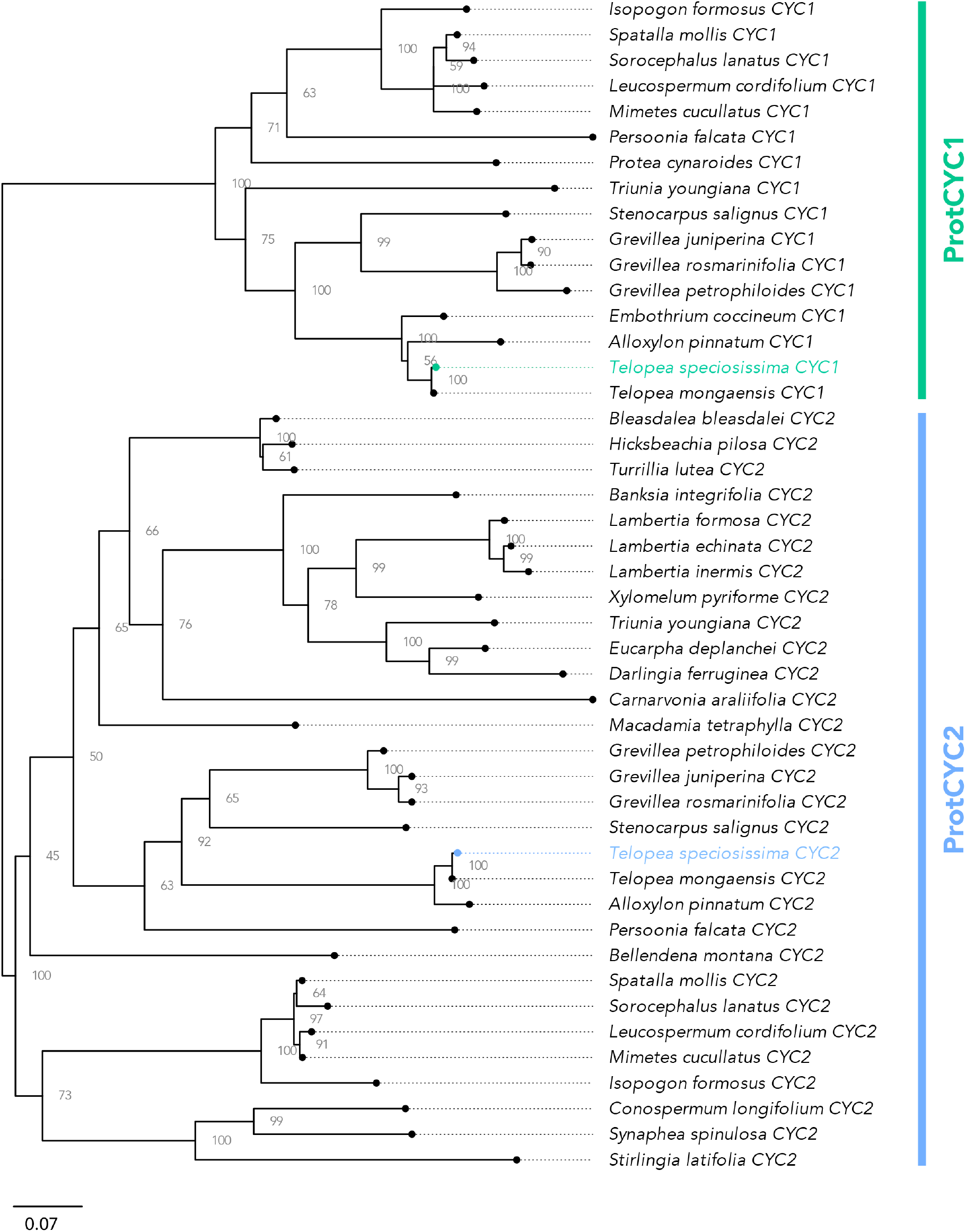
Phylogeny of *CYCLOIDEA* (*CYC*) proteins in Proteaceae, obtained from maximum-likelihood inference with IQ-TREE. Node numbers indicate bootstrap support expressed as percentage. Scale bar represents 0.07 nucleotide substitutions per site. Branches terminate at circles; dotted extensions are for labelling purporses only.

### A molecular resource for biodiversity genomics

The *T. speciosissima* reference genome will enable genome-scale research into Proteaceae evolution, at a wide range of scales. At shallower evolutionary scales, the *Telopea* genus contains five species that exhibit genetic variation consistent with a history of divergence and introgression, likely driven by climatic change (Rossetto et al., 2011, 2012). Recent studies highlight the power of genome-scale approaches for inferring demographic change and mechanistic forces that have influenced such clades, often making use of heterogenetity in patterns of variation across whole genomes (Choi et al., 2021; Soltis & Soltis, 2021). We expect the waratah genome to similarly facilitate studies that provide new insights about historical gene flow and selection, in changing environments.

## CONCLUSIONS

We present a high-quality annotated chromosome-level reference genome of *Telopea speciosissima* assembled from Oxford Nanopore long-reads, 10x Genomics Chromium linked-reads and Hi-C (823 Mb in length, N50 of 69.9 Mb and BUSCO completeness of 97.8 %): the first for a waratah, and only the second publicly available Proteaceae reference genome. We envisage these data will be a platform to underpin evolutionary genomics, gene discovery, breeding and the conservation of Proteaceae and the Australian flora.

## Supporting information

BUSCOMP full report

## ACKNOWLEDGEMENTS

We thank Stuart Allan for providing access to the sequenced plant and assistance with sample collection at Blue Mountains Botanic Garden and Carolyn Connelly for facilitating access to lab materials at the Royal Botanic Garden Sydney. We acknowledge Chris Jackson for advice on repeat annotation. We thank the members of UNSW Research Technology Services, particularly Duncan Smith, for help with software installation on the high-performance computing cluster Katana. We acknowledge Mabel Lum for assistance with the Bioplatforms Australia data portal. ONT and 10x sequencing were conducted at the Australian Genome Research Facility (AGRF). Hi-C library prep and sequencing was conducted at the Ramaciotti Centre for Genomics at the University of New South Wales.

## AUTHOR CONTRIBUTIONS

JGB coordinated the project. MR, MvdM, PL-I, HS, GB, JGB and RJE designed the study and funded the project. GB provided the samples. PL-I and J-YSY performed optimised DNA extraction protocols and performed extractions. SHC performed the genome assembly, scaffolding and annotation. RJE conceptualised and developed Diploidocus and DepthSizer. TGA and RJE performed the DepthSizer benchmarking analysis. RJE performed the copy number analysis and *CYC* phylogenetics. SHC, RJE and JGB wrote the manuscript. All authors edited and approved the final manuscript.

## DATA ACCESSIBILITY

The Tspe_v1 genome was deposited to NCBI under BioProject PRJNA712988 and BioSample along with the raw data (ONT, 10x and Hi-C) to SRA as SRR14018636, SRR14018635 and SRR14018634. The genome may be browsed via Apollo: https://edwapollo.babs.unsw.edu.au/apollo208/1468723/jbrowse/index.html. The NCBI Annotation Release 100 is available at https://ftp.ncbi.nlm.nih.gov/genomes/all/GCF/018/873/765/GCF_018873765.1_Tspe_v1 and the annotation is available for browsing in GDV: https://www.ncbi.nlm.nih.gov/genome/gdv/browser/?acc=GCF_018873765.1&context=genome.

Supplementary data, was deposited to Dryad (https://doi.org/10.5061/dryad.12jm63xzt) and contains files for tracks available on the Apollo genome browser (genome, gaps, mapped ONT and 10x reads and annotations) and the protein sequences from the GeMoMa genome annotation.

Data for species used for genome annotation are available at the following repositories:

*Macadamia integrifolia*

https://ftp.ncbi.nlm.nih.gov/genomes/all/GCA/013/358/625/GCA_013358625.1_SCU_Mint_v3/doi.org/10.25918/5e320fd1e5f06

*Arabidopsis thaliana*

https://ftp.ncbi.nlm.nih.gov/genomes/all/GCF/000/001/735/GCF_000001735.4_TAIR10.1/

*Rosa chinensis*

https://ftp.ncbi.nlm.nih.gov/genomes/all/GCA/002/994/745/GCA_002994745.2_RchiOBHm-V2/

*Nelumbo nucifera*

https://ftp.ncbi.nlm.nih.gov/genomes/all/GCF/000/365/185/GCF_000365185.1_Chinese_Lotus_1.1

