## Supplementary material for "Chromosome-level *de novo* genome assembly of *Telopea speciosissima* (New South Wales waratah) using long-reads, linked-reads and Hi-C": BUSCOMP full report: BUSCOMP_full_report_buscov3.html


### BUSCOMP Full Report

###### *buscov3-full BUSCOMP Analysis*

#### *2021-10-04*

### 1 BUSCOMP Run Summary

```
BUSCOMP V0.12.0: run Sun Oct  3 23:04:21 2021
```

See the run details appendix end of this document for details of the log file, commandline parameters and runtime BUSCOMP errors and warnings.

**NOTE:** To edit this document, open `buscov3-full.N3L20ID0U.full.Rmd` in RStudio, edit and re-knit to HTML.

#### 1.1 BUSCOMP Results Summary

Assemblies can be assessed on a number of criteria, but the main ones (in the absence of a reference “truth” genome) are either to judge contiguity or completeness. NG50 and LG50 values are based on a genome size of 982.0 Mb. If the `genomesize=X` parameter was not set (see command list in appendix), this will be based on the longest assembly (see sequence stats, below).

Of the 14 assemblies analysed (14 BUSCO; 14 fasta; 14 both), 14 genomes were rated as the “best” by at least one criterion:

- `Tspe_v1`: LG50Count, Missing.
- `Tspe_v0.1_necat`: Missing.
- `Tspe_v1.chr`: LG50Count.
- `Tspe_10xpseudohap2`: Missing.
- `Tspe_v0.8`: LG50Count, Missing.
- `Tspe_v0.1_flye`: Missing.
- `Tspe_v0.9`: NG50Length, LG50Count, MaxLength, Missing.
- `Tspe_v0.1_canu`: Missing.
- `Tspe_v0.2`: Missing.
- `Tspe_v0.3`: Complete, Missing.
- `Tspe_v0.4`: Complete, Missing, BUSCO, NoBUSCO.
- `Tspe_v0.5`: Complete, Missing.
- `Tspe_v0.6`: Complete, Missing.
- `Tspe_v0.7`: Complete, Missing.

Best assemblies by assembly contiguity critera:

- **NG50Length.** Longest NG50 contig/scaffold length (67,756,876 bp): `Tspe_v0.9`
- **LG50Count.** Smallest LG50 contig/scaffold count (7): `Tspe_v0.8`, `Tspe_v0.9`, `Tspe_v1`, `Tspe_v1.chr`
- **MaxLength.** Maximum contig/scaffold length (87,816,475 bp): `Tspe_v0.9`

Best assemblies by completeness critera:

- **Complete.** Most Complete (Single & Duplicated) BUSCOMP sequences (99.8 %): `Tspe_v0.3`, `Tspe_v0.4`, `Tspe_v0.5`, `Tspe_v0.6`, `Tspe_v0.7`
- **Missing.** Fewest Missing BUSCOMP sequences (0.0 %): `Tspe_10xpseudohap2`, `Tspe_v0.1_canu`, `Tspe_v0.1_flye`, `Tspe_v0.1_necat`, `Tspe_v0.2`, `Tspe_v0.3`, `Tspe_v0.4`, `Tspe_v0.5`, `Tspe_v0.6`, `Tspe_v0.7`, `Tspe_v0.8`, `Tspe_v0.9`, `Tspe_v1`
- **BUSCO.** Most Complete (Single & Duplicated) BUSCO sequences (92.6 %): `Tspe_v0.4`
- **NoBUSCO.** Fewest Missing BUSCO sequences (4.7 %): `Tspe_v0.4`

### 2 Genome Summary

The following genomes and BUSCO results were analysed by BUSCOMP:

- **Tspe\_10xpseudohap2**. [`BUSCO`|`Fasta`] /srv/scratch/z5239484/Telopea-Aug19/data/2021-04-05.TspeWorkflow/Tspe\_10xpseudohap2.fasta
- **Tspe\_v0.1\_canu**. [`BUSCO`|`Fasta`] /srv/scratch/z5239484/Telopea-Aug19/data/2021-04-05.TspeWorkflow/Tspe\_v0.1\_canu.fasta
- **Tspe\_v0.1\_flye**. [`BUSCO`|`Fasta`] /srv/scratch/z5239484/Telopea-Aug19/data/2021-04-05.TspeWorkflow/Tspe\_v0.1\_flye.fasta
- **Tspe\_v0.1\_necat**. [`BUSCO`|`Fasta`] /srv/scratch/z5239484/Telopea-Aug19/data/2021-04-05.TspeWorkflow/Tspe\_v0.1\_necat.fasta
- **Tspe\_v0.2**. [`BUSCO`|`Fasta`] /srv/scratch/z5239484/Telopea-Aug19/data/2021-04-05.TspeWorkflow/Tspe\_v0.2.fasta
- **Tspe\_v0.3**. [`BUSCO`|`Fasta`] /srv/scratch/z5239484/Telopea-Aug19/data/2021-04-05.TspeWorkflow/Tspe\_v0.3.fasta
- **Tspe\_v0.4**. [`BUSCO`|`Fasta`] /srv/scratch/z5239484/Telopea-Aug19/data/2021-04-05.TspeWorkflow/Tspe\_v0.4.fasta
- **Tspe\_v0.5**. [`BUSCO`|`Fasta`] /srv/scratch/z5239484/Telopea-Aug19/data/2021-04-05.TspeWorkflow/Tspe\_v0.5.fasta
- **Tspe\_v0.6**. [`BUSCO`|`Fasta`] /srv/scratch/z5239484/Telopea-Aug19/data/2021-04-05.TspeWorkflow/Tspe\_v0.6.fasta
- **Tspe\_v0.7**. [`BUSCO`|`Fasta`] /srv/scratch/z5239484/Telopea-Aug19/data/2021-04-05.TspeWorkflow/Tspe\_v0.7.fasta
- **Tspe\_v0.8**. [`BUSCO`|`Fasta`] /srv/scratch/z5239484/Telopea-Aug19/data/2021-04-05.TspeWorkflow/Tspe\_v0.8.fasta
- **Tspe\_v0.9**. [`BUSCO`|`Fasta`] /srv/scratch/z5239484/Telopea-Aug19/data/2021-04-05.TspeWorkflow/Tspe\_v0.9.fasta
- **Tspe\_v1**. [`BUSCO`|`Fasta`] /srv/scratch/z5239484/Telopea-Aug19/data/2021-04-05.TspeWorkflow/Tspe\_v1.fasta
- **Tspe\_v1.chr**. [`BUSCO`|`Fasta`] /srv/scratch/z5239484/Telopea-Aug19/data/2021-04-05.TspeWorkflow/Tspe\_v1.chr.fasta

Details of the directories and files are below:

Genomes with a **Directory** listed had BUSCO results available. If **Sequences** is `True`, these would be have been compiled to generate the BUSCOMP sequence set (unless `buscompseq=F`, or alternative sequences were provided with `buscofas=FASFILE`). Genomes with a **Fasta** listed had sequence data available for BUSCOMP searches.

#### 2.1 Genome statistics

The following genome statistics were also calculated by `RJE_SeqList` for each genome (table, below):

- **SeqNum**: The total number of scaffolds/contigs in the assembly.
- **TotLength**: The total combined length of scaffolds/contigs in the assembly.
- **MinLength**: The length of the shortest scaffold/contig in the assembly.
- **MaxLength**: The length of the longest scaffold/contig in the assembly.
- **MeanLength**: The mean length of scaffolds/contigs in the assembly.
- **MedLength**: The median length of scaffolds/contigs in the assembly.
- **N50Length**: At least half of the assembly is contained on scaffolds/contigs of this length or greater.
- **L50Count**: The smallest number scaffolds/contigs needed to cover half the the assembly.
- **CtgNum**: Number of contigs (`SeqNum`+`GapCount`).
- **N50Ctg**: At least half of the assembly is contained on contigs of this length or greater.
- **L50Ctg**: The smallest number contigs needed to cover half the the assembly.
- **NG50Length**: At least half of the genome is contained on scaffolds/contigs of this length or greater. This is based on `genomesize=X`. If no genome size is given, it will be relative to the biggest assembly.
- **LG50Count**: The smallest number scaffolds/contigs needed to cover half the the genome. This is based on `genomesize=X`. If no genome size is given, it will be relative to the biggest assembly.
- **GapLength**: The total number of undefined “gap” (`N`) nucleotides in the assembly.
- **GapCount**: The total number of undefined “gap” (`N`) regions in the assembly.
- **GC**: The %GC content of the assembly.

| Genome | Description | SeqNum | TotLength | MinLength | MaxLength | MeanLength | MedLength | N50Length | L50Count | CtgNum | N50Ctg | L50Ctg | NG50Length | LG50Count | GapLength | GapCount | GC |
| --- | --- | --- | --- | --- | --- | --- | --- | --- | --- | --- | --- | --- | --- | --- | --- | --- | --- |
| Tspe\_10xpseudohap2 | /srv/scratch/z5239484/Telopea-Aug19/data/2021-04-05.TspeWorkflow/Tspe\_10xpseudohap2.fasta | 27610 | 850975689 | 1000 | 7917366 | 30821.29 | 2990 | 874466 | 247 | 43951 | 72725 | 3268 | 694813 | 331 | 18076790 | 16341 | 40.10 |
| Tspe\_v0.1\_canu | /srv/scratch/z5239484/Telopea-Aug19/data/2021-04-05.TspeWorkflow/Tspe\_v0.1\_canu.fasta | 3983 | 981953849 | 1138 | 12971499 | 246536.24 | 58209 | 1848137 | 132 | 3983 | 1848137 | 132 | 1848137 | 132 | 0 | 0 | 39.95 |
| Tspe\_v0.1\_flye | /srv/scratch/z5239484/Telopea-Aug19/data/2021-04-05.TspeWorkflow/Tspe\_v0.1\_flye.fasta | 2445 | 857703641 | 505 | 12987791 | 350799.04 | 31428 | 2271126 | 94 | 2484 | 2199532 | 101 | 1823111 | 125 | 3900 | 39 | 40.47 |
| Tspe\_v0.1\_necat | /srv/scratch/z5239484/Telopea-Aug19/data/2021-04-05.TspeWorkflow/Tspe\_v0.1\_necat.fasta | 365 | 842143239 | 563 | 37347493 | 2307241.75 | 155267 | 10701597 | 24 | 365 | 10701597 | 24 | 9180758 | 31 | 0 | 0 | 40.15 |
| Tspe\_v0.2 | /srv/scratch/z5239484/Telopea-Aug19/data/2021-04-05.TspeWorkflow/Tspe\_v0.2.fasta | 210 | 827757806 | 1773 | 37347493 | 3941703.84 | 1153652 | 10955845 | 23 | 210 | 10955845 | 23 | 9180758 | 31 | 0 | 0 | 40.13 |
| Tspe\_v0.3 | /srv/scratch/z5239484/Telopea-Aug19/data/2021-04-05.TspeWorkflow/Tspe\_v0.3.fasta | 209 | 834569695 | 3516 | 37634595 | 3993156.44 | 1227984 | 11046571 | 23 | 209 | 11046571 | 23 | 9263258 | 31 | 0 | 0 | 40.27 |
| Tspe\_v0.4 | /srv/scratch/z5239484/Telopea-Aug19/data/2021-04-05.TspeWorkflow/Tspe\_v0.4.fasta | 209 | 835074331 | 3532 | 37649015 | 3995570.96 | 1228174 | 11052047 | 23 | 209 | 11052047 | 23 | 9267273 | 31 | 0 | 0 | 40.25 |
| Tspe\_v0.5 | /srv/scratch/z5239484/Telopea-Aug19/data/2021-04-05.TspeWorkflow/Tspe\_v0.5.fasta | 138 | 835372011 | 3532 | 49795992 | 6053420.37 | 2458095 | 16474627 | 15 | 205 | 11052047 | 23 | 14087303 | 20 | 297663 | 67 | 40.25 |
| Tspe\_v0.6 | /srv/scratch/z5239484/Telopea-Aug19/data/2021-04-05.TspeWorkflow/Tspe\_v0.6.fasta | 138 | 835438898 | 3484 | 49793446 | 6053905.06 | 2461216 | 16468668 | 15 | 159 | 14704285 | 17 | 14096844 | 20 | 256618 | 21 | 40.27 |
| Tspe\_v0.7 | /srv/scratch/z5239484/Telopea-Aug19/data/2021-04-05.TspeWorkflow/Tspe\_v0.7.fasta | 128 | 833847971 | 3484 | 49793446 | 6514437.27 | 2967336 | 16468668 | 15 | 148 | 14704285 | 17 | 14096844 | 20 | 244118 | 20 | 40.27 |
| Tspe\_v0.8 | /srv/scratch/z5239484/Telopea-Aug19/data/2021-04-05.TspeWorkflow/Tspe\_v0.8.fasta | 2357 | 833952765 | 5 | 87707951 | 353819.59 | 11506 | 68901929 | 6 | 3575 | 1773000 | 131 | 67726057 | 7 | 982622 | 1218 | 40.03 |
| Tspe\_v0.9 | /srv/scratch/z5239484/Telopea-Aug19/data/2021-04-05.TspeWorkflow/Tspe\_v0.9.fasta | 1399 | 825066408 | 974 | 87816475 | 589754.40 | 18113 | 69015533 | 6 | 1566 | 12206888 | 21 | 67756876 | 7 | 316262 | 167 | 40.12 |
| Tspe\_v1 | /srv/scratch/z5239484/Telopea-Aug19/data/2021-04-05.TspeWorkflow/Tspe\_v1.fasta | 1289 | 823061212 | 974 | 87811904 | 638526.93 | 19505 | 69013595 | 6 | 1452 | 12206888 | 21 | 67754596 | 7 | 16300 | 163 | 40.11 |
| Tspe\_v1.chr | /srv/scratch/z5239484/Telopea-Aug19/data/2021-04-05.TspeWorkflow/Tspe\_v1.chr.fasta | 11 | 774398234 | 57317785 | 87811904 | 70399839.45 | 69013595 | 69013595 | 6 | 158 | 13444441 | 19 | 67754596 | 7 | 14700 | 147 | 40.06 |

**NOTE:** `NG50Length` and `LG50Count` statistics use `genomesize=X` or the biggest assembly loaded (981.95 Mb). If BUSCOMP has been run more than once on the same data (*e.g.* to update descriptions or sorting), please make sure that a consistent genome size is used, or these values may be wrong. If in doubt, run with `force=T` and force regeneration of statistics.

#### 2.2 Genome coverage assessment plots

In general, a good assembly will be approx. the same size as the genome and in as few pieces as possible. Any assembly smaller than the predicted genome size is clearly missing coverage. Assemblies bigger than the genome size might still be missing chunks of the genome if redundancy/duplication is a problem. In the following plot, the grey line marks the given genome size of 982.0 Mb.

A better indicator of the overall coverage of the genome is the number of `Missing` BUSCO genes. As BUSCO is highly dependent on the accuracy of the sequence and the gene models it makes, the `Missing` BUSCOMP ratings arguably give a more consistent proxy for genome completeness. NOTE: this says nothing about the fragmentation or completeness of the genes themselves.

#### 2.3 Genome contiguity assessment plots

In general, a good assembly will be in fewer, bigger pieces. This is approximated using NG50 and LG50, which are the min. length and number of contigs/scaffolds required to cover at least half the genome. These stats use the given genome size of 982.0 Mb.

**NOTE:** To modify these plots and tables, edit the `*.genomes.tdt` and `*.NxLxxIDxx.rdata.tdt` files and re-knit the `*.NxLxxIDxx.Rmd` file.

### 3 BUSCO Ratings

Compiled BUSCO results for 14 assemblies and 4 groups have been saved in `buscov3-full.genomes.tdt`. BUSCO ratings are defined (quoting from the BUSCO v3 User Guide as:

- `Complete`: Single-copy hits where “BUSCO matches have scored within the expected range of scores and within the expected range of length alignments to the BUSCO profile.”
- `Duplicated`: As `Complete` but 2+ copies.
- `Fragmented`: “BUSCO matches … within the range of scores but not within the range of length alignments to the BUSCO profile.”
- `Missing`: “Either no significant matches at all, or the BUSCO matches scored below the range of scores for the BUSCO profile.”

| Genome | N | Complete | Single | Duplicated | Fragmented | Missing |
| --- | --- | --- | --- | --- | --- | --- |
| Tspe\_10xpseudohap2 | 1440 | 1323 | 1170 | 153 | 44 | 73 |
| Tspe\_v0.1\_canu | 1440 | 1129 | 988 | 141 | 75 | 236 |
| Tspe\_v0.1\_flye | 1440 | 1167 | 1079 | 88 | 60 | 213 |
| Tspe\_v0.1\_necat | 1440 | 1169 | 1083 | 86 | 62 | 209 |
| Tspe\_v0.2 | 1440 | 1167 | 1083 | 84 | 62 | 211 |
| Tspe\_v0.3 | 1440 | 1308 | 1177 | 131 | 40 | 92 |
| Tspe\_v0.4 | 1440 | 1333 | 1178 | 155 | 39 | 68 |
| Tspe\_v0.5 | 1440 | 1328 | 1169 | 159 | 43 | 69 |
| Tspe\_v0.6 | 1440 | 1304 | 1171 | 133 | 42 | 94 |
| Tspe\_v0.7 | 1440 | 1303 | 1170 | 133 | 43 | 94 |
| Tspe\_v0.8 | 1440 | 1231 | 1142 | 89 | 55 | 154 |
| Tspe\_v0.9 | 1440 | 1314 | 1174 | 140 | 41 | 85 |
| Tspe\_v1 | 1440 | 1314 | 1174 | 140 | 42 | 84 |
| Tspe\_v1.chr | 1440 | 1304 | 1172 | 132 | 38 | 98 |
| Tspe\_v1-genome | 1440 | 1317 | 1187 | 130 | 42 | 81 |
| Tspe\_v1-full | 1440 | 1317 | 1187 | 130 | 42 | 81 |
| BUSCOv3 | 1440 | 1391 | 1361 | 30 | 12 | 37 |
| BUSCOMP | 1440 | 1399 | 1386 | 13 | 10 | 31 |

#### 3.1 Genome Groups

`BUSCOMP` compiled the following groups of genomes (where BUSCO data was loaded), keeping the “best” rating for each BUSCO gene across the group:

- **Tspe\_v1-genome**: `Tspe_v1` `Tspe_v1.chr`
- **Tspe\_v1-full**: `Tspe_v1` `Tspe_v1.chr`
- **BUSCOv3**: `Tspe_v0.1_necat` `Tspe_v0.2` `Tspe_v0.3` `Tspe_v0.4` `Tspe_v0.5` `Tspe_v0.6` `Tspe_v0.7` `Tspe_v0.8` `Tspe_v0.9` `Tspe_v1` `Tspe_v1.chr`
- **BUSCOMP**: `Tspe_10xpseudohap2` `Tspe_v0.1_canu` `Tspe_v0.1_flye` `Tspe_v0.1_necat` `Tspe_v0.2` `Tspe_v0.3` `Tspe_v0.4` `Tspe_v0.5` `Tspe_v0.6` `Tspe_v0.7` `Tspe_v0.8` `Tspe_v0.9` `Tspe_v1` `Tspe_v1.chr`

#### 3.2 BUSCO Summary

```
Tspe_10xpseudohap2 BUSCO Results:
        C:91.9%[S:81.2%,D:10.6%],F:3.1%,M:5.1%,n:1440

Tspe_v0.1_canu BUSCO Results:
        C:78.4%[S:68.6%,D:9.8%],F:5.2%,M:16.4%,n:1440

Tspe_v0.1_flye BUSCO Results:
        C:81.0%[S:74.9%,D:6.1%],F:4.2%,M:14.8%,n:1440

Tspe_v0.1_necat BUSCO Results:
        C:81.2%[S:75.2%,D:6.0%],F:4.3%,M:14.5%,n:1440

Tspe_v0.2 BUSCO Results:
        C:81.0%[S:75.2%,D:5.8%],F:4.3%,M:14.7%,n:1440

Tspe_v0.3 BUSCO Results:
        C:90.8%[S:81.7%,D:9.1%],F:2.8%,M:6.4%,n:1440

Tspe_v0.4 BUSCO Results:
        C:92.6%[S:81.8%,D:10.8%],F:2.7%,M:4.7%,n:1440

Tspe_v0.5 BUSCO Results:
        C:92.2%[S:81.2%,D:11.0%],F:3.0%,M:4.8%,n:1440

Tspe_v0.6 BUSCO Results:
        C:90.6%[S:81.3%,D:9.2%],F:2.9%,M:6.5%,n:1440

Tspe_v0.7 BUSCO Results:
        C:90.5%[S:81.2%,D:9.2%],F:3.0%,M:6.5%,n:1440

Tspe_v0.8 BUSCO Results:
        C:85.5%[S:79.3%,D:6.2%],F:3.8%,M:10.7%,n:1440

Tspe_v0.9 BUSCO Results:
        C:91.2%[S:81.5%,D:9.7%],F:2.8%,M:5.9%,n:1440

Tspe_v1 BUSCO Results:
        C:91.2%[S:81.5%,D:9.7%],F:2.9%,M:5.8%,n:1440

Tspe_v1.chr BUSCO Results:
        C:90.6%[S:81.4%,D:9.2%],F:2.6%,M:6.8%,n:1440

Tspe_v1-genome BUSCO Results:
        C:91.5%[S:82.4%,D:9.0%],F:2.9%,M:5.6%,n:1440

Tspe_v1-full BUSCO Results:
        C:91.5%[S:82.4%,D:9.0%],F:2.9%,M:5.6%,n:1440

BUSCOv3 BUSCO Results:
        C:96.6%[S:94.5%,D:2.1%],F:0.8%,M:2.6%,n:1440

BUSCOMP BUSCO Results:
        C:97.2%[S:96.2%,D:0.9%],F:0.7%,M:2.2%,n:1440
```

#### 3.3 BUSCO Gene Details

Full BUSCO results with ratings for each gene have been compiled in `buscov3-full.busco.tdt`:

#### 3.4 Genome Group BUSCO charts

### 4 BUSCOMP Ratings

The best complete BUSCO hit results (based on `Score` and `Length`) have been compiled in `buscov3-full.buscoseq.tdt`. The `Genome` field indicates the assembly with the best hit, which is followed by details of that hit (`Contig`, `Start`, `End`, `Score`, `Length`). BUSCOMP ratings for each assembly are then given in subsequent fields:

```
* `Identical`: 100% coverage and 100% identity in at least one contig/scaffold.
* `Complete`: 95%+ Coverage in a single contig/scaffold. (Note: accuracy/identity is not considered.)
* `Duplicated`: 95%+ Coverage in 2+ contigs/scaffolds.
* `Fragmented`: 95%+ combined coverage but not in any single contig/scaffold.
* `Partial`: 40-95% combined coverage.
* `Ghost`: Hits meeting local cutoff but <40% combined coverage.
* `Missing`: No hits meeting local cutoff.
```

#### 4.1 BUSCOSeq Rating Summary

BUSCOMP ratings (see above) are compiled to summary statistics in `buscov3-full.N3L20ID0U.ratings.tdt`. Note that `Identical` ratings in this table will also be rated as `Complete`, which in turn are `Single` or `Duplicated`. Percentage summaries are plotted below, along with a BUSCO-style one-line summary per assembly/group.

**NOTE:** Group summaries do not include `Identical` ratings.

| X. | Genome | N | Identical | Complete | Single | Duplicated | Fragmented | Partial | Ghost | Missing |
| --- | --- | --- | --- | --- | --- | --- | --- | --- | --- | --- |
| 1 | Tspe\_10xpseudohap2 | 1386 | 615 | 1344 | 1194 | 150 | 35 | 5 | 2 | 0 |
| 2 | Tspe\_v0.1\_canu | 1386 | 77 | 1382 | 1094 | 288 | 2 | 2 | 0 | 0 |
| 3 | Tspe\_v0.1\_flye | 1386 | 461 | 1371 | 1224 | 147 | 11 | 4 | 0 | 0 |
| 4 | Tspe\_v0.1\_necat | 1386 | 141 | 1381 | 1234 | 147 | 1 | 4 | 0 | 0 |
| 5 | Tspe\_v0.2 | 1386 | 140 | 1381 | 1237 | 144 | 1 | 4 | 0 | 0 |
| 6 | Tspe\_v0.3 | 1386 | 487 | 1383 | 1241 | 142 | 0 | 3 | 0 | 0 |
| 7 | Tspe\_v0.4 | 1386 | 580 | 1383 | 1241 | 142 | 0 | 3 | 0 | 0 |
| 8 | Tspe\_v0.5 | 1386 | 580 | 1383 | 1240 | 143 | 0 | 3 | 0 | 0 |
| 9 | Tspe\_v0.6 | 1386 | 479 | 1383 | 1240 | 143 | 0 | 3 | 0 | 0 |
| 10 | Tspe\_v0.7 | 1386 | 479 | 1383 | 1240 | 143 | 0 | 3 | 0 | 0 |
| 11 | Tspe\_v0.8 | 1386 | 220 | 1353 | 1230 | 123 | 21 | 6 | 6 | 0 |
| 12 | Tspe\_v0.9 | 1386 | 546 | 1355 | 1235 | 120 | 18 | 5 | 8 | 0 |
| 13 | Tspe\_v1 | 1386 | 546 | 1355 | 1235 | 120 | 18 | 5 | 8 | 0 |
| 14 | Tspe\_v1.chr | 1386 | 543 | 1346 | 1231 | 115 | 3 | 14 | 19 | 4 |
| 15 | Tspe\_v1-genome | 1386 | 0 | 1355 | 1239 | 116 | 18 | 5 | 8 | 0 |
| 16 | Tspe\_v1-full | 1386 | 0 | 1355 | 1239 | 116 | 18 | 5 | 8 | 0 |
| 17 | BUSCOv3 | 1386 | 0 | 1384 | 1271 | 113 | 0 | 2 | 0 | 0 |
| 18 | BUSCOMP | 1386 | 0 | 1384 | 1280 | 104 | 0 | 2 | 0 | 0 |

```
BUSCOMP BUSCOMP Results [1399 (97.15%) Complete BUSCOs; 0 (0.00%) BUSCOMP Seqs]:
        C:99.9%[S:92.4%,D:7.5%],F:0.0%,P:0.1%,G:0.0%,M:0.0%,n:1386

BUSCOv3 BUSCOMP Results [1391 (96.60%) Complete BUSCOs; 0 (0.00%) BUSCOMP Seqs]:
        C:99.9%[S:91.7%,D:8.2%],F:0.0%,P:0.1%,G:0.0%,M:0.0%,n:1386

Tspe_10xpseudohap2 BUSCOMP Results [1323 (91.88%) Complete BUSCOs; 588 (42.42%) BUSCOMP Seqs]:
        C:97.0%[S:86.1%,D:10.8%,I:44.4%],F:2.5%,P:0.4%,G:0.1%,M:0.0%,n:1386

Tspe_v0.1_canu BUSCOMP Results [1129 (78.40%) Complete BUSCOs; 63 (4.55%) BUSCOMP Seqs]:
        C:99.7%[S:78.9%,D:20.8%,I:5.6%],F:0.1%,P:0.1%,G:0.0%,M:0.0%,n:1386

Tspe_v0.1_flye BUSCOMP Results [1167 (81.04%) Complete BUSCOs; 76 (5.48%) BUSCOMP Seqs]:
        C:98.9%[S:88.3%,D:10.6%,I:33.3%],F:0.8%,P:0.3%,G:0.0%,M:0.0%,n:1386

Tspe_v0.1_necat BUSCOMP Results [1169 (81.18%) Complete BUSCOs; 67 (4.83%) BUSCOMP Seqs]:
        C:99.6%[S:89.0%,D:10.6%,I:10.2%],F:0.1%,P:0.3%,G:0.0%,M:0.0%,n:1386

Tspe_v0.2 BUSCOMP Results [1167 (81.04%) Complete BUSCOs; 6 (0.43%) BUSCOMP Seqs]:
        C:99.6%[S:89.2%,D:10.4%,I:10.1%],F:0.1%,P:0.3%,G:0.0%,M:0.0%,n:1386

Tspe_v0.3 BUSCOMP Results [1308 (90.83%) Complete BUSCOs; 159 (11.47%) BUSCOMP Seqs]:
        C:99.8%[S:89.5%,D:10.2%,I:35.1%],F:0.0%,P:0.2%,G:0.0%,M:0.0%,n:1386

Tspe_v0.4 BUSCOMP Results [1333 (92.57%) Complete BUSCOs; 93 (6.71%) BUSCOMP Seqs]:
        C:99.8%[S:89.5%,D:10.2%,I:41.8%],F:0.0%,P:0.2%,G:0.0%,M:0.0%,n:1386

Tspe_v0.5 BUSCOMP Results [1328 (92.22%) Complete BUSCOs; 15 (1.08%) BUSCOMP Seqs]:
        C:99.8%[S:89.5%,D:10.3%,I:41.8%],F:0.0%,P:0.2%,G:0.0%,M:0.0%,n:1386

Tspe_v0.6 BUSCOMP Results [1304 (90.56%) Complete BUSCOs; 53 (3.82%) BUSCOMP Seqs]:
        C:99.8%[S:89.5%,D:10.3%,I:34.6%],F:0.0%,P:0.2%,G:0.0%,M:0.0%,n:1386

Tspe_v0.7 BUSCOMP Results [1303 (90.49%) Complete BUSCOs; 6 (0.43%) BUSCOMP Seqs]:
        C:99.8%[S:89.5%,D:10.3%,I:34.6%],F:0.0%,P:0.2%,G:0.0%,M:0.0%,n:1386

Tspe_v0.8 BUSCOMP Results [1231 (85.49%) Complete BUSCOs; 79 (5.70%) BUSCOMP Seqs]:
        C:97.6%[S:88.7%,D:8.9%,I:15.9%],F:1.5%,P:0.4%,G:0.4%,M:0.0%,n:1386

Tspe_v0.9 BUSCOMP Results [1314 (91.25%) Complete BUSCOs; 162 (11.69%) BUSCOMP Seqs]:
        C:97.8%[S:89.1%,D:8.7%,I:39.4%],F:1.3%,P:0.4%,G:0.6%,M:0.0%,n:1386

Tspe_v1 BUSCOMP Results [1314 (91.25%) Complete BUSCOs; 1 (0.07%) BUSCOMP Seqs]:
        C:97.8%[S:89.1%,D:8.7%,I:39.4%],F:1.3%,P:0.4%,G:0.6%,M:0.0%,n:1386

Tspe_v1-full BUSCOMP Results [1317 (91.46%) Complete BUSCOs; 0 (0.00%) BUSCOMP Seqs]:
        C:97.8%[S:89.4%,D:8.4%],F:1.3%,P:0.4%,G:0.6%,M:0.0%,n:1386

Tspe_v1-genome BUSCOMP Results [1317 (91.46%) Complete BUSCOs; 0 (0.00%) BUSCOMP Seqs]:
        C:97.8%[S:89.4%,D:8.4%],F:1.3%,P:0.4%,G:0.6%,M:0.0%,n:1386

Tspe_v1.chr BUSCOMP Results [1304 (90.56%) Complete BUSCOs; 18 (1.30%) BUSCOMP Seqs]:
        C:97.1%[S:88.8%,D:8.3%,I:39.2%],F:0.2%,P:1.0%,G:1.4%,M:0.3%,n:1386
```

#### 4.2 BUSCOSeq Full Results Table

Full BUSCOMP results with ratings for each gene in every assembly and group have been compiled in `buscov3-full.N3L20ID0U.buscomp.tdt`:

#### 4.3 Genome Group BUSCOMP charts

### 5 BUSCO and BUSCOMP Comparisons

#### 5.1 BUSCO to BUSCOMP Rating Changes

Ratings changes from BUSCO to BUSCOMP (where `NULL` ratings indicate no BUSCOMP sequence):

| BUSCO | BUSCOMP | Tspe\_10xpseudohap2 | Tspe\_v0.1\_canu | Tspe\_v0.1\_flye | Tspe\_v0.1\_necat | Tspe\_v0.2 | Tspe\_v0.3 | Tspe\_v0.4 | Tspe\_v0.5 | Tspe\_v0.6 | Tspe\_v0.7 | Tspe\_v0.8 | Tspe\_v0.9 | Tspe\_v1 | Tspe\_v1.chr | TOTAL |
| --- | --- | --- | --- | --- | --- | --- | --- | --- | --- | --- | --- | --- | --- | --- | --- | --- |
| Complete | Complete | 1073 | 852 | 973 | 992 | 993 | 1093 | 1110 | 1102 | 1088 | 1086 | 1042 | 1100 | 1100 | 1102 | 14706 |
| Complete | Duplicated | 75 | 133 | 96 | 88 | 87 | 84 | 67 | 66 | 82 | 83 | 80 | 58 | 58 | 58 | 1115 |
| Complete | Fragmented | 17 | 1 | 7 | 1 | 1 | 0 | 0 | 0 | 0 | 0 | 12 | 8 | 8 | 1 | 56 |
| Complete | Ghost | 0 | 0 | 0 | 0 | 0 | 0 | 0 | 0 | 0 | 0 | 4 | 3 | 3 | 3 | 13 |
| Complete | Partial | 5 | 2 | 3 | 2 | 2 | 0 | 1 | 1 | 1 | 1 | 4 | 5 | 5 | 8 | 40 |
| Duplicated | Complete | 63 | 36 | 43 | 36 | 34 | 64 | 69 | 71 | 65 | 66 | 44 | 66 | 66 | 61 | 784 |
| Duplicated | Duplicated | 73 | 93 | 32 | 38 | 38 | 52 | 72 | 74 | 54 | 53 | 30 | 59 | 59 | 56 | 783 |
| Duplicated | Fragmented | 3 | 0 | 0 | 0 | 0 | 0 | 0 | 0 | 0 | 0 | 1 | 2 | 2 | 1 | 9 |
| Duplicated | Ghost | 1 | 0 | 0 | 0 | 0 | 0 | 0 | 0 | 0 | 0 | 0 | 0 | 0 | 0 | 1 |
| Duplicated | NULL | 13 | 12 | 12 | 12 | 12 | 13 | 13 | 13 | 13 | 13 | 13 | 13 | 13 | 13 | 178 |
| Duplicated | Partial | 0 | 0 | 1 | 0 | 0 | 2 | 1 | 1 | 1 | 1 | 1 | 0 | 0 | 1 | 9 |
| Fragmented | Complete | 25 | 53 | 47 | 51 | 51 | 28 | 28 | 32 | 29 | 29 | 39 | 23 | 24 | 25 | 484 |
| Fragmented | Duplicated | 2 | 14 | 5 | 3 | 3 | 2 | 2 | 2 | 4 | 4 | 4 | 3 | 3 | 1 | 52 |
| Fragmented | Fragmented | 8 | 0 | 1 | 0 | 0 | 0 | 0 | 0 | 0 | 0 | 4 | 4 | 4 | 1 | 22 |
| Fragmented | Ghost | 0 | 0 | 0 | 0 | 0 | 0 | 0 | 0 | 0 | 0 | 0 | 1 | 1 | 1 | 3 |
| Fragmented | NULL | 9 | 8 | 7 | 8 | 8 | 9 | 9 | 9 | 9 | 10 | 8 | 10 | 10 | 10 | 124 |
| Fragmented | Partial | 0 | 0 | 0 | 0 | 0 | 1 | 0 | 0 | 0 | 0 | 0 | 0 | 0 | 0 | 1 |
| Missing | Complete | 33 | 153 | 161 | 155 | 159 | 56 | 34 | 35 | 58 | 59 | 105 | 46 | 45 | 43 | 1142 |
| Missing | Duplicated | 0 | 48 | 14 | 18 | 16 | 4 | 1 | 1 | 3 | 3 | 9 | 0 | 0 | 0 | 117 |
| Missing | Fragmented | 7 | 1 | 3 | 0 | 0 | 0 | 0 | 0 | 0 | 0 | 4 | 4 | 4 | 0 | 23 |
| Missing | Ghost | 1 | 0 | 0 | 0 | 0 | 0 | 0 | 0 | 0 | 0 | 2 | 4 | 4 | 15 | 26 |
| Missing | Missing | 0 | 0 | 0 | 0 | 0 | 0 | 0 | 0 | 0 | 0 | 0 | 0 | 0 | 4 | 4 |
| Missing | NULL | 32 | 34 | 35 | 34 | 34 | 32 | 32 | 32 | 32 | 31 | 33 | 31 | 31 | 31 | 454 |
| Missing | Partial | 0 | 0 | 0 | 2 | 2 | 0 | 1 | 1 | 1 | 1 | 1 | 0 | 0 | 5 | 14 |

Full table of Ratings changes from by gene:

**C**omplete, **D**uplicated, **F**ragmented, **P**artial, **G**host, **M**issing, **N**ULL (no BUSCOMP sequence)

##### 5.1.1 BUSCOMP Gain test

There is a risk that performing a low stringency search will identify homologues or pseudogenes of the desired BUSCO gene in error. If there is a second copy of a gene in the genome that is detectable by the search then we would expect the same genes that go from `Missing` to `Complete` in some genomes to go from `Single` to `Duplicated` in others.

To test this, data is reduced for each pair of genomes to BUSCO-BUSCOMP rating pairs of:

- `Single`-`Single`
- `Single`-`Duplicated`
- `Missing`-`Missing`
- `Missing`-`Single`

This is then converted in to `G`ain ratings (`Single`-`Duplicated` & `Missing`-`Single`) or `N`o Gain ratings (`Single`-`Single` & `Missing`-`Missing`). The `Single`-`Duplicated` shift in one genome is then used to set the expected `Missing`-`Single` shift in the other, and assess the probability of observing the `Missing`-`Single` shift using a cumulative binomial distribution, where:

- `k` is the number of observed `GG` pairs (`Single`-`Duplicated` *and* `Missing`-`Single`)
- `n` is the number of `Missing`-`Single` `G`ains in the focal genome (`NG`+`GG`)
- `p` is the proportion of `Single`-`Duplicated` `G`ains in the background genome (`GN`+`GG` / (`GN`+`GG`+`NN`+`NG`))
- `pB` is the probability of observing `k+` `Missing`-`Single` gains, given `p` and `n`

This is output to `*.gain.tdt`, where each row is a Genome and each field gives the probability of the row genome’s `Missing`-`Single` gains, given the column genome’s `Single`-`Duplicated` gains:

| Genome | Tspe\_10xpseudohap2 | Tspe\_v0.1\_canu | Tspe\_v0.1\_flye | Tspe\_v0.1\_necat | Tspe\_v0.2 | Tspe\_v0.3 | Tspe\_v0.4 | Tspe\_v0.5 | Tspe\_v0.6 | Tspe\_v0.7 | Tspe\_v0.8 | Tspe\_v0.9 | Tspe\_v1 | Tspe\_v1.chr |
| --- | --- | --- | --- | --- | --- | --- | --- | --- | --- | --- | --- | --- | --- | --- |
| Tspe\_10xpseudohap2 | 1.000 | 0.608 | 0.644 | 0.642 | 0.642 | 0.646 | 0.648 | 0.647 | 0.645 | 0.645 | 1 | 1.000 | 1.000 | 1.000 |
| Tspe\_v0.1\_canu | 1.000 | 1.000 | 1.000 | 1.000 | 1.000 | 1.000 | 1.000 | 1.000 | 0.634 | 0.634 | 1 | 1.000 | 1.000 | 1.000 |
| Tspe\_v0.1\_flye | 0.633 | 0.583 | 1.000 | 1.000 | 1.000 | 1.000 | 1.000 | 1.000 | 0.634 | 0.634 | 1 | 0.633 | 0.633 | 0.634 |
| Tspe\_v0.1\_necat | 0.596 | 0.602 | 0.635 | 1.000 | 1.000 | 1.000 | 1.000 | 1.000 | 1.000 | 1.000 | 1 | 0.634 | 0.634 | 0.634 |
| Tspe\_v0.2 | 0.596 | 0.602 | 0.635 | 1.000 | 1.000 | 1.000 | 1.000 | 1.000 | 1.000 | 1.000 | 1 | 0.634 | 0.634 | 0.634 |
| Tspe\_v0.3 | 1.000 | 0.607 | 1.000 | 1.000 | 1.000 | 1.000 | 1.000 | 1.000 | 1.000 | 1.000 | 1 | 1.000 | 1.000 | 1.000 |
| Tspe\_v0.4 | 1.000 | 0.642 | 1.000 | 1.000 | 1.000 | 1.000 | 1.000 | 1.000 | 1.000 | 1.000 | 1 | 1.000 | 1.000 | 1.000 |
| Tspe\_v0.5 | 1.000 | 0.641 | 1.000 | 1.000 | 1.000 | 1.000 | 1.000 | 1.000 | 1.000 | 1.000 | 1 | 1.000 | 1.000 | 1.000 |
| Tspe\_v0.6 | 1.000 | 0.590 | 1.000 | 1.000 | 1.000 | 1.000 | 1.000 | 1.000 | 1.000 | 1.000 | 1 | 1.000 | 1.000 | 1.000 |
| Tspe\_v0.7 | 1.000 | 0.589 | 1.000 | 1.000 | 1.000 | 1.000 | 1.000 | 1.000 | 1.000 | 1.000 | 1 | 1.000 | 1.000 | 1.000 |
| Tspe\_v0.8 | 1.000 | 1.000 | 1.000 | 1.000 | 1.000 | 1.000 | 1.000 | 1.000 | 1.000 | 1.000 | 1 | 1.000 | 1.000 | 1.000 |
| Tspe\_v0.9 | 0.608 | 0.605 | 0.605 | 0.639 | 0.639 | 0.605 | 0.606 | 0.607 | 0.607 | 0.608 | 1 | 1.000 | 1.000 | 1.000 |
| Tspe\_v1 | 0.642 | 0.640 | 0.640 | 1.000 | 1.000 | 0.640 | 0.641 | 0.641 | 0.641 | 0.642 | 1 | 1.000 | 1.000 | 1.000 |
| Tspe\_v1.chr | 0.532 | 0.580 | 0.594 | 1.000 | 1.000 | 0.593 | 0.532 | 0.529 | 0.587 | 0.584 | 1 | 1.000 | 1.000 | 1.000 |

Low probabilities indicate that BUSCOMP might be rating paralogues or pseudogenes and not functional orthologues of the BUSCO gene. Note that there is *no* correction for multiple testing, nor any adjustment for lack of independence between samples.

#### 5.2 Unique BUSCO and BUSCOMP Complete Genes

BUSCO and BUSCOMP `Complete` ratings were compared for each BUSCO gene to identify those genes unique to either a single assembly or a group of assemblies. The `BUSCOMP` group is excluded from this analysis, as (typically) are other redundant groups wholly contained within another group. (Inclusion of such groups is guaranteed to result in 2+ groups containing any `Complete` BUSCOs they have.)

```
Tspe_10xpseudohap2 unique Complete genes: 2 BUSCO; 0 BUSCOMP
Tspe_v0.1_canu unique Complete genes: 4 BUSCO; 0 BUSCOMP
Tspe_v0.1_flye unique Complete genes: 0 BUSCO; 0 BUSCOMP
Tspe_v0.1_necat unique Complete genes: 1 BUSCO; 0 BUSCOMP
Tspe_v0.2 unique Complete genes: 0 BUSCO; 0 BUSCOMP
Tspe_v0.3 unique Complete genes: 1 BUSCO; 0 BUSCOMP
Tspe_v0.4 unique Complete genes: 0 BUSCO; 0 BUSCOMP
Tspe_v0.5 unique Complete genes: 0 BUSCO; 0 BUSCOMP
Tspe_v0.6 unique Complete genes: 1 BUSCO; 0 BUSCOMP
Tspe_v0.7 unique Complete genes: 0 BUSCO; 0 BUSCOMP
Tspe_v0.8 unique Complete genes: 1 BUSCO; 0 BUSCOMP
Tspe_v0.9 unique Complete genes: 0 BUSCO; 0 BUSCOMP
Tspe_v1 unique Complete genes: 0 BUSCO; 0 BUSCOMP
Tspe_v1.chr unique Complete genes: 0 BUSCO; 0 BUSCOMP
Tspe_v1-full unique Complete genes: 0 BUSCO; 0 BUSCOMP
```

#### 5.3 Ratings for Missing BUSCO genes

In addition to the unique ratings (above), it can be useful to know how genes `Missing` from one assembly/group are rated in the others. These plots are generated for each assembly/group in turn. The full BUSCO (`*.busco.tdt`) and BUSCOMP (`*.LnnIDxx.buscomp.tdt`) tables are reduced to the subset of genes that are missing in the assembly/group of interest, and then the summary ratings recalculated for that subset.

In each case, three plots are made (assuming both BUSCO and BUSCOMP data is available):

1. BUSCO ratings for missing BUSCO genes.
2. BUSCOMP ratings for missing BUSCO genes. As well as being more relaxed than pure BUSCO results, this will indicate when BUSCOMP has found a gene in the focal assembly/group where BUSCO did not.
3. BUSCOMP ratings for missing BUSCOMP genes. It is expected that assemblies will be much more similar in terms of BUSCOMP coverage.

#### 5.4 Missing Tspe\_10xpseudohap2 BUSCO genes

BUSCO ratings for `Missing` Tspe\_10xpseudohap2 BUSCO genes:

BUSCOMP ratings for `Missing` Tspe\_10xpseudohap2 BUSCO genes:

BUSCOMP ratings for `Missing` Tspe\_10xpseudohap2 BUSCOMP genes:

#### 5.5 Missing Tspe\_v0.1\_canu BUSCO genes

BUSCO ratings for `Missing` Tspe\_v0.1\_canu BUSCO genes:

BUSCOMP ratings for `Missing` Tspe\_v0.1\_canu BUSCO genes:

BUSCOMP ratings for `Missing` Tspe\_v0.1\_canu BUSCOMP genes:

#### 5.6 Missing Tspe\_v0.1\_flye BUSCO genes

BUSCO ratings for `Missing` Tspe\_v0.1\_flye BUSCO genes:

BUSCOMP ratings for `Missing` Tspe\_v0.1\_flye BUSCO genes:

BUSCOMP ratings for `Missing` Tspe\_v0.1\_flye BUSCOMP genes:

#### 5.7 Missing Tspe\_v0.1\_necat BUSCO genes

BUSCO ratings for `Missing` Tspe\_v0.1\_necat BUSCO genes:

BUSCOMP ratings for `Missing` Tspe\_v0.1\_necat BUSCO genes:

BUSCOMP ratings for `Missing` Tspe\_v0.1\_necat BUSCOMP genes:

#### 5.8 Missing Tspe\_v0.2 BUSCO genes

BUSCO ratings for `Missing` Tspe\_v0.2 BUSCO genes:

BUSCOMP ratings for `Missing` Tspe\_v0.2 BUSCO genes:

BUSCOMP ratings for `Missing` Tspe\_v0.2 BUSCOMP genes:

#### 5.9 Missing Tspe\_v0.3 BUSCO genes

BUSCO ratings for `Missing` Tspe\_v0.3 BUSCO genes:

BUSCOMP ratings for `Missing` Tspe\_v0.3 BUSCO genes:

BUSCOMP ratings for `Missing` Tspe\_v0.3 BUSCOMP genes:

#### 5.10 Missing Tspe\_v0.4 BUSCO genes

BUSCO ratings for `Missing` Tspe\_v0.4 BUSCO genes:

BUSCOMP ratings for `Missing` Tspe\_v0.4 BUSCO genes:

BUSCOMP ratings for `Missing` Tspe\_v0.4 BUSCOMP genes:

#### 5.11 Missing Tspe\_v0.5 BUSCO genes

BUSCO ratings for `Missing` Tspe\_v0.5 BUSCO genes:

BUSCOMP ratings for `Missing` Tspe\_v0.5 BUSCO genes:

BUSCOMP ratings for `Missing` Tspe\_v0.5 BUSCOMP genes:

#### 5.12 Missing Tspe\_v0.6 BUSCO genes

BUSCO ratings for `Missing` Tspe\_v0.6 BUSCO genes:

BUSCOMP ratings for `Missing` Tspe\_v0.6 BUSCO genes:

BUSCOMP ratings for `Missing` Tspe\_v0.6 BUSCOMP genes:

#### 5.13 Missing Tspe\_v0.7 BUSCO genes

BUSCO ratings for `Missing` Tspe\_v0.7 BUSCO genes:

BUSCOMP ratings for `Missing` Tspe\_v0.7 BUSCO genes:

BUSCOMP ratings for `Missing` Tspe\_v0.7 BUSCOMP genes:

#### 5.14 Missing Tspe\_v0.8 BUSCO genes

BUSCO ratings for `Missing` Tspe\_v0.8 BUSCO genes:

BUSCOMP ratings for `Missing` Tspe\_v0.8 BUSCO genes:

BUSCOMP ratings for `Missing` Tspe\_v0.8 BUSCOMP genes:

#### 5.15 Missing Tspe\_v0.9 BUSCO genes

BUSCO ratings for `Missing` Tspe\_v0.9 BUSCO genes:

BUSCOMP ratings for `Missing` Tspe\_v0.9 BUSCO genes:

BUSCOMP ratings for `Missing` Tspe\_v0.9 BUSCOMP genes:

#### 5.16 Missing Tspe\_v1 BUSCO genes

BUSCO ratings for `Missing` Tspe\_v1 BUSCO genes:

BUSCOMP ratings for `Missing` Tspe\_v1 BUSCO genes:

BUSCOMP ratings for `Missing` Tspe\_v1 BUSCOMP genes:

#### 5.17 Missing Tspe\_v1.chr BUSCO genes

BUSCO ratings for `Missing` Tspe\_v1.chr BUSCO genes:

BUSCOMP ratings for `Missing` Tspe\_v1.chr BUSCO genes:

BUSCOMP ratings for `Missing` Tspe\_v1.chr BUSCOMP genes:

#### 5.18 Missing Tspe\_v1-genome BUSCO genes

BUSCO ratings for `Missing` Tspe\_v1-genome BUSCO genes:

#### 5.19 Missing Tspe\_v1-full BUSCO genes

BUSCO ratings for `Missing` Tspe\_v1-full BUSCO genes:

#### 5.20 Missing BUSCOv3 BUSCO genes

BUSCO ratings for `Missing` BUSCOv3 BUSCO genes:

BUSCOMP ratings for `Missing` BUSCOv3 BUSCO genes:

BUSCOMP ratings for `Missing` BUSCOv3 BUSCOMP genes:

#### 5.21 Missing BUSCOMP BUSCO genes

BUSCO ratings for `Missing` BUSCOMP BUSCO genes:

BUSCOMP ratings for `Missing` BUSCOMP BUSCO genes:

BUSCOMP ratings for `Missing` BUSCOMP BUSCOMP genes:

### 6 Appendix: BUSCOMP run details

```
BUSCOMP V0.12.0: run Sun Oct  3 23:04:21 2021
```

This analysis was run in:

```
/srv/scratch/ausplant/Telopea-Aug19/paper/2021-10-03.BUSCOMP/waratah-full
```

- Log file: `/srv/scratch/ausplant/Telopea-Aug19/paper/2021-10-03.BUSCOMP/waratah-full/buscov3-full.log`
- Commandline arguments: `runs=/srv/scratch/z5239484/Telopea-Aug19/data/2021-04-05.TspeWorkflow/run*` `fastadir=/srv/scratch/z5239484/Telopea-Aug19/data/2021-04-05.TspeWorkflow/,../../../annotation/2021-10-02.SAAGA/` `basefile=buscov3-full` `forks=24` `groups=waratah-full.groups.tdt`
- Full Command List: `mafft=mafft` `clustalo=clustalo` `clustalw=clustalw2` `blast+path=` `iupath=/home/z3452659/bioware/iupred/iupred` `iuchdir=T` `minregion=5` `iucut=0.2` `rpath=R` `modpurge=T` `runs=/srv/scratch/z5239484/Telopea-Aug19/data/2021-04-05.TspeWorkflow/run*` `fastadir=/srv/scratch/z5239484/Telopea-Aug19/data/2021-04-05.TspeWorkflow/,../../../annotation/2021-10-02.SAAGA/` `basefile=buscov3-full` `forks=24` `groups=waratah-full.groups.tdt` `genomesize=981953849`

#### 6.1 BUSCOMP errors

See run log for further details:

```
#ERR    00:01:03    Problem during BUSCOMP genomeGroups.: <type 'IndexError'> (genomeGroups line 2283) list index out of range
```

#### 6.2 BUSCOMP warnings

See run log for further details:

```
#WARN   00:00:00    Alias/Prefix/Group "Tspe_v0.6-aug" not found: dropping group entry
#WARN   00:00:00    Alias/Prefix/Group "Tspe_v1.gemoma.sprot.renamed.prot" not found: dropping group entry
#WARN   00:00:00    Alias/Prefix/Group "Tspe_v0.3-aug" not found: dropping group entry
#WARN   00:00:00    Alias/Prefix/Group "Tspe_v0.9-met" not found: dropping group entry
#WARN   00:00:00    Alias/Prefix/Group "Tspe_v0.1_flye-aug" not found: dropping group entry
#WARN   00:00:00    Alias/Prefix/Group "Tspe_v0.1_canu-aug" not found: dropping group entry
#WARN   00:00:00    Alias/Prefix/Group "Tspe_v0.2-aug" not found: dropping group entry
#WARN   00:00:00    Alias/Prefix/Group "Tspe_10xpseudohap2-met" not found: dropping group entry
#WARN   00:00:00    Alias/Prefix/Group "Tspe_v1.gemoma.prot" not found: dropping group entry
#WARN   00:00:00    Alias/Prefix/Group "Tspe_v1-aug" not found: dropping group entry
#WARN   00:00:00    Alias/Prefix/Group "Tspe_v0.5-aug" not found: dropping group entry
#WARN   00:00:00    Alias/Prefix/Group "Tspe_v1.gemoma" not found: dropping group entry
#WARN   00:00:00    Alias/Prefix/Group "Tspe_v1.gemoma.sprot.longest.prot" not found: dropping group entry
#WARN   00:00:00    Alias/Prefix/Group "Tspe_v0.7-met" not found: dropping group entry
#WARN   00:00:00    Alias/Prefix/Group "Tspe_v1.chr-met" not found: dropping group entry
#WARN   00:00:00    Alias/Prefix/Group "Tspe_v1.gemoma.sprot.longest" not found: dropping group entry
#WARN   00:00:00    Alias/Prefix/Group "Tspe_v0.1_necat-aug" not found: dropping group entry
#WARN   00:00:00    Alias/Prefix/Group "Tspe_v1.gemoma.sprot.renamed" not found: dropping group entry
#WARN   00:00:00    Alias/Prefix/Group "Tspe_10xpseudohap2-aug" not found: dropping group entry
#WARN   00:00:00    Alias/Prefix/Group "Tspe_v0.5-met" not found: dropping group entry
#WARN   00:00:00    Alias/Prefix/Group "Tspe_v1.chr-aug" not found: dropping group entry
#WARN   00:00:00    Alias/Prefix/Group "Tspe_v0.3-met" not found: dropping group entry
#WARN   00:00:00    Alias/Prefix/Group "Tspe_v0.9-aug" not found: dropping group entry
#WARN   00:00:00    Alias/Prefix/Group "Tspe_v0.1_flye-met" not found: dropping group entry
#WARN   00:00:00    Alias/Prefix/Group "Tspe_v1-met" not found: dropping group entry
#WARN   00:00:00    Alias/Prefix/Group "Tspe_v0.8-aug" not found: dropping group entry
#WARN   00:00:00    Alias/Prefix/Group "Tspe_v0.1_necat-met" not found: dropping group entry
#WARN   00:00:00    Alias/Prefix/Group "Tspe_v0.4-met" not found: dropping group entry
#WARN   00:00:00    Alias/Prefix/Group "Tspe_v0.8-met" not found: dropping group entry
#WARN   00:00:00    Alias/Prefix/Group "Tspe_v0.1_canu-met" not found: dropping group entry
#WARN   00:00:00    Alias/Prefix/Group "Tspe_v0.6-met" not found: dropping group entry
#WARN   00:00:00    Alias/Prefix/Group "Tspe_v0.7-aug" not found: dropping group entry
#WARN   00:00:00    Alias/Prefix/Group "Tspe_v0.4-aug" not found: dropping group entry
#WARN   00:00:00    Alias/Prefix/Group "Tspe_v0.2-met" not found: dropping group entry
#WARN   00:00:00    Alias/Prefix/Group "Tspe_10xpseudohap2-met" not found: dropping group entry
#WARN   00:00:00    Alias/Prefix/Group "Tspe_v0.1_canu-met" not found: dropping group entry
#WARN   00:00:00    Alias/Prefix/Group "Tspe_v0.1_flye-met" not found: dropping group entry
#WARN   00:00:00    Alias/Prefix/Group "Tspe_v0.1_necat-met" not found: dropping group entry
#WARN   00:00:00    Alias/Prefix/Group "Tspe_v0.2-met" not found: dropping group entry
#WARN   00:00:00    Alias/Prefix/Group "Tspe_v0.3-met" not found: dropping group entry
#WARN   00:00:00    Alias/Prefix/Group "Tspe_v0.4-met" not found: dropping group entry
#WARN   00:00:00    Alias/Prefix/Group "Tspe_v0.5-met" not found: dropping group entry
#WARN   00:00:00    Alias/Prefix/Group "Tspe_v0.6-met" not found: dropping group entry
#WARN   00:00:00    Alias/Prefix/Group "Tspe_v0.7-met" not found: dropping group entry
#WARN   00:00:00    Alias/Prefix/Group "Tspe_v0.8-met" not found: dropping group entry
#WARN   00:00:00    Alias/Prefix/Group "Tspe_v0.9-met" not found: dropping group entry
#WARN   00:00:00    Alias/Prefix/Group "Tspe_v1-met" not found: dropping group entry
#WARN   00:00:00    Alias/Prefix/Group "Tspe_v1.chr-met" not found: dropping group entry
#WARN   00:00:00    Alias/Prefix/Group "Tspe_10xpseudohap2-aug" not found: dropping group entry
#WARN   00:00:00    Alias/Prefix/Group "Tspe_v0.1_canu-aug" not found: dropping group entry
#WARN   00:00:00    Alias/Prefix/Group "Tspe_v0.1_flye-aug" not found: dropping group entry
#WARN   00:00:00    Alias/Prefix/Group "Tspe_v0.1_necat-aug" not found: dropping group entry
#WARN   00:00:00    Alias/Prefix/Group "Tspe_v0.2-aug" not found: dropping group entry
#WARN   00:00:00    Alias/Prefix/Group "Tspe_v0.3-aug" not found: dropping group entry
#WARN   00:00:00    Alias/Prefix/Group "Tspe_v0.4-aug" not found: dropping group entry
#WARN   00:00:00    Alias/Prefix/Group "Tspe_v0.5-aug" not found: dropping group entry
#WARN   00:00:00    Alias/Prefix/Group "Tspe_v0.7-aug" not found: dropping group entry
#WARN   00:00:00    Alias/Prefix/Group "Tspe_v0.6-aug" not found: dropping group entry
#WARN   00:00:00    Alias/Prefix/Group "Tspe_v0.8-aug" not found: dropping group entry
#WARN   00:00:00    Alias/Prefix/Group "Tspe_v0.9-aug" not found: dropping group entry
#WARN   00:00:00    Alias/Prefix/Group "Tspe_v1-aug" not found: dropping group entry
#WARN   00:00:00    Alias/Prefix/Group "Tspe_v1.chr-met" not found: dropping group entry
#WARN   00:00:00    Alias/Prefix/Group "Tspe_v1-aug" not found: dropping group entry
#WARN   00:00:00    Alias/Prefix/Group "Tspe_v1-met" not found: dropping group entry
#WARN   00:00:00    Alias/Prefix/Group "Tspe_v1.chr-aug" not found: dropping group entry
#WARN   00:00:00    Alias/Prefix/Group "Tspe_v1.chr-met" not found: dropping group entry
#WARN   00:00:00    Alias/Prefix/Group "Tspe_v1.gemoma" not found: dropping group entry
#WARN   00:00:00    Alias/Prefix/Group "Tspe_v1.gemoma.prot" not found: dropping group entry
#WARN   00:00:00    Alias/Prefix/Group "Tspe_v1.gemoma.sprot.longest" not found: dropping group entry
#WARN   00:00:00    Alias/Prefix/Group "Tspe_v1.gemoma.sprot.longest.prot" not found: dropping group entry
#WARN   00:00:00    Alias/Prefix/Group "Tspe_v1.gemoma.sprot.renamed" not found: dropping group entry
#WARN   00:00:00    Alias/Prefix/Group "Tspe_v1.gemoma.sprot.renamed.prot" not found: dropping group entry
#WARN   00:00:00    Alias/Prefix/Group "Tspe_v1-aug" not found: dropping group entry
#WARN   00:00:00    Alias/Prefix/Group "Tspe_v1-met" not found: dropping group entry
#WARN   00:00:00    Alias/Prefix/Group "Tspe_v1.chr-aug" not found: dropping group entry
#WARN   00:00:00    Alias/Prefix/Group "Tspe_v1.chr-met" not found: dropping group entry
#WARN   00:00:00    Alias/Prefix/Group "Tspe_v1.gemoma" not found: dropping group entry
#WARN   00:00:00    Alias/Prefix/Group "Tspe_v1.gemoma.sprot.longest" not found: dropping group entry
#WARN   00:00:00    Alias/Prefix/Group "Tspe_v1.gemoma.sprot.renamed" not found: dropping group entry
#WARN   00:00:00    Alias/Prefix/Group "Tspe_v1.gemoma.prot" not found: dropping group entry
#WARN   00:00:00    Alias/Prefix/Group "Tspe_v1.gemoma.sprot.longest.prot" not found: dropping group entry
#WARN   00:00:00    Alias/Prefix/Group "Tspe_v1.gemoma.sprot.renamed.prot" not found: dropping group entry
#WARN   01:21:37    36 Length/AlnLen mismatches from PAF file. (May be disparity with N treatment.)
```

---

**Report contents:**

- Run summary
- BUSCOMP summary
- Genome summary
- BUSCO Ratings
- BUSCO full results compilation
- BUSCOMP Sequence details and rating
- BUSCOMP re-rating of genomes
- BUSCOMP re-rating full results
- Unique BUSCO and BUSCOMP Complete genes
- Ratings for Missing BUSCO genes
- Appendix: BUSCOMP run details

---

Output generated by BUSCOMP v0.12.0 © 2019 Richard Edwards |
