## Supplementary material for "Chromosome-level *de novo* genome assembly of *Telopea speciosissima* (New South Wales waratah) using long-reads, linked-reads and Hi-C": BUSCOMP full report: BUSCOMP_full_report_buscov5.html


### BUSCOMP Full Report

###### *buscov5-full BUSCOMP Analysis*

#### *2021-10-04*

### 1 BUSCOMP Run Summary

```
BUSCOMP V0.12.1: run Mon Oct  4 21:50:25 2021
```

See the run details appendix end of this document for details of the log file, commandline parameters and runtime BUSCOMP errors and warnings.

**NOTE:** To edit this document, open `buscov5-full.N3L20ID0U.full.Rmd` in RStudio, edit and re-knit to HTML.

#### 1.1 BUSCOMP Results Summary

Assemblies can be assessed on a number of criteria, but the main ones (in the absence of a reference “truth” genome) are either to judge contiguity or completeness. NG50 and LG50 values are based on a genome size of 982.0 Mb. If the `genomesize=X` parameter was not set (see command list in appendix), this will be based on the longest assembly (see sequence stats, below).

Of the 34 assemblies analysed (34 BUSCO; 34 fasta; 34 both), 31 genomes were rated as the “best” by at least one criterion:

- `Tspe_v1.gemoma.sprot.renamed.prot`: Missing.
- `Tspe_v0.3-aug`: Complete, Missing.
- `Tspe_v0.9-met`: NG50Length, LG50Count, MaxLength, Missing.
- `Tspe_v0.1_flye-aug`: Missing.
- `Tspe_v0.1_canu-aug`: Missing.
- `Tspe_v0.2-aug`: Complete, Missing.
- `Tspe_v0.8-aug`: LG50Count, Missing.
- `Tspe_10xpseudohap2-met`: Missing.
- `Tspe_v1.gemoma.prot`: Missing.
- `Tspe_v0.1_canu-met`: Missing.
- `Tspe_v1-aug`: LG50Count, Missing.
- `Tspe_v0.5-aug`: Complete, Missing.
- `Tspe_v1.gemoma.sprot.longest.prot`: Missing.
- `Tspe_v0.7-met`: Complete, Missing.
- `Tspe_v1.chr-met`: LG50Count.
- `Tspe_v0.1_necat-aug`: Complete, Missing.
- `Tspe_10xpseudohap2-aug`: Missing.
- `Tspe_v0.5-met`: Complete, Missing, BUSCO, NoBUSCO.
- `Tspe_v1.chr-aug`: LG50Count.
- `Tspe_v0.3-met`: Complete, Missing.
- `Tspe_v0.9-aug`: NG50Length, LG50Count, MaxLength, Missing.
- `Tspe_v0.1_flye-met`: Missing.
- `Tspe_v0.8-met`: LG50Count, Missing.
- `Tspe_v1-met`: LG50Count, Missing.
- `Tspe_v0.1_necat-met`: Complete, Missing.
- `Tspe_v0.6-aug`: Complete, Missing.
- `Tspe_v0.4-met`: Complete, Missing, NoBUSCO.
- `Tspe_v0.6-met`: Complete, Missing.
- `Tspe_v0.7-aug`: Complete, Missing.
- `Tspe_v0.4-aug`: Complete, Missing.
- `Tspe_v0.2-met`: Complete, Missing.

Best assemblies by assembly contiguity critera:

- **NG50Length.** Longest NG50 contig/scaffold length (67,756,876 bp): `Tspe_v0.9-aug`, `Tspe_v0.9-met`
- **LG50Count.** Smallest LG50 contig/scaffold count (7): `Tspe_v0.8-aug`, `Tspe_v0.8-met`, `Tspe_v0.9-aug`, `Tspe_v0.9-met`, `Tspe_v1-aug`, `Tspe_v1-met`, `Tspe_v1.chr-aug`, `Tspe_v1.chr-met`
- **MaxLength.** Maximum contig/scaffold length (87,816,475 bp): `Tspe_v0.9-aug`, `Tspe_v0.9-met`

Best assemblies by completeness critera:

- **Complete.** Most Complete (Single & Duplicated) BUSCOMP sequences (99.9 %): `Tspe_v0.1_necat-aug`, `Tspe_v0.1_necat-met`, `Tspe_v0.2-aug`, `Tspe_v0.2-met`, `Tspe_v0.3-aug`, `Tspe_v0.3-met`, `Tspe_v0.4-aug`, `Tspe_v0.4-met`, `Tspe_v0.5-aug`, `Tspe_v0.5-met`, `Tspe_v0.6-aug`, `Tspe_v0.6-met`, `Tspe_v0.7-aug`, `Tspe_v0.7-met`
- **Missing.** Fewest Missing BUSCOMP sequences (0.0 %): `Tspe_10xpseudohap2-aug`, `Tspe_10xpseudohap2-met`, `Tspe_v0.1_canu-aug`, `Tspe_v0.1_canu-met`, `Tspe_v0.1_flye-aug`, `Tspe_v0.1_flye-met`, `Tspe_v0.1_necat-aug`, `Tspe_v0.1_necat-met`, `Tspe_v0.2-aug`, `Tspe_v0.2-met`, `Tspe_v0.3-aug`, `Tspe_v0.3-met`, `Tspe_v0.4-aug`, `Tspe_v0.4-met`, `Tspe_v0.5-aug`, `Tspe_v0.5-met`, `Tspe_v0.6-aug`, `Tspe_v0.6-met`, `Tspe_v0.7-aug`, `Tspe_v0.7-met`, `Tspe_v0.8-aug`, `Tspe_v0.8-met`, `Tspe_v0.9-aug`, `Tspe_v0.9-met`, `Tspe_v1-aug`, `Tspe_v1-met`, `Tspe_v1.gemoma.prot`, `Tspe_v1.gemoma.sprot.longest.prot`, `Tspe_v1.gemoma.sprot.renamed.prot`
- **BUSCO.** Most Complete (Single & Duplicated) BUSCO sequences (99.3 %): `Tspe_v0.5-met`
- **NoBUSCO.** Fewest Missing BUSCO sequences (0.2 %): `Tspe_v0.4-met`, `Tspe_v0.5-met`

### 2 Genome Summary

The following genomes and BUSCO results were analysed by BUSCOMP:

- **Tspe\_10xpseudohap2-aug**. [`BUSCO`|`Fasta`] ../fasta/Tspe\_10xpseudohap2-aug.fasta
- **Tspe\_10xpseudohap2-met**. [`BUSCO`|`Fasta`] ../fasta/Tspe\_10xpseudohap2-met.fasta
- **Tspe\_v0.1\_canu-aug**. [`BUSCO`|`Fasta`] ../fasta/Tspe\_v0.1\_canu-aug.fasta
- **Tspe\_v0.1\_canu-met**. [`BUSCO`|`Fasta`] ../fasta/Tspe\_v0.1\_canu-met.fasta
- **Tspe\_v0.1\_flye-aug**. [`BUSCO`|`Fasta`] ../fasta/Tspe\_v0.1\_flye-aug.fasta
- **Tspe\_v0.1\_flye-met**. [`BUSCO`|`Fasta`] ../fasta/Tspe\_v0.1\_flye-met.fasta
- **Tspe\_v0.1\_necat-aug**. [`BUSCO`|`Fasta`] ../fasta/Tspe\_v0.1\_necat-aug.fasta
- **Tspe\_v0.1\_necat-met**. [`BUSCO`|`Fasta`] ../fasta/Tspe\_v0.1\_necat-met.fasta
- **Tspe\_v0.2-aug**. [`BUSCO`|`Fasta`] ../fasta/Tspe\_v0.2-aug.fasta
- **Tspe\_v0.2-met**. [`BUSCO`|`Fasta`] ../fasta/Tspe\_v0.2-met.fasta
- **Tspe\_v0.3-aug**. [`BUSCO`|`Fasta`] ../fasta/Tspe\_v0.3-aug.fasta
- **Tspe\_v0.3-met**. [`BUSCO`|`Fasta`] ../fasta/Tspe\_v0.3-met.fasta
- **Tspe\_v0.4-aug**. [`BUSCO`|`Fasta`] ../fasta/Tspe\_v0.4-aug.fasta
- **Tspe\_v0.4-met**. [`BUSCO`|`Fasta`] ../fasta/Tspe\_v0.4-met.fasta
- **Tspe\_v0.5-aug**. [`BUSCO`|`Fasta`] ../fasta/Tspe\_v0.5-aug.fasta
- **Tspe\_v0.5-met**. [`BUSCO`|`Fasta`] ../fasta/Tspe\_v0.5-met.fasta
- **Tspe\_v0.6-aug**. [`BUSCO`|`Fasta`] ../fasta/Tspe\_v0.6-aug.fasta
- **Tspe\_v0.6-met**. [`BUSCO`|`Fasta`] ../fasta/Tspe\_v0.6-met.fasta
- **Tspe\_v0.7-aug**. [`BUSCO`|`Fasta`] ../fasta/Tspe\_v0.7-aug.fasta
- **Tspe\_v0.7-met**. [`BUSCO`|`Fasta`] ../fasta/Tspe\_v0.7-met.fasta
- **Tspe\_v0.8-aug**. [`BUSCO`|`Fasta`] ../fasta/Tspe\_v0.8-aug.fasta
- **Tspe\_v0.8-met**. [`BUSCO`|`Fasta`] ../fasta/Tspe\_v0.8-met.fasta
- **Tspe\_v0.9-aug**. [`BUSCO`|`Fasta`] ../fasta/Tspe\_v0.9-aug.fasta
- **Tspe\_v0.9-met**. [`BUSCO`|`Fasta`] ../fasta/Tspe\_v0.9-met.fasta
- **Tspe\_v1-aug**. [`BUSCO`|`Fasta`] ../fasta/Tspe\_v1-aug.fasta
- **Tspe\_v1-met**. [`BUSCO`|`Fasta`] ../fasta/Tspe\_v1-met.fasta
- **Tspe\_v1.chr-aug**. [`BUSCO`|`Fasta`] ../fasta/Tspe\_v1.chr-aug.fasta
- **Tspe\_v1.chr-met**. [`BUSCO`|`Fasta`] ../fasta/Tspe\_v1.chr-met.fasta
- **Tspe\_v1.gemoma**. [`BUSCO`|`Fasta`] ../../../annotation/2021-10-02.SAAGA/Tspe\_v1.gemoma.fna
- **Tspe\_v1.gemoma.prot**. [`BUSCO`] Tspe\_v1.gemoma.prot (no Fasta)
- **Tspe\_v1.gemoma.sprot.longest**. [`BUSCO`|`Fasta`] ../../../annotation/2021-10-02.SAAGA/Tspe\_v1.gemoma.sprot.longest.fna
- **Tspe\_v1.gemoma.sprot.longest.prot**. [`BUSCO`] Tspe\_v1.gemoma.sprot.longest.prot (no Fasta)
- **Tspe\_v1.gemoma.sprot.renamed**. [`BUSCO`|`Fasta`] ../../../annotation/2021-10-02.SAAGA/Tspe\_v1.gemoma.sprot.renamed.fna
- **Tspe\_v1.gemoma.sprot.renamed.prot**. [`BUSCO`] Tspe\_v1.gemoma.sprot.renamed.prot (no Fasta)

| Genome | Description | SeqNum | TotLength | MinLength | MaxLength | MeanLength | MedLength | N50Length | L50Count | CtgNum | N50Ctg | L50Ctg | NG50Length | LG50Count | GapLength | GapCount | GC |
| --- | --- | --- | --- | --- | --- | --- | --- | --- | --- | --- | --- | --- | --- | --- | --- | --- | --- |
| Tspe\_10xpseudohap2-aug | ../fasta/Tspe\_10xpseudohap2-aug.fasta | 27610 | 850975689 | 1000 | 7917366 | 30821.285 | 2990 | 874466 | 247 | 43951 | 72725 | 3268 | 694813 | 331 | 18076790 | 16341 | 40.10 |
| Tspe\_10xpseudohap2-met | ../fasta/Tspe\_10xpseudohap2-met.fasta | 27610 | 850975689 | 1000 | 7917366 | 30821.285 | 2990 | 874466 | 247 | 43951 | 72725 | 3268 | 694813 | 331 | 18076790 | 16341 | 40.10 |
| Tspe\_v0.1\_canu-aug | ../fasta/Tspe\_v0.1\_canu-aug.fasta | 3983 | 981953849 | 1138 | 12971499 | 246536.241 | 58209 | 1848137 | 132 | 3983 | 1848137 | 132 | 1848137 | 132 | 0 | 0 | 39.95 |
| Tspe\_v0.1\_canu-met | ../fasta/Tspe\_v0.1\_canu-met.fasta | 3983 | 981953849 | 1138 | 12971499 | 246536.241 | 58209 | 1848137 | 132 | 3983 | 1848137 | 132 | 1848137 | 132 | 0 | 0 | 39.95 |
| Tspe\_v0.1\_flye-aug | ../fasta/Tspe\_v0.1\_flye-aug.fasta | 2445 | 857703641 | 505 | 12987791 | 350799.035 | 31428 | 2271126 | 94 | 2484 | 2199532 | 101 | 1823111 | 125 | 3900 | 39 | 40.47 |
| Tspe\_v0.1\_flye-met | ../fasta/Tspe\_v0.1\_flye-met.fasta | 2445 | 857703641 | 505 | 12987791 | 350799.035 | 31428 | 2271126 | 94 | 2484 | 2199532 | 101 | 1823111 | 125 | 3900 | 39 | 40.47 |
| Tspe\_v0.1\_necat-aug | ../fasta/Tspe\_v0.1\_necat-aug.fasta | 365 | 842143239 | 563 | 37347493 | 2307241.751 | 155267 | 10701597 | 24 | 365 | 10701597 | 24 | 9180758 | 31 | 0 | 0 | 40.15 |
| Tspe\_v0.1\_necat-met | ../fasta/Tspe\_v0.1\_necat-met.fasta | 365 | 842143239 | 563 | 37347493 | 2307241.751 | 155267 | 10701597 | 24 | 365 | 10701597 | 24 | 9180758 | 31 | 0 | 0 | 40.15 |
| Tspe\_v0.2-aug | ../fasta/Tspe\_v0.2-aug.fasta | 210 | 827757806 | 1773 | 37347493 | 3941703.838 | 1153652 | 10955845 | 23 | 210 | 10955845 | 23 | 9180758 | 31 | 0 | 0 | 40.13 |
| Tspe\_v0.2-met | ../fasta/Tspe\_v0.2-met.fasta | 210 | 827757806 | 1773 | 37347493 | 3941703.838 | 1153652 | 10955845 | 23 | 210 | 10955845 | 23 | 9180758 | 31 | 0 | 0 | 40.13 |
| Tspe\_v0.3-aug | ../fasta/Tspe\_v0.3-aug.fasta | 209 | 834569695 | 3516 | 37634595 | 3993156.435 | 1227984 | 11046571 | 23 | 209 | 11046571 | 23 | 9263258 | 31 | 0 | 0 | 40.27 |
| Tspe\_v0.3-met | ../fasta/Tspe\_v0.3-met.fasta | 209 | 834569695 | 3516 | 37634595 | 3993156.435 | 1227984 | 11046571 | 23 | 209 | 11046571 | 23 | 9263258 | 31 | 0 | 0 | 40.27 |
| Tspe\_v0.4-aug | ../fasta/Tspe\_v0.4-aug.fasta | 209 | 835074331 | 3532 | 37649015 | 3995570.962 | 1228174 | 11052047 | 23 | 209 | 11052047 | 23 | 9267273 | 31 | 0 | 0 | 40.25 |
| Tspe\_v0.4-met | ../fasta/Tspe\_v0.4-met.fasta | 209 | 835074331 | 3532 | 37649015 | 3995570.962 | 1228174 | 11052047 | 23 | 209 | 11052047 | 23 | 9267273 | 31 | 0 | 0 | 40.25 |
| Tspe\_v0.5-aug | ../fasta/Tspe\_v0.5-aug.fasta | 138 | 835372011 | 3532 | 49795992 | 6053420.370 | 2458095 | 16474627 | 15 | 205 | 11052047 | 23 | 14087303 | 20 | 297663 | 67 | 40.25 |
| Tspe\_v0.5-met | ../fasta/Tspe\_v0.5-met.fasta | 138 | 835372011 | 3532 | 49795992 | 6053420.370 | 2458095 | 16474627 | 15 | 205 | 11052047 | 23 | 14087303 | 20 | 297663 | 67 | 40.25 |
| Tspe\_v0.6-aug | ../fasta/Tspe\_v0.6-aug.fasta | 138 | 835438898 | 3484 | 49793446 | 6053905.058 | 2461216 | 16468668 | 15 | 159 | 14704285 | 17 | 14096844 | 20 | 256618 | 21 | 40.27 |
| Tspe\_v0.6-met | ../fasta/Tspe\_v0.6-met.fasta | 138 | 835438898 | 3484 | 49793446 | 6053905.058 | 2461216 | 16468668 | 15 | 159 | 14704285 | 17 | 14096844 | 20 | 256618 | 21 | 40.27 |
| Tspe\_v0.7-aug | ../fasta/Tspe\_v0.7-aug.fasta | 128 | 833847971 | 3484 | 49793446 | 6514437.273 | 2967336 | 16468668 | 15 | 148 | 14704285 | 17 | 14096844 | 20 | 244118 | 20 | 40.27 |
| Tspe\_v0.7-met | ../fasta/Tspe\_v0.7-met.fasta | 128 | 833847971 | 3484 | 49793446 | 6514437.273 | 2967336 | 16468668 | 15 | 148 | 14704285 | 17 | 14096844 | 20 | 244118 | 20 | 40.27 |
| Tspe\_v0.8-aug | ../fasta/Tspe\_v0.8-aug.fasta | 2357 | 833952765 | 5 | 87707951 | 353819.586 | 11506 | 68901929 | 6 | 3575 | 1773000 | 131 | 67726057 | 7 | 982622 | 1218 | 40.03 |
| Tspe\_v0.8-met | ../fasta/Tspe\_v0.8-met.fasta | 2357 | 833952765 | 5 | 87707951 | 353819.586 | 11506 | 68901929 | 6 | 3575 | 1773000 | 131 | 67726057 | 7 | 982622 | 1218 | 40.03 |
| Tspe\_v0.9-aug | ../fasta/Tspe\_v0.9-aug.fasta | 1399 | 825066408 | 974 | 87816475 | 589754.402 | 18113 | 69015533 | 6 | 1566 | 12206888 | 21 | 67756876 | 7 | 316262 | 167 | 40.12 |
| Tspe\_v0.9-met | ../fasta/Tspe\_v0.9-met.fasta | 1399 | 825066408 | 974 | 87816475 | 589754.402 | 18113 | 69015533 | 6 | 1566 | 12206888 | 21 | 67756876 | 7 | 316262 | 167 | 40.12 |
| Tspe\_v1-aug | ../fasta/Tspe\_v1-aug.fasta | 1289 | 823061212 | 974 | 87811904 | 638526.929 | 19505 | 69013595 | 6 | 1452 | 12206888 | 21 | 67754596 | 7 | 16300 | 163 | 40.11 |
| Tspe\_v1-met | ../fasta/Tspe\_v1-met.fasta | 1289 | 823061212 | 974 | 87811904 | 638526.929 | 19505 | 69013595 | 6 | 1452 | 12206888 | 21 | 67754596 | 7 | 16300 | 163 | 40.11 |
| Tspe\_v1.chr-aug | ../fasta/Tspe\_v1.chr-aug.fasta | 11 | 774398234 | 57317785 | 87811904 | 70399839.455 | 69013595 | 69013595 | 6 | 158 | 13444441 | 19 | 67754596 | 7 | 14700 | 147 | 40.06 |
| Tspe\_v1.chr-met | ../fasta/Tspe\_v1.chr-met.fasta | 11 | 774398234 | 57317785 | 87811904 | 70399839.455 | 69013595 | 69013595 | 6 | 158 | 13444441 | 19 | 67754596 | 7 | 14700 | 147 | 40.06 |
| Tspe\_v1.gemoma | ../../../annotation/2021-10-02.SAAGA/Tspe\_v1.gemoma.fna | 46877 | 53235801 | 21 | 15915 | 1135.649 | 951 | 1533 | 11287 | 46879 | 1533 | 11287 | 0 | -1 | 200 | 2 | 44.47 |
| Tspe\_v1.gemoma.sprot.longest | ../../../annotation/2021-10-02.SAAGA/Tspe\_v1.gemoma.sprot.longest.fna | 40158 | 43793349 | 21 | 15915 | 1090.526 | 885 | 1518 | 9261 | 40160 | 1518 | 9261 | 0 | -1 | 200 | 2 | 44.48 |
| Tspe\_v1.gemoma.sprot.renamed | ../../../annotation/2021-10-02.SAAGA/Tspe\_v1.gemoma.sprot.renamed.fna | 46877 | 53235801 | 21 | 15915 | 1135.649 | 951 | 1533 | 11287 | 46879 | 1533 | 11287 | 0 | -1 | 200 | 2 | 44.47 |

**NOTE:** `NG50Length` and `LG50Count` statistics use `genomesize=X` or the biggest assembly loaded (981.95 Mb). If BUSCOMP has been run more than once on the same data (*e.g.* to update descriptions or sorting), please make sure that a consistent genome size is used, or these values may be wrong. If in doubt, run with `force=T` and force regeneration of statistics.

#### 2.2 Genome coverage assessment plots

In general, a good assembly will be approx. the same size as the genome and in as few pieces as possible. Any assembly smaller than the predicted genome size is clearly missing coverage. Assemblies bigger than the genome size might still be missing chunks of the genome if redundancy/duplication is a problem. In the following plot, the grey line marks the given genome size of 982.0 Mb.

A better indicator of the overall coverage of the genome is the number of `Missing` BUSCO genes. As BUSCO is highly dependent on the accuracy of the sequence and the gene models it makes, the `Missing` BUSCOMP ratings arguably give a more consistent proxy for genome completeness. NOTE: this says nothing about the fragmentation or completeness of the genes themselves.

#### 2.3 Genome contiguity assessment plots

In general, a good assembly will be in fewer, bigger pieces. This is approximated using NG50 and LG50, which are the min. length and number of contigs/scaffolds required to cover at least half the genome. These stats use the given genome size of 982.0 Mb.

**NOTE:** To modify these plots and tables, edit the `*.genomes.tdt` and `*.NxLxxIDxx.rdata.tdt` files and re-knit the `*.NxLxxIDxx.Rmd` file.

### 3 BUSCO Ratings

Compiled BUSCO results for 34 assemblies and 7 groups have been saved in `buscov5-full.genomes.tdt`. BUSCO ratings are defined (quoting from the BUSCO v3 User Guide as:

- `Complete`: Single-copy hits where “BUSCO matches have scored within the expected range of scores and within the expected range of length alignments to the BUSCO profile.”
- `Duplicated`: As `Complete` but 2+ copies.
- `Fragmented`: “BUSCO matches … within the range of scores but not within the range of length alignments to the BUSCO profile.”
- `Missing`: “Either no significant matches at all, or the BUSCO matches scored below the range of scores for the BUSCO profile.”

| Genome | N | Complete | Single | Duplicated | Fragmented | Missing |
| --- | --- | --- | --- | --- | --- | --- |
| Tspe\_10xpseudohap2-aug | 1614 | 1522 | 1377 | 145 | 39 | 53 |
| Tspe\_10xpseudohap2-met | 1614 | 1567 | 1389 | 178 | 32 | 15 |
| Tspe\_v0.1\_canu-aug | 1614 | 1345 | 1201 | 144 | 84 | 185 |
| Tspe\_v0.1\_canu-met | 1614 | 1520 | 1275 | 245 | 61 | 33 |
| Tspe\_v0.1\_flye-aug | 1614 | 1389 | 1313 | 76 | 74 | 151 |
| Tspe\_v0.1\_flye-met | 1614 | 1519 | 1374 | 145 | 55 | 40 |
| Tspe\_v0.1\_necat-aug | 1614 | 1381 | 1304 | 77 | 81 | 152 |
| Tspe\_v0.1\_necat-met | 1614 | 1533 | 1386 | 147 | 57 | 24 |
| Tspe\_v0.2-aug | 1614 | 1368 | 1289 | 79 | 91 | 155 |
| Tspe\_v0.2-met | 1614 | 1532 | 1389 | 143 | 58 | 24 |
| Tspe\_v0.3-aug | 1614 | 1533 | 1404 | 129 | 30 | 51 |
| Tspe\_v0.3-met | 1614 | 1590 | 1412 | 178 | 19 | 5 |
| Tspe\_v0.4-aug | 1614 | 1555 | 1408 | 147 | 23 | 36 |
| Tspe\_v0.4-met | 1614 | 1602 | 1410 | 192 | 9 | 3 |
| Tspe\_v0.5-aug | 1614 | 1558 | 1412 | 146 | 22 | 34 |
| Tspe\_v0.5-met | 1614 | 1603 | 1411 | 192 | 8 | 3 |
| Tspe\_v0.6-aug | 1614 | 1532 | 1400 | 132 | 34 | 48 |
| Tspe\_v0.6-met | 1614 | 1587 | 1416 | 171 | 20 | 7 |
| Tspe\_v0.7-aug | 1614 | 1533 | 1403 | 130 | 32 | 49 |
| Tspe\_v0.7-met | 1614 | 1587 | 1416 | 171 | 20 | 7 |
| Tspe\_v0.8-aug | 1614 | 1457 | 1369 | 88 | 51 | 106 |
| Tspe\_v0.8-met | 1614 | 1544 | 1405 | 139 | 49 | 21 |
| Tspe\_v0.9-aug | 1614 | 1527 | 1390 | 137 | 31 | 56 |
| Tspe\_v0.9-met | 1614 | 1579 | 1399 | 180 | 27 | 8 |
| Tspe\_v1-aug | 1614 | 1526 | 1389 | 137 | 32 | 56 |
| Tspe\_v1-met | 1614 | 1579 | 1399 | 180 | 27 | 8 |
| Tspe\_v1.chr-aug | 1614 | 1512 | 1379 | 133 | 30 | 72 |
| Augustus | 1614 | 1599 | 1590 | 9 | 5 | 10 |
| Tspe\_v1.chr-met | 1614 | 1561 | 1383 | 178 | 21 | 32 |
| Tspe\_v1-genome | 1614 | 1596 | 1479 | 117 | 13 | 5 |
| MetaEuk | 1614 | 1609 | 1549 | 60 | 4 | 1 |
| Tspe\_v1.gemoma | 1614 | 1521 | 1332 | 189 | 54 | 39 |
| Tspe\_v1.gemoma.prot | 1614 | 1524 | 1334 | 190 | 52 | 38 |
| Tspe\_v1.gemoma.sprot.longest | 1614 | 1521 | 1354 | 167 | 54 | 39 |
| Tspe\_v1.gemoma.sprot.longest.prot | 1614 | 1524 | 1357 | 167 | 52 | 38 |
| Tspe\_v1.gemoma.sprot.renamed | 1614 | 1521 | 1332 | 189 | 54 | 39 |
| Tspe\_v1-transcriptome | 1614 | 1521 | 1354 | 167 | 54 | 39 |
| Tspe\_v1.gemoma.sprot.renamed.prot | 1614 | 1524 | 1334 | 190 | 52 | 38 |
| Tspe\_v1-proteome | 1614 | 1524 | 1357 | 167 | 52 | 38 |
| Tspe\_v1-full | 1614 | 1598 | 1495 | 103 | 12 | 4 |
| BUSCOMP | 1614 | 1613 | 1607 | 6 | 0 | 1 |

#### 3.1 Genome Groups

`BUSCOMP` compiled the following groups of genomes (where BUSCO data was loaded), keeping the “best” rating for each BUSCO gene across the group:

- **Augustus**: `Tspe_10xpseudohap2-aug` `Tspe_v0.1_canu-aug` `Tspe_v0.1_flye-aug` `Tspe_v0.1_necat-aug` `Tspe_v0.2-aug` `Tspe_v0.3-aug` `Tspe_v0.4-aug` `Tspe_v0.5-aug` `Tspe_v0.6-aug` `Tspe_v0.7-aug` `Tspe_v0.8-aug` `Tspe_v0.9-aug` `Tspe_v1-aug` `Tspe_v1.chr-aug`
- **Tspe\_v1-genome**: `Tspe_v1-aug` `Tspe_v1-met` `Tspe_v1.chr-aug` `Tspe_v1.chr-met`
- **MetaEuk**: `Tspe_10xpseudohap2-met` `Tspe_v0.1_canu-met` `Tspe_v0.1_flye-met` `Tspe_v0.1_necat-met` `Tspe_v0.2-met` `Tspe_v0.3-met` `Tspe_v0.4-met` `Tspe_v0.5-met` `Tspe_v0.6-met` `Tspe_v0.7-met` `Tspe_v0.8-met` `Tspe_v0.9-met` `Tspe_v1-met` `Tspe_v1.chr-met`
- **Tspe\_v1-transcriptome**: `Tspe_v1.gemoma` `Tspe_v1.gemoma.sprot.longest` `Tspe_v1.gemoma.sprot.renamed`
- **Tspe\_v1-proteome**: `Tspe_v1.gemoma.prot` `Tspe_v1.gemoma.sprot.longest.prot` `Tspe_v1.gemoma.sprot.renamed.prot`
- **Tspe\_v1-full**: `Tspe_v1-aug` `Tspe_v1-met` `Tspe_v1.chr-aug` `Tspe_v1.chr-met` `Tspe_v1.gemoma` `Tspe_v1.gemoma.prot` `Tspe_v1.gemoma.sprot.longest` `Tspe_v1.gemoma.sprot.longest.prot` `Tspe_v1.gemoma.sprot.renamed` `Tspe_v1.gemoma.sprot.renamed.prot`
- **BUSCOMP**: `Tspe_10xpseudohap2-aug` `Tspe_10xpseudohap2-met` `Tspe_v0.1_canu-aug` `Tspe_v0.1_canu-met` `Tspe_v0.1_flye-aug` `Tspe_v0.1_flye-met` `Tspe_v0.1_necat-aug` `Tspe_v0.1_necat-met` `Tspe_v0.2-aug` `Tspe_v0.2-met` `Tspe_v0.3-aug` `Tspe_v0.3-met` `Tspe_v0.4-aug` `Tspe_v0.4-met` `Tspe_v0.5-aug` `Tspe_v0.5-met` `Tspe_v0.6-aug` `Tspe_v0.6-met` `Tspe_v0.7-aug` `Tspe_v0.7-met` `Tspe_v0.8-aug` `Tspe_v0.8-met` `Tspe_v0.9-aug` `Tspe_v0.9-met` `Tspe_v1-aug` `Tspe_v1-met` `Tspe_v1.chr-aug` `Tspe_v1.chr-met` `Tspe_v1.gemoma` `Tspe_v1.gemoma.prot` `Tspe_v1.gemoma.sprot.longest` `Tspe_v1.gemoma.sprot.longest.prot` `Tspe_v1.gemoma.sprot.renamed` `Tspe_v1.gemoma.sprot.renamed.prot`

#### 3.2 BUSCO Summary

```
Tspe_10xpseudohap2-aug BUSCO Results:
        C:94.3%[S:85.3%,D:9.0%],F:2.4%,M:3.3%,n:1614

Tspe_10xpseudohap2-met BUSCO Results:
        C:97.1%[S:86.1%,D:11.0%],F:2.0%,M:0.9%,n:1614

Tspe_v0.1_canu-aug BUSCO Results:
        C:83.3%[S:74.4%,D:8.9%],F:5.2%,M:11.5%,n:1614

Tspe_v0.1_canu-met BUSCO Results:
        C:94.2%[S:79.0%,D:15.2%],F:3.8%,M:2.0%,n:1614

Tspe_v0.1_flye-aug BUSCO Results:
        C:86.1%[S:81.4%,D:4.7%],F:4.6%,M:9.4%,n:1614

Tspe_v0.1_flye-met BUSCO Results:
        C:94.1%[S:85.1%,D:9.0%],F:3.4%,M:2.5%,n:1614

Tspe_v0.1_necat-aug BUSCO Results:
        C:85.6%[S:80.8%,D:4.8%],F:5.0%,M:9.4%,n:1614

Tspe_v0.1_necat-met BUSCO Results:
        C:95.0%[S:85.9%,D:9.1%],F:3.5%,M:1.5%,n:1614

Tspe_v0.2-aug BUSCO Results:
        C:84.8%[S:79.9%,D:4.9%],F:5.6%,M:9.6%,n:1614

Tspe_v0.2-met BUSCO Results:
        C:94.9%[S:86.1%,D:8.9%],F:3.6%,M:1.5%,n:1614

Tspe_v0.3-aug BUSCO Results:
        C:95.0%[S:87.0%,D:8.0%],F:1.9%,M:3.2%,n:1614

Tspe_v0.3-met BUSCO Results:
        C:98.5%[S:87.5%,D:11.0%],F:1.2%,M:0.3%,n:1614

Tspe_v0.4-aug BUSCO Results:
        C:96.3%[S:87.2%,D:9.1%],F:1.4%,M:2.2%,n:1614

Tspe_v0.4-met BUSCO Results:
        C:99.3%[S:87.4%,D:11.9%],F:0.6%,M:0.2%,n:1614

Tspe_v0.5-aug BUSCO Results:
        C:96.5%[S:87.5%,D:9.0%],F:1.4%,M:2.1%,n:1614

Tspe_v0.5-met BUSCO Results:
        C:99.3%[S:87.4%,D:11.9%],F:0.5%,M:0.2%,n:1614

Tspe_v0.6-aug BUSCO Results:
        C:94.9%[S:86.7%,D:8.2%],F:2.1%,M:3.0%,n:1614

Tspe_v0.6-met BUSCO Results:
        C:98.3%[S:87.7%,D:10.6%],F:1.2%,M:0.4%,n:1614

Tspe_v0.7-aug BUSCO Results:
        C:95.0%[S:86.9%,D:8.1%],F:2.0%,M:3.0%,n:1614

Tspe_v0.7-met BUSCO Results:
        C:98.3%[S:87.7%,D:10.6%],F:1.2%,M:0.4%,n:1614

Tspe_v0.8-aug BUSCO Results:
        C:90.3%[S:84.8%,D:5.5%],F:3.2%,M:6.6%,n:1614

Tspe_v0.8-met BUSCO Results:
        C:95.7%[S:87.1%,D:8.6%],F:3.0%,M:1.3%,n:1614

Tspe_v0.9-aug BUSCO Results:
        C:94.6%[S:86.1%,D:8.5%],F:1.9%,M:3.5%,n:1614

Tspe_v0.9-met BUSCO Results:
        C:97.8%[S:86.7%,D:11.2%],F:1.7%,M:0.5%,n:1614

Tspe_v1-aug BUSCO Results:
        C:94.5%[S:86.1%,D:8.5%],F:2.0%,M:3.5%,n:1614

Tspe_v1-met BUSCO Results:
        C:97.8%[S:86.7%,D:11.2%],F:1.7%,M:0.5%,n:1614

Tspe_v1.chr-aug BUSCO Results:
        C:93.7%[S:85.4%,D:8.2%],F:1.9%,M:4.5%,n:1614

Augustus BUSCO Results:
        C:99.1%[S:98.5%,D:0.6%],F:0.3%,M:0.6%,n:1614

Tspe_v1.chr-met BUSCO Results:
        C:96.7%[S:85.7%,D:11.0%],F:1.3%,M:2.0%,n:1614

Tspe_v1-genome BUSCO Results:
        C:98.9%[S:91.6%,D:7.2%],F:0.8%,M:0.3%,n:1614

MetaEuk BUSCO Results:
        C:99.7%[S:96.0%,D:3.7%],F:0.2%,M:0.1%,n:1614

Tspe_v1.gemoma BUSCO Results:
        C:94.2%[S:82.5%,D:11.7%],F:3.3%,M:2.4%,n:1614

Tspe_v1.gemoma.prot BUSCO Results:
        C:94.4%[S:82.7%,D:11.8%],F:3.2%,M:2.4%,n:1614

Tspe_v1.gemoma.sprot.longest BUSCO Results:
        C:94.2%[S:83.9%,D:10.3%],F:3.3%,M:2.4%,n:1614

Tspe_v1.gemoma.sprot.longest.prot BUSCO Results:
        C:94.4%[S:84.1%,D:10.3%],F:3.2%,M:2.4%,n:1614

Tspe_v1.gemoma.sprot.renamed BUSCO Results:
        C:94.2%[S:82.5%,D:11.7%],F:3.3%,M:2.4%,n:1614

Tspe_v1-transcriptome BUSCO Results:
        C:94.2%[S:83.9%,D:10.3%],F:3.3%,M:2.4%,n:1614

Tspe_v1.gemoma.sprot.renamed.prot BUSCO Results:
        C:94.4%[S:82.7%,D:11.8%],F:3.2%,M:2.4%,n:1614

Tspe_v1-proteome BUSCO Results:
        C:94.4%[S:84.1%,D:10.3%],F:3.2%,M:2.4%,n:1614

Tspe_v1-full BUSCO Results:
        C:99.0%[S:92.6%,D:6.4%],F:0.7%,M:0.2%,n:1614

BUSCOMP BUSCO Results:
        C:99.9%[S:99.6%,D:0.4%],F:0.0%,M:0.1%,n:1614
```

#### 3.3 BUSCO Gene Details

Full BUSCO results with ratings for each gene have been compiled in `buscov5-full.busco.tdt`:

#### 3.4 Genome Group BUSCO charts

### 4 BUSCOMP Ratings

The best complete BUSCO hit results (based on `Score` and `Length`) have been compiled in `buscov5-full.buscoseq.tdt`. The `Genome` field indicates the assembly with the best hit, which is followed by details of that hit (`Contig`, `Start`, `End`, `Score`, `Length`). BUSCOMP ratings for each assembly are then given in subsequent fields:

```
* `Identical`: 100% coverage and 100% identity in at least one contig/scaffold.
* `Complete`: 95%+ Coverage in a single contig/scaffold. (Note: accuracy/identity is not considered.)
* `Duplicated`: 95%+ Coverage in 2+ contigs/scaffolds.
* `Fragmented`: 95%+ combined coverage but not in any single contig/scaffold.
* `Partial`: 40-95% combined coverage.
* `Ghost`: Hits meeting local cutoff but <40% combined coverage.
* `Missing`: No hits meeting local cutoff.
```

#### 4.1 BUSCOSeq Rating Summary

BUSCOMP ratings (see above) are compiled to summary statistics in `buscov5-full.N3L20ID0U.ratings.tdt`. Note that `Identical` ratings in this table will also be rated as `Complete`, which in turn are `Single` or `Duplicated`. Percentage summaries are plotted below, along with a BUSCO-style one-line summary per assembly/group.

**NOTE:** Group summaries do not include `Identical` ratings.

| X. | Genome | N | Identical | Complete | Single | Duplicated | Fragmented | Partial | Ghost | Missing |
| --- | --- | --- | --- | --- | --- | --- | --- | --- | --- | --- |
| 1 | Tspe\_10xpseudohap2-aug | 1569 | 735 | 1521 | 1351 | 170 | 37 | 7 | 4 | 0 |
| 2 | Tspe\_10xpseudohap2-met | 1569 | 735 | 1521 | 1351 | 170 | 37 | 7 | 4 | 0 |
| 3 | Tspe\_v0.1\_canu-aug | 1569 | 114 | 1560 | 1242 | 318 | 7 | 1 | 1 | 0 |
| 4 | Tspe\_v0.1\_canu-met | 1569 | 114 | 1560 | 1242 | 318 | 7 | 1 | 1 | 0 |
| 5 | Tspe\_v0.1\_flye-aug | 1569 | 539 | 1552 | 1381 | 171 | 13 | 3 | 1 | 0 |
| 6 | Tspe\_v0.1\_flye-met | 1569 | 539 | 1552 | 1381 | 171 | 13 | 3 | 1 | 0 |
| 7 | Tspe\_v0.1\_necat-aug | 1569 | 182 | 1567 | 1397 | 170 | 0 | 2 | 0 | 0 |
| 8 | Tspe\_v0.1\_necat-met | 1569 | 182 | 1567 | 1397 | 170 | 0 | 2 | 0 | 0 |
| 9 | Tspe\_v0.2-aug | 1569 | 182 | 1567 | 1401 | 166 | 0 | 2 | 0 | 0 |
| 10 | Tspe\_v0.2-met | 1569 | 182 | 1567 | 1401 | 166 | 0 | 2 | 0 | 0 |
| 11 | Tspe\_v0.3-aug | 1569 | 525 | 1567 | 1401 | 166 | 0 | 2 | 0 | 0 |
| 12 | Tspe\_v0.3-met | 1569 | 525 | 1567 | 1401 | 166 | 0 | 2 | 0 | 0 |
| 13 | Tspe\_v0.4-aug | 1569 | 664 | 1567 | 1400 | 167 | 0 | 2 | 0 | 0 |
| 14 | Tspe\_v0.4-met | 1569 | 664 | 1567 | 1400 | 167 | 0 | 2 | 0 | 0 |
| 15 | Tspe\_v0.5-aug | 1569 | 663 | 1567 | 1399 | 168 | 0 | 1 | 1 | 0 |
| 16 | Tspe\_v0.5-met | 1569 | 663 | 1567 | 1399 | 168 | 0 | 1 | 1 | 0 |
| 17 | Tspe\_v0.6-aug | 1569 | 544 | 1567 | 1400 | 167 | 0 | 1 | 1 | 0 |
| 18 | Tspe\_v0.6-met | 1569 | 544 | 1567 | 1400 | 167 | 0 | 1 | 1 | 0 |
| 19 | Tspe\_v0.7-aug | 1569 | 544 | 1567 | 1400 | 167 | 0 | 1 | 1 | 0 |
| 20 | Tspe\_v0.7-met | 1569 | 544 | 1567 | 1400 | 167 | 0 | 1 | 1 | 0 |
| 21 | Tspe\_v0.8-aug | 1569 | 277 | 1528 | 1382 | 146 | 31 | 3 | 7 | 0 |
| 22 | Tspe\_v0.8-met | 1569 | 277 | 1528 | 1382 | 146 | 31 | 3 | 7 | 0 |
| 23 | Tspe\_v0.9-aug | 1569 | 827 | 1529 | 1382 | 147 | 29 | 5 | 6 | 0 |
| 24 | Tspe\_v0.9-met | 1569 | 827 | 1529 | 1382 | 147 | 29 | 5 | 6 | 0 |
| 25 | Tspe\_v1-aug | 1569 | 827 | 1529 | 1382 | 147 | 29 | 5 | 6 | 0 |
| 26 | Tspe\_v1-met | 1569 | 827 | 1529 | 1382 | 147 | 29 | 5 | 6 | 0 |
| 27 | Tspe\_v1.chr-aug | 1569 | 819 | 1514 | 1371 | 143 | 1 | 21 | 20 | 13 |
| 28 | Augustus | 1569 | 0 | 1569 | 1444 | 125 | 0 | 0 | 0 | 0 |
| 29 | Tspe\_v1.chr-met | 1569 | 819 | 1514 | 1371 | 143 | 1 | 21 | 20 | 13 |
| 30 | Tspe\_v1-genome | 1569 | 0 | 1529 | 1385 | 144 | 29 | 5 | 6 | 0 |
| 31 | MetaEuk | 1569 | 0 | 1569 | 1444 | 125 | 0 | 0 | 0 | 0 |
| 32 | Tspe\_v1.gemoma | 1569 | 689 | 1313 | 1165 | 148 | 18 | 209 | 10 | 19 |
| 34 | Tspe\_v1.gemoma.sprot.longest | 1569 | 689 | 1313 | 1181 | 132 | 18 | 209 | 10 | 19 |
| 36 | Tspe\_v1.gemoma.sprot.renamed | 1569 | 689 | 1313 | 1165 | 148 | 18 | 209 | 10 | 19 |
| 37 | Tspe\_v1-transcriptome | 1569 | 0 | 1313 | 1181 | 132 | 18 | 209 | 10 | 19 |
| 39 | Tspe\_v1-proteome | 1569 | 0 | 0 | 0 | 0 | 0 | 0 | 0 | 0 |
| 40 | Tspe\_v1-full | 1569 | 0 | 1529 | 1418 | 111 | 29 | 11 | 0 | 0 |
| 41 | BUSCOMP | 1569 | 0 | 1569 | 1470 | 99 | 0 | 0 | 0 | 0 |

```
Augustus BUSCOMP Results [1599 (99.07%) Complete BUSCOs; 0 (0.00%) BUSCOMP Seqs]:
        C:100.0%[S:92.0%,D:8.0%],F:0.0%,P:0.0%,G:0.0%,M:0.0%,n:1569

BUSCOMP BUSCOMP Results [1613 (99.94%) Complete BUSCOs; 0 (0.00%) BUSCOMP Seqs]:
        C:100.0%[S:93.7%,D:6.3%],F:0.0%,P:0.0%,G:0.0%,M:0.0%,n:1569

MetaEuk BUSCOMP Results [1609 (99.69%) Complete BUSCOs; 0 (0.00%) BUSCOMP Seqs]:
        C:100.0%[S:92.0%,D:8.0%],F:0.0%,P:0.0%,G:0.0%,M:0.0%,n:1569

Tspe_10xpseudohap2-aug BUSCOMP Results [1522 (94.30%) Complete BUSCOs; 380 (24.22%) BUSCOMP Seqs]:
        C:96.9%[S:86.1%,D:10.8%,I:46.8%],F:2.4%,P:0.4%,G:0.3%,M:0.0%,n:1569

Tspe_10xpseudohap2-met BUSCOMP Results [1567 (97.09%) Complete BUSCOs; 139 (8.86%) BUSCOMP Seqs]:
        C:96.9%[S:86.1%,D:10.8%,I:46.8%],F:2.4%,P:0.4%,G:0.3%,M:0.0%,n:1569

Tspe_v0.1_canu-aug BUSCOMP Results [1345 (83.33%) Complete BUSCOs; 38 (2.42%) BUSCOMP Seqs]:
        C:99.4%[S:79.2%,D:20.3%,I:7.3%],F:0.4%,P:0.1%,G:0.1%,M:0.0%,n:1569

Tspe_v0.1_canu-met BUSCOMP Results [1520 (94.18%) Complete BUSCOs; 31 (1.98%) BUSCOMP Seqs]:
        C:99.4%[S:79.2%,D:20.3%,I:7.3%],F:0.4%,P:0.1%,G:0.1%,M:0.0%,n:1569

Tspe_v0.1_flye-aug BUSCOMP Results [1389 (86.06%) Complete BUSCOs; 43 (2.74%) BUSCOMP Seqs]:
        C:98.9%[S:88.0%,D:10.9%,I:34.4%],F:0.8%,P:0.2%,G:0.1%,M:0.0%,n:1569

Tspe_v0.1_flye-met BUSCOMP Results [1519 (94.11%) Complete BUSCOs; 41 (2.61%) BUSCOMP Seqs]:
        C:98.9%[S:88.0%,D:10.9%,I:34.4%],F:0.8%,P:0.2%,G:0.1%,M:0.0%,n:1569

Tspe_v0.1_necat-aug BUSCOMP Results [1381 (85.56%) Complete BUSCOs; 41 (2.61%) BUSCOMP Seqs]:
        C:99.9%[S:89.0%,D:10.8%,I:11.6%],F:0.0%,P:0.1%,G:0.0%,M:0.0%,n:1569

Tspe_v0.1_necat-met BUSCOMP Results [1533 (94.98%) Complete BUSCOs; 19 (1.21%) BUSCOMP Seqs]:
        C:99.9%[S:89.0%,D:10.8%,I:11.6%],F:0.0%,P:0.1%,G:0.0%,M:0.0%,n:1569

Tspe_v0.2-aug BUSCOMP Results [1368 (84.76%) Complete BUSCOs; 4 (0.25%) BUSCOMP Seqs]:
        C:99.9%[S:89.3%,D:10.6%,I:11.6%],F:0.0%,P:0.1%,G:0.0%,M:0.0%,n:1569

Tspe_v0.2-met BUSCOMP Results [1532 (94.92%) Complete BUSCOs; 0 (0.00%) BUSCOMP Seqs]:
        C:99.9%[S:89.3%,D:10.6%,I:11.6%],F:0.0%,P:0.1%,G:0.0%,M:0.0%,n:1569

Tspe_v0.3-aug BUSCOMP Results [1533 (94.98%) Complete BUSCOs; 81 (5.16%) BUSCOMP Seqs]:
        C:99.9%[S:89.3%,D:10.6%,I:33.5%],F:0.0%,P:0.1%,G:0.0%,M:0.0%,n:1569

Tspe_v0.3-met BUSCOMP Results [1590 (98.51%) Complete BUSCOs; 32 (2.04%) BUSCOMP Seqs]:
        C:99.9%[S:89.3%,D:10.6%,I:33.5%],F:0.0%,P:0.1%,G:0.0%,M:0.0%,n:1569

Tspe_v0.4-aug BUSCOMP Results [1555 (96.34%) Complete BUSCOs; 66 (4.21%) BUSCOMP Seqs]:
        C:99.9%[S:89.2%,D:10.6%,I:42.3%],F:0.0%,P:0.1%,G:0.0%,M:0.0%,n:1569

Tspe_v0.4-met BUSCOMP Results [1602 (99.26%) Complete BUSCOs; 9 (0.57%) BUSCOMP Seqs]:
        C:99.9%[S:89.2%,D:10.6%,I:42.3%],F:0.0%,P:0.1%,G:0.0%,M:0.0%,n:1569

Tspe_v0.5-aug BUSCOMP Results [1558 (96.53%) Complete BUSCOs; 11 (0.70%) BUSCOMP Seqs]:
        C:99.9%[S:89.2%,D:10.7%,I:42.3%],F:0.0%,P:0.1%,G:0.1%,M:0.0%,n:1569

Tspe_v0.5-met BUSCOMP Results [1603 (99.32%) Complete BUSCOs; 0 (0.00%) BUSCOMP Seqs]:
        C:99.9%[S:89.2%,D:10.7%,I:42.3%],F:0.0%,P:0.1%,G:0.1%,M:0.0%,n:1569

Tspe_v0.6-aug BUSCOMP Results [1532 (94.92%) Complete BUSCOs; 25 (1.59%) BUSCOMP Seqs]:
        C:99.9%[S:89.2%,D:10.6%,I:34.7%],F:0.0%,P:0.1%,G:0.1%,M:0.0%,n:1569

Tspe_v0.6-met BUSCOMP Results [1587 (98.33%) Complete BUSCOs; 11 (0.70%) BUSCOMP Seqs]:
        C:99.9%[S:89.2%,D:10.6%,I:34.7%],F:0.0%,P:0.1%,G:0.1%,M:0.0%,n:1569

Tspe_v0.7-aug BUSCOMP Results [1533 (94.98%) Complete BUSCOs; 2 (0.13%) BUSCOMP Seqs]:
        C:99.9%[S:89.2%,D:10.6%,I:34.7%],F:0.0%,P:0.1%,G:0.1%,M:0.0%,n:1569

Tspe_v0.7-met BUSCOMP Results [1587 (98.33%) Complete BUSCOs; 0 (0.00%) BUSCOMP Seqs]:
        C:99.9%[S:89.2%,D:10.6%,I:34.7%],F:0.0%,P:0.1%,G:0.1%,M:0.0%,n:1569

Tspe_v0.8-aug BUSCOMP Results [1457 (90.27%) Complete BUSCOs; 37 (2.36%) BUSCOMP Seqs]:
        C:97.4%[S:88.1%,D:9.3%,I:17.7%],F:2.0%,P:0.2%,G:0.4%,M:0.0%,n:1569

Tspe_v0.8-met BUSCOMP Results [1544 (95.66%) Complete BUSCOs; 26 (1.66%) BUSCOMP Seqs]:
        C:97.4%[S:88.1%,D:9.3%,I:17.7%],F:2.0%,P:0.2%,G:0.4%,M:0.0%,n:1569

Tspe_v0.9-aug BUSCOMP Results [1527 (94.61%) Complete BUSCOs; 83 (5.29%) BUSCOMP Seqs]:
        C:97.5%[S:88.1%,D:9.4%,I:52.7%],F:1.8%,P:0.3%,G:0.4%,M:0.0%,n:1569

Tspe_v0.9-met BUSCOMP Results [1579 (97.83%) Complete BUSCOs; 27 (1.72%) BUSCOMP Seqs]:
        C:97.5%[S:88.1%,D:9.4%,I:52.7%],F:1.8%,P:0.3%,G:0.4%,M:0.0%,n:1569

Tspe_v1-aug BUSCOMP Results [1526 (94.55%) Complete BUSCOs; 4 (0.25%) BUSCOMP Seqs]:
        C:97.5%[S:88.1%,D:9.4%,I:52.7%],F:1.8%,P:0.3%,G:0.4%,M:0.0%,n:1569

Tspe_v1-full BUSCOMP Results [1598 (99.01%) Complete BUSCOs; 0 (0.00%) BUSCOMP Seqs]:
        C:97.5%[S:90.4%,D:7.1%],F:1.8%,P:0.7%,G:0.0%,M:0.0%,n:1569

Tspe_v1-genome BUSCOMP Results [1596 (98.88%) Complete BUSCOs; 0 (0.00%) BUSCOMP Seqs]:
        C:97.5%[S:88.3%,D:9.2%],F:1.8%,P:0.3%,G:0.4%,M:0.0%,n:1569

Tspe_v1-met BUSCOMP Results [1579 (97.83%) Complete BUSCOs; 0 (0.00%) BUSCOMP Seqs]:
        C:97.5%[S:88.1%,D:9.4%,I:52.7%],F:1.8%,P:0.3%,G:0.4%,M:0.0%,n:1569

Tspe_v1-proteome BUSCOMP Results [1524 (94.42%) Complete BUSCOs; 0 (0.00%) BUSCOMP Seqs]:
        C:0.0%[S:0.0%,D:0.0%],F:0.0%,P:0.0%,G:0.0%,M:0.0%,n:1569

Tspe_v1-transcriptome BUSCOMP Results [1521 (94.24%) Complete BUSCOs; 0 (0.00%) BUSCOMP Seqs]:
        C:83.7%[S:75.3%,D:8.4%],F:1.1%,P:13.3%,G:0.6%,M:1.2%,n:1569

Tspe_v1.chr-aug BUSCOMP Results [1512 (93.68%) Complete BUSCOs; 3 (0.19%) BUSCOMP Seqs]:
        C:96.5%[S:87.4%,D:9.1%,I:52.2%],F:0.1%,P:1.3%,G:1.3%,M:0.8%,n:1569

Tspe_v1.chr-met BUSCOMP Results [1561 (96.72%) Complete BUSCOs; 0 (0.00%) BUSCOMP Seqs]:
        C:96.5%[S:87.4%,D:9.1%,I:52.2%],F:0.1%,P:1.3%,G:1.3%,M:0.8%,n:1569

Tspe_v1.gemoma BUSCOMP Results [1521 (94.24%) Complete BUSCOs; 417 (26.58%) BUSCOMP Seqs]:
        C:83.7%[S:74.3%,D:9.4%,I:43.9%],F:1.1%,P:13.3%,G:0.6%,M:1.2%,n:1569

Tspe_v1.gemoma.sprot.longest BUSCOMP Results [1521 (94.24%) Complete BUSCOs; 4 (0.25%) BUSCOMP Seqs]:
        C:83.7%[S:75.3%,D:8.4%,I:43.9%],F:1.1%,P:13.3%,G:0.6%,M:1.2%,n:1569

Tspe_v1.gemoma.sprot.renamed BUSCOMP Results [1521 (94.24%) Complete BUSCOs; 0 (0.00%) BUSCOMP Seqs]:
        C:83.7%[S:74.3%,D:9.4%,I:43.9%],F:1.1%,P:13.3%,G:0.6%,M:1.2%,n:1569
```

#### 4.2 BUSCOSeq Full Results Table

Full BUSCOMP results with ratings for each gene in every assembly and group have been compiled in `buscov5-full.N3L20ID0U.buscomp.tdt`:

#### 4.3 Genome Group BUSCOMP charts

### 5 BUSCO and BUSCOMP Comparisons

#### 5.1 BUSCO to BUSCOMP Rating Changes

Ratings changes from BUSCO to BUSCOMP (where `NULL` ratings indicate no BUSCOMP sequence):

| BUSCO | BUSCOMP | Tspe\_10xpseudohap2.aug | Tspe\_10xpseudohap2.met | Tspe\_v0.1\_canu.aug | Tspe\_v0.1\_canu.met | Tspe\_v0.1\_flye.aug | Tspe\_v0.1\_flye.met | Tspe\_v0.1\_necat.aug | Tspe\_v0.1\_necat.met | Tspe\_v0.2.aug | Tspe\_v0.2.met | Tspe\_v0.3.aug | Tspe\_v0.3.met | Tspe\_v0.4.aug | Tspe\_v0.4.met | Tspe\_v0.5.aug | Tspe\_v0.5.met | Tspe\_v0.6.aug | Tspe\_v0.6.met | Tspe\_v0.7.aug | Tspe\_v0.7.met | Tspe\_v0.8.aug | Tspe\_v0.8.met | Tspe\_v0.9.aug | Tspe\_v0.9.met | Tspe\_v1.aug | Tspe\_v1.met | Tspe\_v1.chr.aug | Tspe\_v1.chr.met | Tspe\_v1.gemoma | Tspe\_v1.gemoma.sprot.longest | Tspe\_v1.gemoma.sprot.renamed | TOTAL |
| --- | --- | --- | --- | --- | --- | --- | --- | --- | --- | --- | --- | --- | --- | --- | --- | --- | --- | --- | --- | --- | --- | --- | --- | --- | --- | --- | --- | --- | --- | --- | --- | --- | --- |
| Complete | Complete | 1244 | 1280 | 994 | 1117 | 1160 | 1258 | 1158 | 1274 | 1150 | 1278 | 1280 | 1319 | 1294 | 1330 | 1296 | 1330 | 1275 | 1319 | 1275 | 1319 | 1232 | 1280 | 1267 | 1301 | 1267 | 1301 | 1261 | 1291 | 1133 | 1149 | 1133 | 38565 |
| Complete | Duplicated | 84 | 58 | 174 | 123 | 111 | 77 | 113 | 76 | 106 | 75 | 88 | 58 | 79 | 44 | 81 | 45 | 88 | 60 | 91 | 60 | 83 | 74 | 71 | 46 | 70 | 46 | 68 | 44 | 30 | 30 | 30 | 2283 |
| Complete | Fragmented | 12 | 13 | 1 | 3 | 3 | 2 | 0 | 0 | 0 | 0 | 0 | 0 | 0 | 0 | 0 | 0 | 0 | 0 | 0 | 0 | 15 | 12 | 13 | 10 | 13 | 10 | 1 | 0 | 8 | 8 | 8 | 132 |
| Complete | Ghost | 0 | 1 | 1 | 1 | 1 | 1 | 0 | 0 | 0 | 0 | 0 | 0 | 0 | 0 | 1 | 1 | 1 | 1 | 1 | 1 | 2 | 3 | 2 | 2 | 2 | 2 | 2 | 2 | 1 | 1 | 1 | 31 |
| Complete | Missing | 0 | 0 | 0 | 0 | 0 | 0 | 0 | 0 | 0 | 0 | 0 | 0 | 0 | 0 | 0 | 0 | 0 | 0 | 0 | 0 | 0 | 0 | 0 | 0 | 0 | 0 | 0 | 1 | 0 | 0 | 0 | 1 |
| Complete | NULL | 34 | 35 | 30 | 31 | 36 | 33 | 31 | 34 | 31 | 34 | 34 | 33 | 33 | 34 | 33 | 34 | 35 | 35 | 35 | 35 | 35 | 34 | 34 | 35 | 34 | 35 | 33 | 34 | 34 | 38 | 34 | 1050 |
| Complete | Partial | 3 | 2 | 1 | 0 | 2 | 3 | 2 | 2 | 2 | 2 | 2 | 2 | 2 | 2 | 1 | 1 | 1 | 1 | 1 | 1 | 2 | 2 | 3 | 5 | 3 | 5 | 14 | 11 | 126 | 128 | 126 | 458 |
| Duplicated | Complete | 49 | 55 | 34 | 52 | 26 | 49 | 31 | 48 | 31 | 47 | 45 | 61 | 50 | 60 | 50 | 60 | 50 | 57 | 49 | 57 | 27 | 58 | 54 | 68 | 53 | 68 | 50 | 67 | 16 | 16 | 16 | 1454 |
| Duplicated | Duplicated | 85 | 111 | 99 | 181 | 44 | 87 | 37 | 89 | 39 | 86 | 74 | 106 | 86 | 122 | 85 | 122 | 73 | 105 | 72 | 105 | 51 | 70 | 73 | 101 | 74 | 101 | 72 | 99 | 118 | 102 | 118 | 2787 |
| Duplicated | Fragmented | 1 | 2 | 1 | 0 | 0 | 0 | 0 | 0 | 0 | 0 | 0 | 0 | 0 | 0 | 0 | 0 | 0 | 0 | 0 | 0 | 1 | 1 | 0 | 2 | 0 | 2 | 0 | 0 | 5 | 5 | 5 | 25 |
| Duplicated | Ghost | 0 | 0 | 0 | 0 | 0 | 0 | 0 | 0 | 0 | 0 | 0 | 0 | 0 | 0 | 0 | 0 | 0 | 0 | 0 | 0 | 0 | 0 | 0 | 0 | 0 | 0 | 1 | 1 | 0 | 0 | 0 | 2 |
| Duplicated | Missing | 0 | 0 | 0 | 0 | 0 | 0 | 0 | 0 | 0 | 0 | 0 | 0 | 0 | 0 | 0 | 0 | 0 | 0 | 0 | 0 | 0 | 0 | 0 | 0 | 0 | 0 | 1 | 0 | 0 | 0 | 0 | 1 |
| Duplicated | NULL | 9 | 9 | 10 | 11 | 6 | 9 | 9 | 10 | 9 | 10 | 10 | 11 | 11 | 10 | 11 | 10 | 9 | 9 | 9 | 9 | 9 | 10 | 9 | 9 | 9 | 9 | 9 | 9 | 10 | 6 | 10 | 290 |
| Duplicated | Partial | 1 | 1 | 0 | 1 | 0 | 0 | 0 | 0 | 0 | 0 | 0 | 0 | 0 | 0 | 0 | 0 | 0 | 0 | 0 | 0 | 0 | 0 | 1 | 0 | 1 | 0 | 0 | 2 | 40 | 38 | 40 | 125 |
| Fragmented | Complete | 22 | 11 | 65 | 47 | 62 | 44 | 74 | 52 | 82 | 53 | 29 | 17 | 22 | 8 | 21 | 7 | 31 | 18 | 31 | 18 | 43 | 30 | 22 | 10 | 23 | 10 | 24 | 9 | 14 | 14 | 14 | 927 |
| Fragmented | Duplicated | 1 | 1 | 16 | 9 | 9 | 4 | 5 | 5 | 7 | 5 | 1 | 2 | 1 | 1 | 1 | 1 | 3 | 2 | 1 | 2 | 4 | 2 | 2 | 0 | 2 | 0 | 2 | 0 | 0 | 0 | 0 | 89 |
| Fragmented | Fragmented | 11 | 17 | 0 | 3 | 2 | 6 | 0 | 0 | 0 | 0 | 0 | 0 | 0 | 0 | 0 | 0 | 0 | 0 | 0 | 0 | 3 | 14 | 5 | 14 | 5 | 14 | 0 | 1 | 5 | 5 | 5 | 110 |
| Fragmented | Ghost | 2 | 1 | 0 | 0 | 0 | 0 | 0 | 0 | 0 | 0 | 0 | 0 | 0 | 0 | 0 | 0 | 0 | 0 | 0 | 0 | 0 | 3 | 1 | 3 | 1 | 3 | 1 | 4 | 4 | 4 | 4 | 31 |
| Fragmented | Missing | 0 | 0 | 0 | 0 | 0 | 0 | 0 | 0 | 0 | 0 | 0 | 0 | 0 | 0 | 0 | 0 | 0 | 0 | 0 | 0 | 0 | 0 | 0 | 0 | 0 | 0 | 0 | 0 | 1 | 1 | 1 | 3 |
| Fragmented | NULL | 1 | 0 | 3 | 2 | 1 | 1 | 2 | 0 | 2 | 0 | 0 | 0 | 0 | 0 | 0 | 0 | 0 | 0 | 0 | 0 | 0 | 0 | 1 | 0 | 1 | 0 | 1 | 0 | 0 | 0 | 0 | 15 |
| Fragmented | Partial | 2 | 2 | 0 | 0 | 0 | 0 | 0 | 0 | 0 | 0 | 0 | 0 | 0 | 0 | 0 | 0 | 0 | 0 | 0 | 0 | 1 | 0 | 0 | 0 | 0 | 0 | 2 | 7 | 30 | 30 | 30 | 104 |
| Missing | Complete | 36 | 5 | 149 | 26 | 133 | 30 | 134 | 23 | 138 | 23 | 47 | 4 | 34 | 2 | 32 | 2 | 44 | 6 | 45 | 6 | 80 | 14 | 39 | 3 | 39 | 3 | 36 | 4 | 2 | 2 | 2 | 1143 |
| Missing | Duplicated | 0 | 0 | 29 | 5 | 7 | 3 | 15 | 0 | 14 | 0 | 3 | 0 | 1 | 0 | 1 | 0 | 3 | 0 | 3 | 0 | 8 | 0 | 1 | 0 | 1 | 0 | 1 | 0 | 0 | 0 | 0 | 95 |
| Missing | Fragmented | 13 | 5 | 5 | 1 | 8 | 5 | 0 | 0 | 0 | 0 | 0 | 0 | 0 | 0 | 0 | 0 | 0 | 0 | 0 | 0 | 12 | 4 | 11 | 3 | 11 | 3 | 0 | 0 | 0 | 0 | 0 | 81 |
| Missing | Ghost | 2 | 2 | 0 | 0 | 0 | 0 | 0 | 0 | 0 | 0 | 0 | 0 | 0 | 0 | 0 | 0 | 0 | 0 | 0 | 0 | 5 | 1 | 3 | 1 | 3 | 1 | 16 | 13 | 5 | 5 | 5 | 62 |
| Missing | Missing | 0 | 0 | 0 | 0 | 0 | 0 | 0 | 0 | 0 | 0 | 0 | 0 | 0 | 0 | 0 | 0 | 0 | 0 | 0 | 0 | 0 | 0 | 0 | 0 | 0 | 0 | 12 | 12 | 18 | 18 | 18 | 78 |
| Missing | NULL | 1 | 1 | 2 | 1 | 2 | 2 | 3 | 1 | 3 | 1 | 1 | 1 | 1 | 1 | 1 | 1 | 1 | 1 | 1 | 1 | 1 | 1 | 1 | 1 | 1 | 1 | 2 | 2 | 1 | 1 | 1 | 40 |
| Missing | Partial | 1 | 2 | 0 | 0 | 1 | 0 | 0 | 0 | 0 | 0 | 0 | 0 | 0 | 0 | 0 | 0 | 0 | 0 | 0 | 0 | 0 | 1 | 1 | 0 | 1 | 0 | 5 | 1 | 13 | 13 | 13 | 52 |

```
Tspe_10xpseudohap2-aug unique Complete genes: 0 BUSCO; 0 BUSCOMP
Tspe_10xpseudohap2-met unique Complete genes: 0 BUSCO; 0 BUSCOMP
Tspe_v0.1_canu-aug unique Complete genes: 0 BUSCO; 0 BUSCOMP
Tspe_v0.1_canu-met unique Complete genes: 0 BUSCO; 0 BUSCOMP
Tspe_v0.1_flye-aug unique Complete genes: 0 BUSCO; 0 BUSCOMP
Tspe_v0.1_flye-met unique Complete genes: 0 BUSCO; 0 BUSCOMP
Tspe_v0.1_necat-aug unique Complete genes: 0 BUSCO; 0 BUSCOMP
Tspe_v0.1_necat-met unique Complete genes: 0 BUSCO; 0 BUSCOMP
Tspe_v0.2-aug unique Complete genes: 0 BUSCO; 0 BUSCOMP
Tspe_v0.2-met unique Complete genes: 0 BUSCO; 0 BUSCOMP
Tspe_v0.3-aug unique Complete genes: 0 BUSCO; 0 BUSCOMP
Tspe_v0.3-met unique Complete genes: 0 BUSCO; 0 BUSCOMP
Tspe_v0.4-aug unique Complete genes: 0 BUSCO; 0 BUSCOMP
Tspe_v0.4-met unique Complete genes: 0 BUSCO; 0 BUSCOMP
Tspe_v0.5-aug unique Complete genes: 0 BUSCO; 0 BUSCOMP
Tspe_v0.5-met unique Complete genes: 0 BUSCO; 0 BUSCOMP
Tspe_v0.6-aug unique Complete genes: 0 BUSCO; 0 BUSCOMP
Tspe_v0.6-met unique Complete genes: 0 BUSCO; 0 BUSCOMP
Tspe_v0.7-aug unique Complete genes: 0 BUSCO; 0 BUSCOMP
Tspe_v0.7-met unique Complete genes: 0 BUSCO; 0 BUSCOMP
Tspe_v0.8-aug unique Complete genes: 0 BUSCO; 0 BUSCOMP
Tspe_v0.8-met unique Complete genes: 0 BUSCO; 0 BUSCOMP
Tspe_v0.9-aug unique Complete genes: 0 BUSCO; 0 BUSCOMP
Tspe_v0.9-met unique Complete genes: 0 BUSCO; 0 BUSCOMP
Tspe_v1-aug unique Complete genes: 0 BUSCO; 0 BUSCOMP
Tspe_v1-met unique Complete genes: 0 BUSCO; 0 BUSCOMP
Tspe_v1.chr-aug unique Complete genes: 0 BUSCO; 0 BUSCOMP
Tspe_v1.chr-met unique Complete genes: 0 BUSCO; 0 BUSCOMP
Tspe_v1.gemoma unique Complete genes: 0 BUSCO; 0 BUSCOMP
Tspe_v1.gemoma.prot unique Complete genes: 0 BUSCO; 0 BUSCOMP
Tspe_v1.gemoma.sprot.longest unique Complete genes: 0 BUSCO; 0 BUSCOMP
Tspe_v1.gemoma.sprot.longest.prot unique Complete genes: 0 BUSCO; 0 BUSCOMP
Tspe_v1.gemoma.sprot.renamed unique Complete genes: 0 BUSCO; 0 BUSCOMP
Tspe_v1.gemoma.sprot.renamed.prot unique Complete genes: 0 BUSCO; 0 BUSCOMP
Augustus unique Complete genes: 0 BUSCO; 0 BUSCOMP
MetaEuk unique Complete genes: 0 BUSCO; 0 BUSCOMP
Tspe_v1-proteome unique Complete genes: 0 BUSCO; 0 BUSCOMP
Tspe_v1-transcriptome unique Complete genes: 0 BUSCO; 0 BUSCOMP
```

#### 5.3 Ratings for Missing BUSCO genes

In addition to the unique ratings (above), it can be useful to know how genes `Missing` from one assembly/group are rated in the others. These plots are generated for each assembly/group in turn. The full BUSCO (`*.busco.tdt`) and BUSCOMP (`*.LnnIDxx.buscomp.tdt`) tables are reduced to the subset of genes that are missing in the assembly/group of interest, and then the summary ratings recalculated for that subset.

In each case, three plots are made (assuming both BUSCO and BUSCOMP data is available):

1. BUSCO ratings for missing BUSCO genes.
2. BUSCOMP ratings for missing BUSCO genes. As well as being more relaxed than pure BUSCO results, this will indicate when BUSCOMP has found a gene in the focal assembly/group where BUSCO did not.
3. BUSCOMP ratings for missing BUSCOMP genes. It is expected that assemblies will be much more similar in terms of BUSCOMP coverage.

#### 5.4 Missing Tspe\_10xpseudohap2-aug BUSCO genes

BUSCO ratings for `Missing` Tspe\_10xpseudohap2-aug BUSCO genes:

BUSCOMP ratings for `Missing` Tspe\_10xpseudohap2.aug BUSCO genes:

BUSCOMP ratings for `Missing` Tspe\_10xpseudohap2.aug BUSCOMP genes:

#### 5.5 Missing Tspe\_10xpseudohap2-met BUSCO genes

BUSCO ratings for `Missing` Tspe\_10xpseudohap2-met BUSCO genes:

BUSCOMP ratings for `Missing` Tspe\_10xpseudohap2.met BUSCO genes:

BUSCOMP ratings for `Missing` Tspe\_10xpseudohap2.met BUSCOMP genes:

#### 5.6 Missing Tspe\_v0.1\_canu-aug BUSCO genes

BUSCO ratings for `Missing` Tspe\_v0.1\_canu-aug BUSCO genes:

BUSCOMP ratings for `Missing` Tspe\_v0.1\_canu.aug BUSCO genes:

BUSCOMP ratings for `Missing` Tspe\_v0.1\_canu.aug BUSCOMP genes:

#### 5.7 Missing Tspe\_v0.1\_canu-met BUSCO genes

BUSCO ratings for `Missing` Tspe\_v0.1\_canu-met BUSCO genes:

BUSCOMP ratings for `Missing` Tspe\_v0.1\_canu.met BUSCO genes:

BUSCOMP ratings for `Missing` Tspe\_v0.1\_canu.met BUSCOMP genes:

#### 5.8 Missing Tspe\_v0.1\_flye-aug BUSCO genes

BUSCO ratings for `Missing` Tspe\_v0.1\_flye-aug BUSCO genes:

BUSCOMP ratings for `Missing` Tspe\_v0.1\_flye.aug BUSCO genes:

BUSCOMP ratings for `Missing` Tspe\_v0.1\_flye.aug BUSCOMP genes:

#### 5.9 Missing Tspe\_v0.1\_flye-met BUSCO genes

BUSCO ratings for `Missing` Tspe\_v0.1\_flye-met BUSCO genes:

BUSCOMP ratings for `Missing` Tspe\_v0.1\_flye.met BUSCO genes:

BUSCOMP ratings for `Missing` Tspe\_v0.1\_flye.met BUSCOMP genes:

#### 5.10 Missing Tspe\_v0.1\_necat-aug BUSCO genes

BUSCO ratings for `Missing` Tspe\_v0.1\_necat-aug BUSCO genes:

BUSCOMP ratings for `Missing` Tspe\_v0.1\_necat.aug BUSCO genes:

BUSCOMP ratings for `Missing` Tspe\_v0.1\_necat.aug BUSCOMP genes:

#### 5.11 Missing Tspe\_v0.1\_necat-met BUSCO genes

BUSCO ratings for `Missing` Tspe\_v0.1\_necat-met BUSCO genes:

BUSCOMP ratings for `Missing` Tspe\_v0.1\_necat.met BUSCO genes:

BUSCOMP ratings for `Missing` Tspe\_v0.1\_necat.met BUSCOMP genes:

#### 5.12 Missing Tspe\_v0.2-aug BUSCO genes

BUSCO ratings for `Missing` Tspe\_v0.2-aug BUSCO genes:

BUSCOMP ratings for `Missing` Tspe\_v0.2.aug BUSCO genes:

BUSCOMP ratings for `Missing` Tspe\_v0.2.aug BUSCOMP genes:

#### 5.13 Missing Tspe\_v0.2-met BUSCO genes

BUSCO ratings for `Missing` Tspe\_v0.2-met BUSCO genes:

BUSCOMP ratings for `Missing` Tspe\_v0.2.met BUSCO genes:

BUSCOMP ratings for `Missing` Tspe\_v0.2.met BUSCOMP genes:

#### 5.14 Missing Tspe\_v0.3-aug BUSCO genes

BUSCO ratings for `Missing` Tspe\_v0.3-aug BUSCO genes:

BUSCOMP ratings for `Missing` Tspe\_v0.3.aug BUSCO genes:

BUSCOMP ratings for `Missing` Tspe\_v0.3.aug BUSCOMP genes:

#### 5.15 Missing Tspe\_v0.3-met BUSCO genes

BUSCO ratings for `Missing` Tspe\_v0.3-met BUSCO genes:

BUSCOMP ratings for `Missing` Tspe\_v0.3.met BUSCO genes:

BUSCOMP ratings for `Missing` Tspe\_v0.3.met BUSCOMP genes:

#### 5.16 Missing Tspe\_v0.4-aug BUSCO genes

BUSCO ratings for `Missing` Tspe\_v0.4-aug BUSCO genes:

BUSCOMP ratings for `Missing` Tspe\_v0.4.aug BUSCO genes:

BUSCOMP ratings for `Missing` Tspe\_v0.4.aug BUSCOMP genes:

#### 5.17 Missing Tspe\_v0.4-met BUSCO genes

BUSCO ratings for `Missing` Tspe\_v0.4-met BUSCO genes:

BUSCOMP ratings for `Missing` Tspe\_v0.4.met BUSCO genes:

BUSCOMP ratings for `Missing` Tspe\_v0.4.met BUSCOMP genes:

#### 5.18 Missing Tspe\_v0.5-aug BUSCO genes

BUSCO ratings for `Missing` Tspe\_v0.5-aug BUSCO genes:

BUSCOMP ratings for `Missing` Tspe\_v0.5.aug BUSCO genes:

BUSCOMP ratings for `Missing` Tspe\_v0.5.aug BUSCOMP genes:

#### 5.19 Missing Tspe\_v0.5-met BUSCO genes

BUSCO ratings for `Missing` Tspe\_v0.5-met BUSCO genes:

BUSCOMP ratings for `Missing` Tspe\_v0.5.met BUSCO genes:

BUSCOMP ratings for `Missing` Tspe\_v0.5.met BUSCOMP genes:

#### 5.20 Missing Tspe\_v0.6-aug BUSCO genes

BUSCO ratings for `Missing` Tspe\_v0.6-aug BUSCO genes:

BUSCOMP ratings for `Missing` Tspe\_v0.6.aug BUSCO genes:

BUSCOMP ratings for `Missing` Tspe\_v0.6.aug BUSCOMP genes:

#### 5.21 Missing Tspe\_v0.6-met BUSCO genes

BUSCO ratings for `Missing` Tspe\_v0.6-met BUSCO genes:

BUSCOMP ratings for `Missing` Tspe\_v0.6.met BUSCO genes:

BUSCOMP ratings for `Missing` Tspe\_v0.6.met BUSCOMP genes:

#### 5.22 Missing Tspe\_v0.7-aug BUSCO genes

BUSCO ratings for `Missing` Tspe\_v0.7-aug BUSCO genes:

BUSCOMP ratings for `Missing` Tspe\_v0.7.aug BUSCO genes:

BUSCOMP ratings for `Missing` Tspe\_v0.7.aug BUSCOMP genes:

#### 5.23 Missing Tspe\_v0.7-met BUSCO genes

BUSCO ratings for `Missing` Tspe\_v0.7-met BUSCO genes:

BUSCOMP ratings for `Missing` Tspe\_v0.7.met BUSCO genes:

BUSCOMP ratings for `Missing` Tspe\_v0.7.met BUSCOMP genes:

#### 5.24 Missing Tspe\_v0.8-aug BUSCO genes

BUSCO ratings for `Missing` Tspe\_v0.8-aug BUSCO genes:

BUSCOMP ratings for `Missing` Tspe\_v0.8.aug BUSCO genes:

BUSCOMP ratings for `Missing` Tspe\_v0.8.aug BUSCOMP genes:

#### 5.25 Missing Tspe\_v0.8-met BUSCO genes

BUSCO ratings for `Missing` Tspe\_v0.8-met BUSCO genes:

BUSCOMP ratings for `Missing` Tspe\_v0.8.met BUSCO genes:

BUSCOMP ratings for `Missing` Tspe\_v0.8.met BUSCOMP genes:

#### 5.26 Missing Tspe\_v0.9-aug BUSCO genes

BUSCO ratings for `Missing` Tspe\_v0.9-aug BUSCO genes:

BUSCOMP ratings for `Missing` Tspe\_v0.9.aug BUSCO genes:

BUSCOMP ratings for `Missing` Tspe\_v0.9.aug BUSCOMP genes:

#### 5.27 Missing Tspe\_v0.9-met BUSCO genes

BUSCO ratings for `Missing` Tspe\_v0.9-met BUSCO genes:

BUSCOMP ratings for `Missing` Tspe\_v0.9.met BUSCO genes:

BUSCOMP ratings for `Missing` Tspe\_v0.9.met BUSCOMP genes:

#### 5.28 Missing Tspe\_v1-aug BUSCO genes

BUSCO ratings for `Missing` Tspe\_v1-aug BUSCO genes:

BUSCOMP ratings for `Missing` Tspe\_v1.aug BUSCO genes:

BUSCOMP ratings for `Missing` Tspe\_v1.aug BUSCOMP genes:

#### 5.29 Missing Tspe\_v1-met BUSCO genes

BUSCO ratings for `Missing` Tspe\_v1-met BUSCO genes:

BUSCOMP ratings for `Missing` Tspe\_v1.met BUSCO genes:

BUSCOMP ratings for `Missing` Tspe\_v1.met BUSCOMP genes:

#### 5.30 Missing Tspe\_v1.chr-aug BUSCO genes

BUSCO ratings for `Missing` Tspe\_v1.chr-aug BUSCO genes:

BUSCOMP ratings for `Missing` Tspe\_v1.chr.aug BUSCO genes:

BUSCOMP ratings for `Missing` Tspe\_v1.chr.aug BUSCOMP genes:

#### 5.31 Missing Augustus BUSCO genes

BUSCO ratings for `Missing` Augustus BUSCO genes:

BUSCOMP ratings for `Missing` Augustus BUSCO genes:

BUSCOMP ratings for `Missing` Augustus BUSCOMP genes:

#### 5.32 Missing Tspe\_v1.chr-met BUSCO genes

BUSCO ratings for `Missing` Tspe\_v1.chr-met BUSCO genes:

BUSCOMP ratings for `Missing` Tspe\_v1.chr.met BUSCO genes:

BUSCOMP ratings for `Missing` Tspe\_v1.chr.met BUSCOMP genes:

#### 5.33 Missing Tspe\_v1-genome BUSCO genes

BUSCO ratings for `Missing` Tspe\_v1-genome BUSCO genes:

#### 5.34 Missing MetaEuk BUSCO genes

BUSCO ratings for `Missing` MetaEuk BUSCO genes:

BUSCOMP ratings for `Missing` MetaEuk BUSCO genes:

BUSCOMP ratings for `Missing` MetaEuk BUSCOMP genes:

#### 5.35 Missing Tspe\_v1.gemoma BUSCO genes

BUSCO ratings for `Missing` Tspe\_v1.gemoma BUSCO genes:

BUSCOMP ratings for `Missing` Tspe\_v1.gemoma BUSCO genes:

BUSCOMP ratings for `Missing` Tspe\_v1.gemoma BUSCOMP genes:

#### 5.36 Missing Tspe\_v1.gemoma.prot BUSCO genes

BUSCO ratings for `Missing` Tspe\_v1.gemoma.prot BUSCO genes:

BUSCOMP ratings for `Missing` Tspe\_v1.gemoma.prot BUSCO genes:

#### 5.37 Missing Tspe\_v1.gemoma.sprot.longest BUSCO genes

BUSCO ratings for `Missing` Tspe\_v1.gemoma.sprot.longest BUSCO genes:

BUSCOMP ratings for `Missing` Tspe\_v1.gemoma.sprot.longest BUSCO genes:

BUSCOMP ratings for `Missing` Tspe\_v1.gemoma.sprot.longest BUSCOMP genes:

#### 5.38 Missing Tspe\_v1.gemoma.sprot.longest.prot BUSCO genes

BUSCO ratings for `Missing` Tspe\_v1.gemoma.sprot.longest.prot BUSCO genes:

BUSCOMP ratings for `Missing` Tspe\_v1.gemoma.sprot.longest.prot BUSCO genes:

#### 5.39 Missing Tspe\_v1.gemoma.sprot.renamed BUSCO genes

BUSCO ratings for `Missing` Tspe\_v1.gemoma.sprot.renamed BUSCO genes:

BUSCOMP ratings for `Missing` Tspe\_v1.gemoma.sprot.renamed BUSCO genes:

BUSCOMP ratings for `Missing` Tspe\_v1.gemoma.sprot.renamed BUSCOMP genes:

#### 5.40 Missing Tspe\_v1-transcriptome BUSCO genes

BUSCO ratings for `Missing` Tspe\_v1-transcriptome BUSCO genes:

#### 5.41 Missing Tspe\_v1.gemoma.sprot.renamed.prot BUSCO genes

BUSCO ratings for `Missing` Tspe\_v1.gemoma.sprot.renamed.prot BUSCO genes:

BUSCOMP ratings for `Missing` Tspe\_v1.gemoma.sprot.renamed.prot BUSCO genes:

#### 5.42 Missing Tspe\_v1-proteome BUSCO genes

BUSCO ratings for `Missing` Tspe\_v1-proteome BUSCO genes:

#### 5.43 Missing Tspe\_v1-full BUSCO genes

BUSCO ratings for `Missing` Tspe\_v1-full BUSCO genes:

#### 5.44 Missing BUSCOMP BUSCO genes

BUSCO ratings for `Missing` BUSCOMP BUSCO genes:

BUSCOMP ratings for `Missing` BUSCOMP BUSCO genes:

BUSCOMP ratings for `Missing` BUSCOMP BUSCOMP genes:

### 6 Appendix: BUSCOMP run details

```
BUSCOMP V0.12.1: run Mon Oct  4 21:50:25 2021
```

This analysis was run in:

```
/srv/scratch/ausplant/Telopea-Aug19/paper/2021-10-03.BUSCOMP/waratah-full
```

- Log file: `/srv/scratch/ausplant/Telopea-Aug19/paper/2021-10-03.BUSCOMP/waratah-full/buscov5-full.2.log`
- Commandline arguments: `runs=/srv/scratch/ausplant/Telopea-Aug19/annotation/2021-10-02.BUSCOv5/busco5metaeuk/run*,/srv/scratch/ausplant/Telopea-Aug19/annotation/2021-10-02.BUSCOv5/busco5augustus/run*,/srv/scratch/ausplant/Telopea-Aug19/annotation/2021-10-02.SAAGA/busco5/run*` `fastadir=/srv/scratch/z5239484/Telopea-Aug19/data/2021-04-05.TspeWorkflow/,../fasta/,../../../annotation/2021-10-02.SAAGA/` `basefile=buscov5-full` `forks=24` `backups=F` `log=buscov5-full.2.log` `newlog`
- Full Command List: `mafft=mafft` `clustalo=clustalo` `clustalw=clustalw2` `blast+path=` `iupath=/home/z3452659/bioware/iupred/iupred` `iuchdir=T` `minregion=5` `iucut=0.2` `rpath=R` `modpurge=T` `runs=/srv/scratch/ausplant/Telopea-Aug19/annotation/2021-10-02.BUSCOv5/busco5metaeuk/run*,/srv/scratch/ausplant/Telopea-Aug19/annotation/2021-10-02.BUSCOv5/busco5augustus/run*,/srv/scratch/ausplant/Telopea-Aug19/annotation/2021-10-02.SAAGA/busco5/run*` `fastadir=/srv/scratch/z5239484/Telopea-Aug19/data/2021-04-05.TspeWorkflow/,../fasta/,../../../annotation/2021-10-02.SAAGA/` `basefile=buscov5-full` `forks=24` `backups=F` `log=buscov5-full.2.log` `newlog` `genomesize=981953849`

#### 6.1 BUSCOMP errors

BUSCOMP returned no runtime errors.

#### 6.2 BUSCOMP warnings

See run log for further details:

```
#WARN   00:00:00    No Fasta file found for Tspe_v1.gemoma.prot
#WARN   00:00:00    No Fasta file found for Tspe_v1.gemoma.sprot.longest.prot
#WARN   00:00:00    No Fasta file found for Tspe_v1.gemoma.sprot.renamed.prot
#WARN   00:00:02    Using loaded stats data - might need to check for genome size consistency for NG50 and LG50.
```

---

**Report contents:**

- Run summary
- BUSCOMP summary
- Genome summary
- BUSCO Ratings
- BUSCO full results compilation
- BUSCOMP Sequence details and rating
- BUSCOMP re-rating of genomes
- BUSCOMP re-rating full results
- Unique BUSCO and BUSCOMP Complete genes
- Ratings for Missing BUSCO genes
- Appendix: BUSCOMP run details

---

Output generated by BUSCOMP v0.12.1 © 2019 Richard Edwards |
